# Altered neurodevelopment, CSF volume, and emotional reactivity following postnatal Zika virus infection in infant macaques

**DOI:** 10.64898/2026.03.27.714817

**Authors:** Nisarg Desai, Kaitlyn Love, Alex van Schoor, Sienna Freeman, Muskan Ali, Rebecca Richardson, Zsofia A Kovacs-Balint, Ruy Amaru Tobar Mosqueira, Rachel Lebovic, Jose Acevedo-Polo, Roza Vlasova, Martin Styner, Mar M Sanchez, Kathryn Moore, Nils Schoof, Patrick Whang, Vidisha Singh, Venkata-Viswanadh Edara, Mehul S. Suthar, Ann Chahroudi, Jessica Raper

**Affiliations:** Emory National Biomedical Research Center, Emory University, 954 Gatewood Rd NE, Atlanta GA 30329, USA; Department of Psychiatry, 101 Manning Drive, CB# 7160, University of North Carolina, Chapel Hill, NC 27599, USA; Department of Psychiatry and Behavioral Sciences, Emory University School of Medicine, Emory Brain Health Center, 12 Executive Park Dr NE #200, Atlanta, GA 30329, USA; Department of Pediatrics, Emory University School of Medicine, 2015 Uppergate Dr, Atlanta GA 30329, USA; Children’s Healthcare of Atlanta,1575 Northeast Expressway, Atlanta GA 30329, USA; Marcus Autism Center, Emory University School of Medicine, Department of Pediatrics and Children’s Healthcare of Atlanta, 1920 Briarcliff Rd NE, Atlanta, GA 30329, USA; Department of Biomedical Engineering, University of Basel, Hegenheimermattweg 167C, 4123 Allschwil, Switzerland

**Keywords:** Nonhuman primate, Sex differences, Neurobehavioral Assessment, Emotional Regulation, Visual Acuity

## Abstract

Although congenital Zika virus (ZIKV) syndrome is well-characterized, the neurodevelopmental consequences of postnatal infection are understudied. We used a rhesus macaque (RM) model to investigate the consequences of postnatal ZIKV infection on brain and behavior. Male and female infant RMs infected with ZIKV at 1 month of age were compared to sex-, age-, and rearing-matched controls: uninfected and treated with Poly-IC as an innate immune control. ZIKV-infected infants showed poor state control in early infancy and increased hostility during an acute stressor. They displayed sex-specific behavioral alterations during caregiver separation though attachment bonds were preserved. MRI at 3 months revealed larger total cerebrospinal fluid (CSF) volume, reduced visual processing region volumes, and sex-specific brain structural alterations: males exhibited amygdala hypertrophy, whereas females had volumetric reductions in temporal-limbic and temporal-auditory cortices. These findings indicate that postnatal ZIKV infection is associated with alterations in sensory, social and emotion-regulatory systems and in CSF volume, that may warrant long-term monitoring of exposed children.

## INTRODUCTION

Despite the discovery of Zika virus (ZIKV) in 1947, the first reports of neurological complications from ZIKV infection occurred in French Polynesia in 2013 ^1–3^. A few years later, an outbreak of ZIKV throughout Brazil and South America demonstrated a link between birth defects and fetal ZIKV exposure, termed congenital Zika syndrome (CZS) ^4,5^. It is estimated that 5-13% of infants with fetal ZIKV exposure present with microcephaly or other CZS-related birth defects, including cortical thinning, calcification of subcortical structures, impaired vision, and hearing loss ^6,7^. Despite the majority of infants with fetal ZIKV exposure appearing normal at birth, some infants can develop microcephaly or other neurodevelopmental problems after birth ^8,9^, suggesting adverse impacts on the brain during postnatal development. Although CZS is well-characterized, the neurodevelopmental consequences of postnatal infection have remained understudied.

The brain undergoes rapid growth from birth to two years of age in humans ^10–14^, creating a period of considerable vulnerability to insults. Studies have shown that children account for between 10-31% of ZIKV infections ^15–17^ with the virus transmitted through mosquito bites or breast milk ^18,19^. While most pediatric ZIKV infection reports appear to be mild (fever and rash) ^20^, neurological complications have been reported, including Guillian-Barre Syndrome, polyneuropathy, encephalitis, demyelinating diseases, cranial nerve abnormalities, and inflammatory diseases of the central nervous system (CNS) ^21–24^. Beyond the acute infection period, there have been few longitudinal studies of neurodevelopment following postnatal ZIKV infection. One prospective study of 60 children with ZIKV infection between 1 and 12 months of age found that 15% had adverse neurologic, hearing or eye examinations at 20-30 months of age ^17^. These data suggest that ZIKV neurotropism can lead to adverse neurodevelopmental consequences for the vulnerable developing brain, but the full extent of this impact is still largely unknown. Studies with translational nonhuman primate (NHP) models of similar immune and brain development as humans provide a critical advantage to address these questions.

We have developed a NHP model to investigate the developmental consequences of postnatal ZIKV infection during infancy. Our previous preliminary findings demonstrated that ZIKV disseminates to the central and peripheral nervous systems in infant female rhesus macaques (RMs) infected at 5 weeks of age and lead to altered brain development with corresponding behavioral changes at 6 and 12 months of age ^25,26^. The current study aimed to extend these findings using a larger cohort of both male and female infant RMs infected with ZIKV compared to age-, sex-, and rearing-matched uninfected and innate immune stimulated controls. Overall, our findings suggest that ZIKV infection during infancy led to significant alterations in emotional processing, including poor state control detected shortly after infection, increased irritability at 5 months of age, but blunted distress at ∼8 months of age. These changes in emotional behavior were accompanied by alterations in neurodevelopment as detected by MRI at 3 months of age with a surprising sex dependent vulnerability to ZIKV infection, such that male RMs exhibited more brain structural changes as compared to females. Importantly, innate immune stimulation alone did not produce changes in behavior and neuronal development, suggesting that results are specific to ZIKV infection.

## RESULTS

### Viral Dynamics and Response to Poly-IC

Twelve infant RMs (6 females and 6 males) were infected subcutaneously over the cranial dorsum with 10^5^ plaque forming units of ZIKV (strain PRVABC59) at a median of 35 days of life (range: 33-39 days; approximately 5 weeks of age; Fig 1A). ZIKV RNA peaked in plasma at 2-3 days post-infection (dpi), with concentrations ranging from 10^5^ to 10^8^ copies of ZIKV RNA per milliliter (Fig 1B). Similar to previous studies in adult and infant RMs ^25,27–30^, ZIKV RNA was rapidly cleared from plasma, reaching undetectable concentrations between 5 and 21 dpi, ten animals were cleared by 14 dpi, but two females were still detectable at 14 dpi. Viral burden did not differ credibly between the sexes (male versus female difference in log-AUC = −0.93, 89% HDI [−2.51, 0.69], pd = 0.83; fold-change 0.40 [0.08, 1.99]), as shown by the area under the curve for plasma viral RNA (Fig 1C, Supplementary Table S13). As a control for the innate immune activation that follows ZIKV infection, six additional infant RMs (3 females and 3 males) were treated with Poly-IC (PIC), a synthetic double stranded RNA that stimulates Toll-like receptor 3, at a median age of 38 days of life (range: 35-39 days). Six infant RMs (3 females and 3 males) served as uninfected, untreated controls (UIC). Poly-IC treated infant RMs showed credibly elevated IFN-gamma (Poly-IC minus control difference in log10 fold change = 1.34, 89% HDI [0.10, 2.53]; probability of increase 0.95), IL-1Ra (0.58, [0.12, 1.03]; 0.97), and IP-10 (0.60, [0.17, 1.02]; 0.98) together with a probable increase in IFN-alpha2a (0.63, [−0.16, 1.44]; 0.90). These expected increases in systemic cytokines and chemokines confirm robust innate immune activation in the Poly-IC group relative to uninfected controls (Fig 1D).

**Figure 1:**
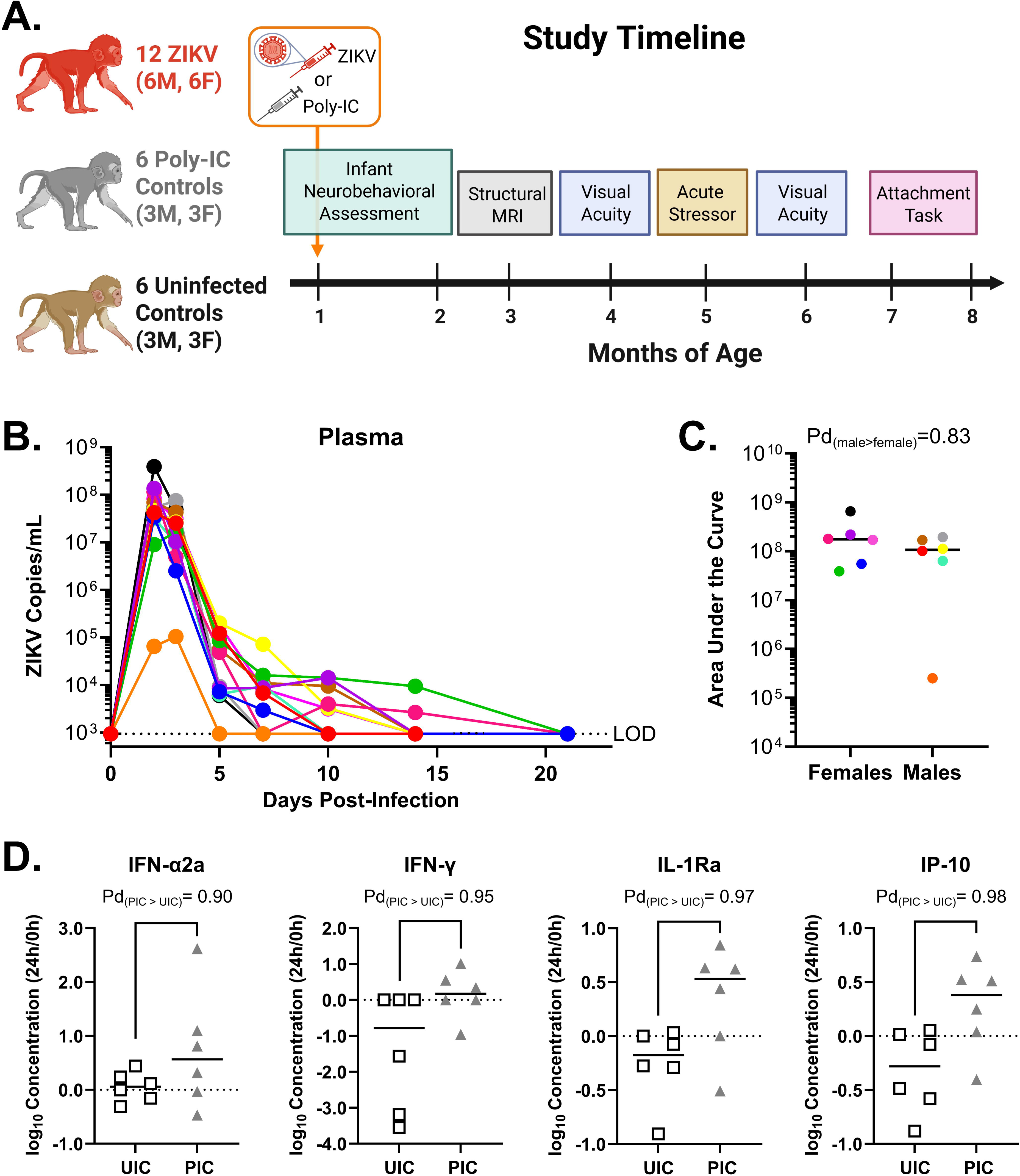
Viral dynamics of ZIKV infection and increased inflammatory response to innate immune activation in infant RMs. Study design and timeline. (A) Rhesus macaques received subcutaneous injections with 10^5^ plaque forming units (PFU) ZIKV strain PRVABC59 or Poly-IC. Sex and age matched uninjected animals were used as uninfected controls. All animals underwent a battery of assessments from 2 weeks to ∼8 months of life. (B). ZIKV RNA load as measured by qRT-PCR in plasma with limit of detection (LOD) 939 copies/ml (C) and area under the curve, each colored circle and line represents a different ZIKV-infected animal. (D) Cytokine and chemokine response to Poly-IC administration (PIC; grey triangles) compared to uninfected controls (UIC; open squares). Probability of direction (Pd) derived from the Bayesian Gaussian regression model for AUC and log10 fold-change increase in plasma levels. Figure partially made with biorender.com and GraphPad Prism. All Rights Reserved for photos copyright © Emory National Biomedical Research Center, Emory University.

### Neurobehavioral Assessment

To determine the impact of postnatal ZIKV infection on development, we analyzed four key domains from the Infant Neurobehavioral Assessment Scale (INAS) collected weekly from 2 to 9 weeks of age ^31,32^. Bayesian cumulative ordinal regression models were used to disentangle the effects of ZIKV infection from non-viral innate immune stimulation (PIC) and normal developmental progression. Figure 2 shows the model estimates for the contrasts performed with samples from the posterior. Because the models include a treatment-by-age interaction, each curve is the estimated group difference as a continuous function of age, and its slope reflects how that difference changes with age (a steeper slope indicates a more rapidly growing difference) rather than a change at any single time point. Interpretation should therefore focus on whether and when a difference becomes credible (its interval excludes zero) and on the overall trajectory across age, particularly the emergence and persistence of differences after challenge; per-week estimates from a complementary model with age modelled as a factor are provided in Supplementary Table S12.

**Figure 2:**
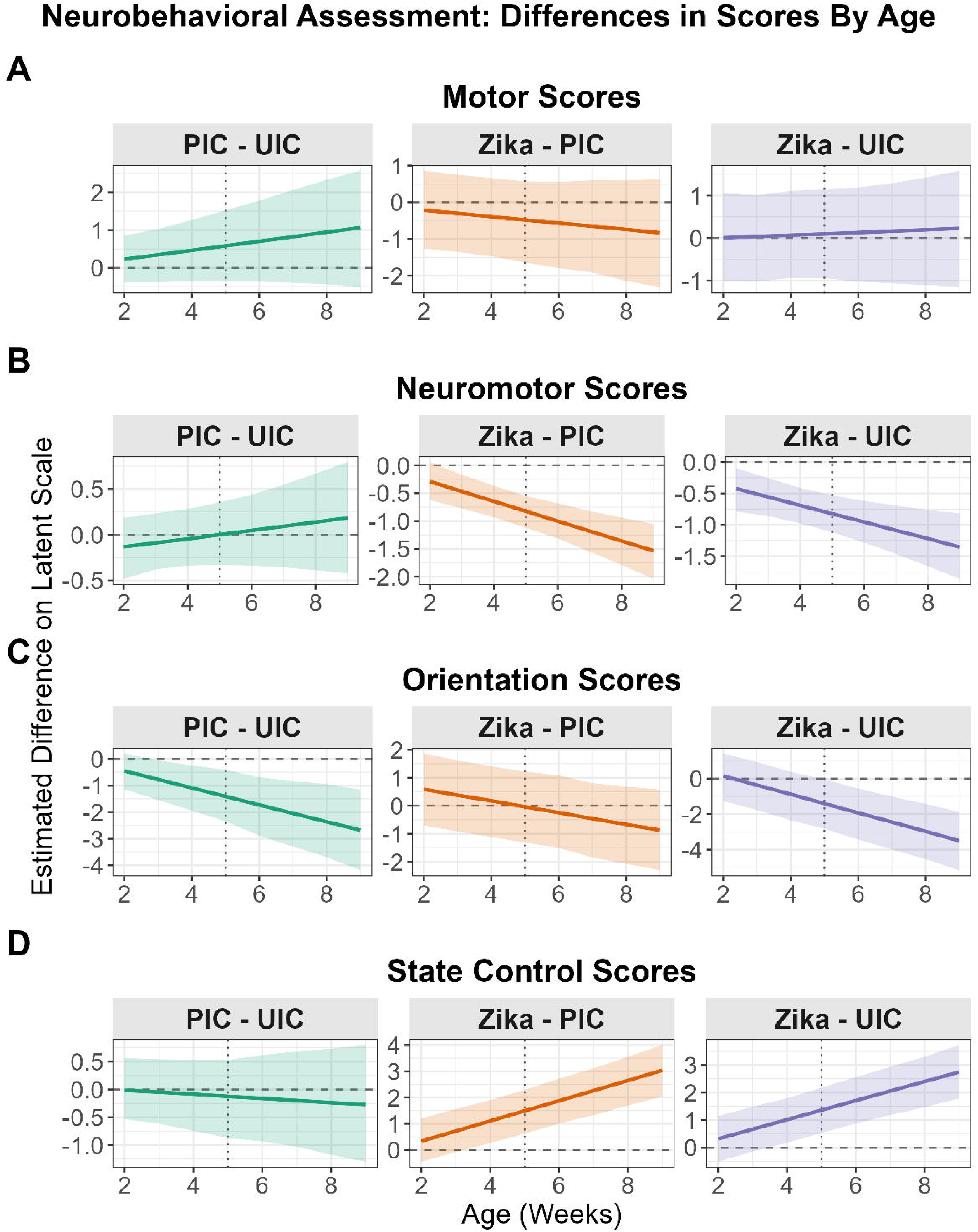
ZIKV infection and innate immune activation alter neurobehavioral scores in infant RMs. Bayesian model posterior estimates of treatment contrasts on neurobehavioral assessment scores over age. Each horizontal panel displays the estimated score difference on the latent logit scale for a specific assessment (A) Motor Scores, (B) Neuromotor Scores, (C) Orientation Scores, and (D) State Control Scores. Each panel within it displays treatment group comparison: Poly-IC (PIC) versus Uninfected Control (UIC) in green, ZIKV versus PIC in orange, and ZIKV versus UIC in purple (n = 12 ZIKV, 6 UIC and 6 PIC). Solid lines represent the posterior median difference, and the surrounding shaded ribbons indicate the 89% Highest Posterior Density Interval (HDI). The horizontal dashed line at zero indicates no difference between groups; credible differences are suggested where an 89% HDI does not overlap with zero. The vertical dotted line in each panel indicates the timing of ZIKV challenge and Poly-IC administration (a median of 35 days of life; approximately 5 weeks of age); differences to the left of this line precede the challenge.

First, we examined motor maturity and neuromotor reflexes. Motor maturity scores indicated similar gross motor development across all three groups (all 89% Highest Posterior Density Intervals, HDIs, crossed zero; Fig 2A). However, neuromotor scores, which assess sensory responsiveness and neurological reflexes, revealed a progressive deficit in the ZIKV-infected infants. At baseline prior to infection, the ZIKV group displayed lower scores compared to the UIC group (median estimate = −0.42, 89% HDI = [−0.77, −0.09]; Fig 2B). This disparity grew post-infection to −0.83 at 5 weeks (89% HDI = [−1.11, −0.52]) and peak magnitude of −1.35 (89% HDI = [−1.87, −0.81]) at 9 weeks of age. A similar pattern was observed when comparing ZIKV-infected infants to the PIC group, with a credible deficit emerging at 3 weeks (median estimate = −0.47, 89% HDI = [−0.77, −0.16]) and becoming progressively more severe thereafter (Fig 2B). There were no credible differences between the PIC and UIC control groups at any time point for neuromotor scores, strengthening the conclusion that the observed deficit in the ZIKV group is a robust effect, even with the small differences present before infection. A sensitivity analysis estimating each week separately (age as a factor; Supplementary Table S12) confirmed that a small neuromotor difference is already credible before challenge (ZIKV vs. PIC at 3 weeks = −0.70, 89% HDI [−1.23, −0.21]), so we interpret this early difference as pre-existing individual variation rather than a consequence of infection.

To assess higher-order processing, we examined orientation scores, which reflect attention and sensory awareness toward visual and auditory cues. Orientation abilities were equivalent across all groups at baseline and at the time of infection. Following infection, however, a credible deficit rapidly emerged in ZIKV-infected infants. Orientation scores dropped below UIC at 5 weeks of age (median estimate = −1.38, 89% HDI = [−2.80, −0.13]) and progressively worsened through 9 weeks of age (median estimate = −3.45, 89% HDI = [−5.13, −1.99]; Fig 2C). Notably, the PIC group showed a similar trajectory of decline when compared to the UIC group, suggesting that the disruption of developing attentional systems may be driven by inflammatory mechanisms common to both ZIKV infection and non-viral innate immune activation. Accordingly, scores for the ZIKV and PIC groups were similar across all timepoints (Fig 2C).

Finally, we evaluated state control ratings to determine if ZIKV infection altered emotional regulation. Higher scores on the state control domain indicate greater emotional reactivity (i.e. more irritable, more distressed and less consolable). Unlike the orientation domain where PIC animals mirrored ZIKV trends, poor state control scores were specific to ZIKV infection. All groups scored similarly at baseline (2-3 weeks of age). A credible effect emerged at 4 weeks of age, the weekly assessment preceding the ZIKV challenge (ZIKV vs. UIC median estimate = 1.00, 89% HDI = [0.14, 1.77]; ZIKV vs. PIC median estimate = 1.10, 89% HDI = [0.30, 1.96]; Fig 2D). However, because the primary model represents the treatment effect as a linear function of age, we confirmed the timing of onset with a sensitivity analysis that estimated each week separately (age modelled as a factor; Supplementary Table S12). In that analysis the ZIKV-specific increase, credible relative to both UIC and PIC, first emerged at the post-infection assessment (approximately 5 weeks of age), whereas the apparent 4-week signal in the primary model reflected a transient, non-specific reduction in the uninfected control group (ZIKV vs. PIC at 4 weeks = 0.19, 89% HDI [−0.87, 1.28]; PIC vs. UIC = 1.75, [0.51, 3.00]) rather than a ZIKV-specific effect. The differences in state control scores intensified weekly, culminating in a large effect by 9 weeks of age (ZIKV vs UIC median estimate = 2.75, 89% HDI = [1.84, 3.77]; ZIKV vs PIC mean estimate = 3.03, 89% HDI = [2.06, 4.04]). Crucially, the PIC group did not differ from UIC at any point, demonstrating that the observed persistent worsening of emotional regulation is a specific consequence of ZIKV infection rather than a generalized response to innate immune activation.

### Visual Attention Task

To determine whether the decreased INAS orientation scores observed in ZIKV-infected infant RMs were indicative of a fixed visual impairment, we used eye tracking to examine visual acuity with high, medium, and low contrast stimuli at 4 and 6 months of age. Initial pairwise comparisons revealed only minor differences between the UIC and PIC groups, and because ZIKV did not differ from either control group individually (Supplementary Table S9), these treatments were merged into a combined group (hereafter “control”).

Despite the deficit in orientation observed in early infancy, ZIKV-infected infants exhibited preserved visual acuity compared to the control group at 4 and 6 months of age. Pairwise contrasts comparing the ZIKV and control groups across specific ages (4 and 6 months) and contrast levels (high, medium, low) consistently yielded differences with 89% credible intervals spanning zero (Fig 3, supplementary materials Tables S2 & S3). These results indicate that ZIKV infection only led to temporary early deficits in visual orientation, without long-term loss of visual acuity.

**Figure 3:**
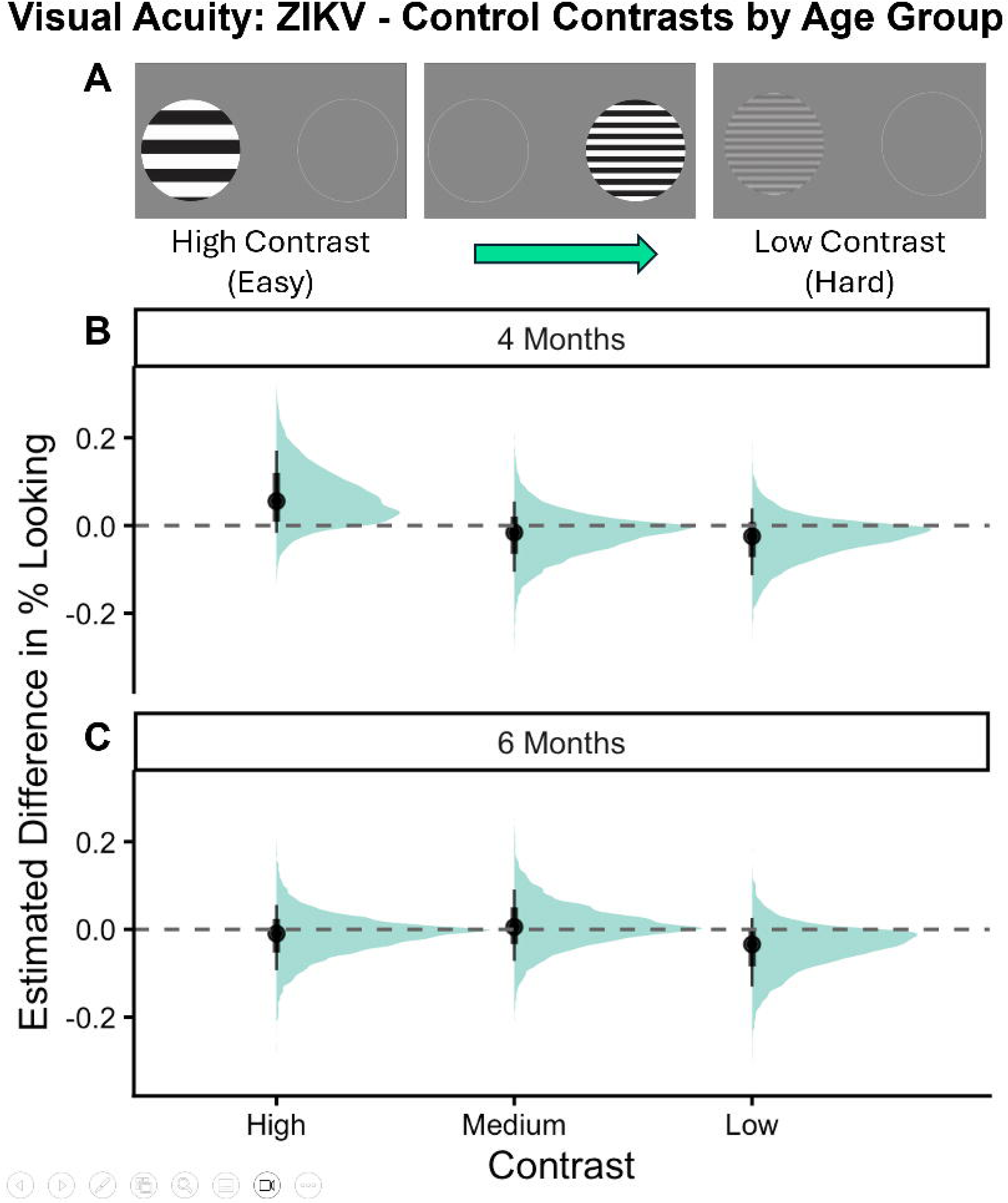
Visual acuity did not differ between ZIKV and control infant RMs. Posterior distributions of the ‘ZIKV - Control’ group contrast across different ages and stimulus contrast levels, n = 12 ZIKV and 12 control (6 uninfected + 6 Poly-IC). The plot displays the estimated difference in % Looking between the ZIKV and control experimental groups, derived from the Bayesian beta regression model. (A) Top panel shows the three contrast conditions in the eye tracking apparatus. The other two panels (B), (C) represent a distinct age group. Within each panel, the posterior distribution for the contrast is visualized at different stimulus contrast levels along the x-axis. The half-eye plots show the full probability density of the estimated difference. The black horizontal lines within each plot indicate the 89% (wider line) and 66% (narrower, thicker line) credible intervals, with the posterior median marked by a black dot. The horizontal dashed line at y=0 represents no difference; a contrast can be considered statistically meaningful where its credible interval does not overlap this line. All Rights Reserved for photos copyright © Emory National Biomedical Research Center, Emory University.

### Acute Stress Assessment

To determine whether the ZIKV-infected RMs poor state control on the INAS was an early marker of a persistent deficit in emotional regulation, we used the human intruder (HI) paradigm as an acute stressor at 5 months of age. The HI paradigm is based on a task used for assessing dispositional anxiety and behavioral inhibition in children ^33^ and robustly quantifies behavioral reactivity in RMs ^25,30,34^.

First, we confirmed that the paradigm elicited the expected behavioral repertoire across the cohort. Regardless of infection status, all RM infants exhibited the species-typical response toward the different levels of threat conditions of the HI paradigm (see supplementary materials for details). These consistent responses confirm that the paradigm effectively engaged the infants’ defensive circuits.

While the overall pattern of response was preserved, ZIKV-infected infant RMs showed credible elevations in hostile behaviors during the less salient threat conditions, differentiating them from controls in both the Alone (ZIKV-Control median estimate = 1.46; 89% HDI: [0.63, 2.26]) and Profile (ZIKV - Control median estimate = 1.54; 89% HDI: [0.66, 2.39]) conditions (Fig 4a). No ZKIV - Control differences were detected for freezing duration, anxious behaviors, self-directed behaviors, affiliative vocals, screams, or fearful behaviors (general sex effects independent of group are detailed in supplementary materials). Beyond the group-level difference in hostile behaviors across both sexes, we observed a sex-specific deficit in communicative defense. Threat barks were scored as a distinct threat vocalization in this assessment, separate from the hostile-behavior composite (Supplementary Table S1). While ZIKV-infected infant RMs were more hostile overall, when comparing ZIKV vs. controls within sexes (ZIKV - Control contrasts), ZIKV-infected females produced credibly fewer threat barks compared to controls across conditions overall (Females, ZIKV - Control median estimate = −1.05; 89% HDI: [−1.78, −0.32]). When stratifying these contrasts by condition for a more fine-grained insight (ZIKV - Control contrasts by sex and condition), this reduction was specific to the social threat phases, appearing in both the Profile (Female ZIKV - Control median estimate = −1.32; 89% HDI: [−2.28, −0.37]) and Stare (Female ZIKV - Control median estimate = −1.17; 89% HDI: [−2.07, −0.19]) conditions, whereas males showed no differences compared to controls (Profile condition, Male ZIKV - Control median estimate = −0.50; 89% HDI: [−1.57, 0.51]; Stare condition, Male ZIKV - Control median estimate = −0.30; 89% HDI: [−1.34, 0.73]; Fig 4b). These effects held when the two control groups were analyzed separately: the elevated hostility in the Alone (ZIKV vs. UIC median estimate = 1.64, 89% HDI: [0.71, 2.50]; ZIKV vs. PIC = 2.05, [0.84, 3.20]) and Profile (ZIKV vs. UIC = 1.65, [0.64, 2.61]; ZIKV vs. PIC = 1.39, [0.25, 2.43]) conditions, and the reduced threat barks in females (ZIKV vs. UIC = −1.07, [−1.84, −0.28]; ZIKV vs. PIC = −1.17, [−2.04, −0.22]), each remained credible relative to both the naïve (UIC) and immune-stimulated (PIC) control (Supplementary Table S7).

**Figure 4.**
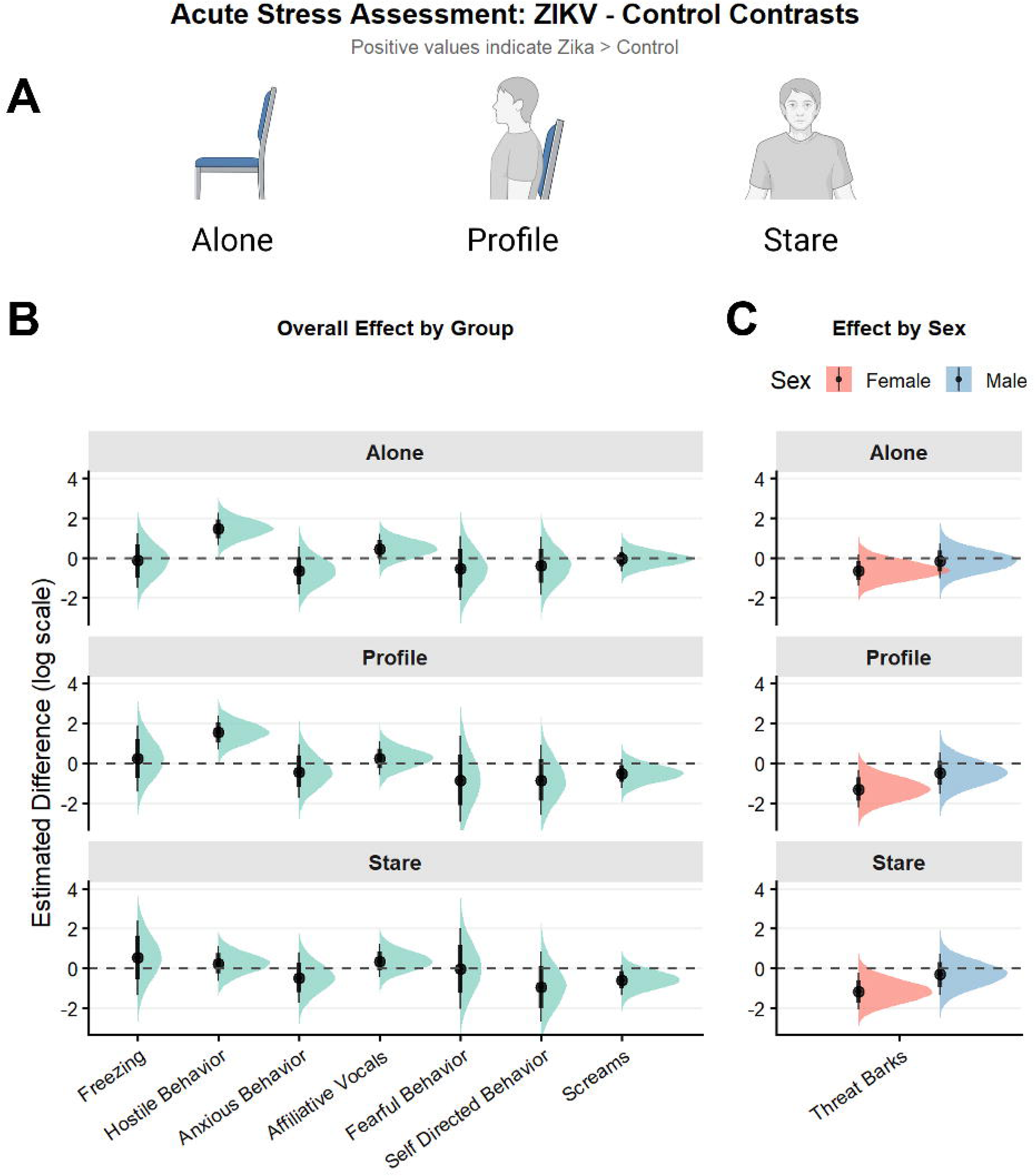
Increased hostility among ZIKV-infected infant RMs. (A) Schematic showing the three conditions during the acute stress assessment: no intruder in the Alone, the intruder facing sideways in the Profile and intruder facing the infant the Stare condition. (B) Posterior distributions of the overall ‘ZIKV - Control’ contrast across multiple behaviors during the acute stress assessment for both sexes (n = 12 ZIKV and 12 control (6 uninfected + 6 Poly-IC)) and (C) stratified by sex for threat barks due to interaction (n = 6 ZIKV and 6 control (3 uninfected + 3 Poly-IC) per sex). This plot displays the estimated difference on the log scale between the ZIKV and control groups for the tested behavioral outcomes. Estimates are derived from separate Bayesian negative binomial regression models for each behavior. Three panels separate the different conditions: Alone, Profile, and Stare. Each half-eye plot illustrates the full probability density of the estimated difference. The internal black lines denote the 89% (wider) and 66% (narrower, thicker) credible intervals, with the posterior median marked by a black dot. The horizontal dashed line at y=0 represents no difference between the groups; a contrast is considered meaningful when its credible interval does not overlap this line. Schematic made with biorender.com. All Rights Reserved for photos copyright © Emory National Biomedical Research Center, Emory University.

### Attachment Assessment

Early attachment to a primary caregiver can impact social and emotional development in later life ^35–37^. To evaluate whether poor state control and changes in emotional reactivity seen after ZIKV infection were due to alterations in early attachment, we examined the bond to a primary caregiver using a two-choice attachment task between 7 to 8 months of age. Similar to controls, ZIKV-infected RM infants exhibited a strong attachment to their primary caregiver, as shown by the Index of Preference (females: median estimate = −0.246, 89% HDI: [−1.04, 0.504]; males: median estimate = −0.576, 89% HDI: [−1.41, 0.342]; Fig 5, Table S4). There were also no differences seen in attention-seeking or affiliative behaviors. These results suggest that changes in state control and behavioral reactivity are not due to disrupted social attachment.

**Figure 5.**
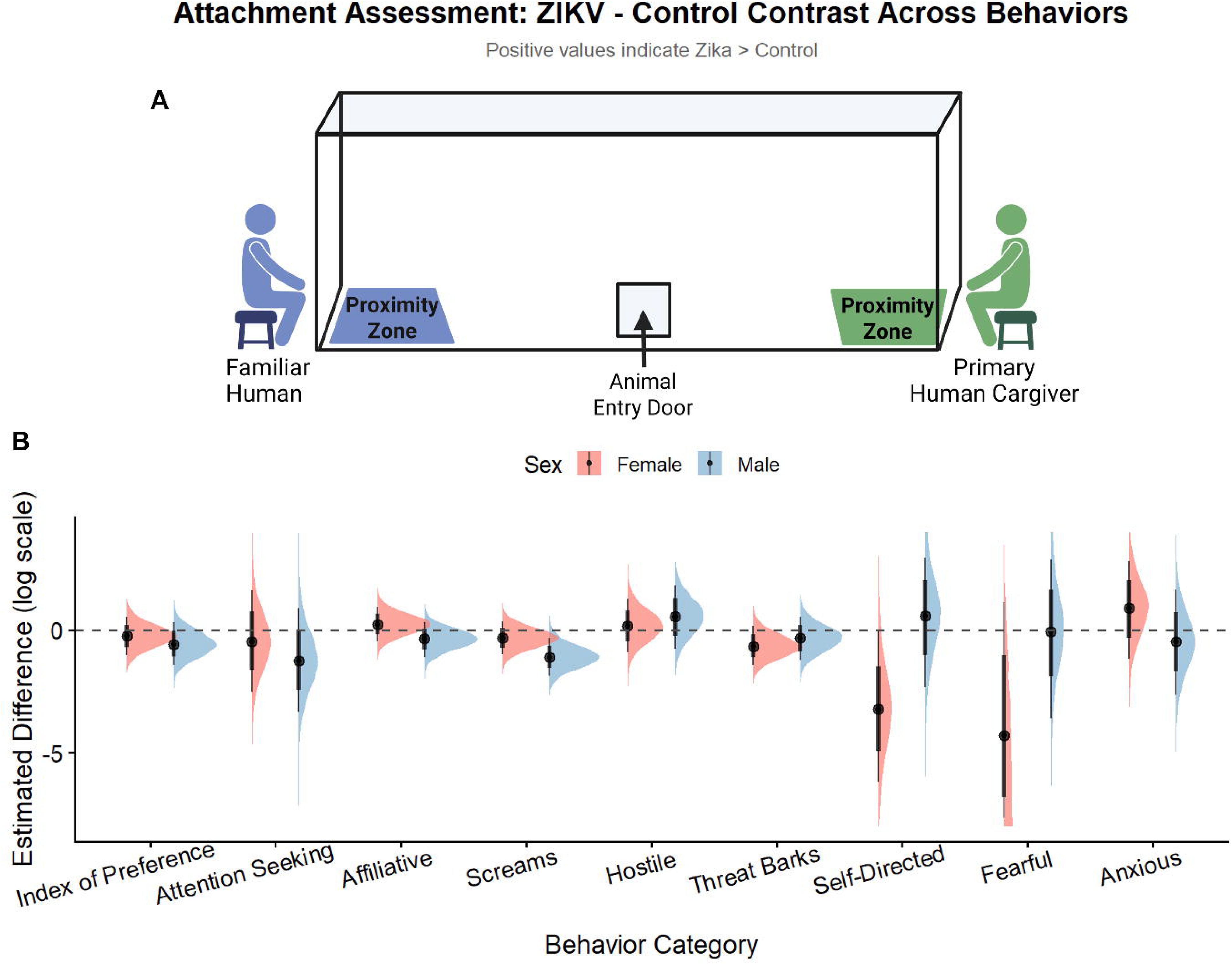
Attachment is not impacted by ZIKV infection during infancy. (A) Schematic showing the setup for the attachment task: the infant enters the cage from the middle and is free to choose to spend time close to either caregiver.(B) Sex-specific posterior distributions of the ‘ZIKV - Control’ contrast across multiple behaviors during the attachment task (n = 6 ZIKV and 6 control (3 uninfected + 3 Poly-IC) per sex). This plot displays the estimated difference on the log scale between control and ZIKV groups for the tested behavioral outcomes. Estimates are derived from separate Bayesian beta (for index of preference) or negative binomial (other behaviors) regression models for each behavior. For each behavior on the x-axis, the contrast is visualized side-by-side for females and males, distinguished by color. Each half-eye plot illustrates the full probability density of the estimated difference. The internal black lines denote the 89% (wider) and 66% (narrower, thicker) credible intervals, with the posterior median marked by a black dot. The horizontal dashed line at y=0 represents no difference between the groups; a contrast is considered meaningful when its credible interval does not overlap this line. Schematic made with biorender.com. All Rights Reserved for photos copyright © Emory National Biomedical Research Center, Emory University.

The two-choice attachment task can invoke a suite of separation responses because the infants can see their primary caregiver, but cannot reach them. Despite the preserved attachment preference, ZIKV-infected infants displayed a blunted distress response during the task compared to controls, including distinct sex-specific alterations in behavior. ZIKV-infected males exhibited fewer scream vocalizations than control males (ZIKV - Control median estimate = −1.10, 89% HDI: [−1.90, −0.36]; Fig 5), suggesting a blunted separation distress response. ZIKV-infected and control females emitted a similar number of screams (median estimate = −0.325, 89% HDI: [−0.981, 0.371]; Fig 5). In contrast, ZIKV-infected and control females did not differ credibly in self-directed behaviors (e.g., self-grooming; median estimate = −0.20, 89% HDI: [−1.54, 1.22]; Fig 5), and males showed no credible difference in self-directed behavior (median estimate = 0.32, 89% HDI: [−1.17, 2.02]). Hostile behaviors alone showed no credible difference in either sex (females: median estimate = 0.17, 89% HDI: [−0.92, 1.25]; males: median estimate = 0.55, 89% HDI: [−0.77, 1.83]; Fig 5). Threat barks showed a female-biased reduction concordant with the acute-stress finding (females: median estimate = −0.66, 89% HDI: [−1.44, 0.15]; rate ratio 0.51; probability of direction 0.90; males: median estimate = −0.33, 89% HDI: [−1.22, 0.57]), although in this smaller assessment it did not reach the 89% credibility threshold. No ZIKV - Control differences with 89% HDIs excluding zero were detected for fearful behaviors, or anxious behaviors. All analyzed behaviors are summarized in the supplementary materials (Table S5).

### Neuroimaging Assessment

Previous studies demonstrate that ZIKV neuroinvasion significantly altered postnatal brain development ^25,26,30,38^. This prior research was performed with small sample sizes (n=2-3 per group). We therefore conducted structural MRI scans at 3 months of age to further assess the evidence for structural changes after ZIKV infection during infancy in this new larger cohort. We analyzed volumes of brain regions important for emotional and sensory processing using Bayesian linear regression, comparing ZIKV-infected infants to controls (combined UIC and PIC) and their interaction with sex. The imaging analyses included 24 scans acquired at 3 months of age: 12 ZIKV-infected infants (6 female, 6 male) and 12 controls (6 female, 6 male, comprising 3 uninfected and 3 Poly-IC-treated animals of each sex). We combined UIC and PIC groups into a single control group as there were no differences in their total brain volume (TBV) (t = 0.28, df = 6.90, p = 0.79) and total intracranial volume (ICV) (t = 0.44, df = 5.56, p = 0.68). In addition, an analysis retaining all three groups (UIC, PIC, and ZIKV) found no credible differences between UIC and PIC in any regional volume, and confirmed that the principal ZIKV effects were preserved when each control group was compared with ZIKV separately (Supplementary Table S10). Initial analyses revealed that ZIKV infection influenced overall CNS growth, resulting in differences in TBV (ZIKV - Control median estimate (males) = 1.55, 89% HDI: [0.84, 2.23]; females = median estimate: 0.10, 89% HDI: [−0.60, 0.80]) and ICV, our overall measure of brain size and defined as total grey matter, white matter and CSF volume, between groups (ZIKV - Control median estimate (males) = 1.69, 89% HDI: [1.04, 2.32]; females = median estimate: 0.56, 89% HDI: [−0.07, 1.14]; see supplementary Fig. S1). Therefore, to distinguish region-specific structural (volumetric) effects and pathology from those driven by overall changes in brain size, we focused our analysis on regional volumes statistically adjusted for ICV, entering this as covariate in the statistical models.

Disrupted CSF volumes remained a core feature of ZIKV-induced pathology, though with notable sex differences. ZIKV-infected females had substantially greater total cerebrospinal fluid (CSF) volume than control females (median estimate = 1.36, 89% HDI: [0.80, 1.91]), yet exhibited decreased lateral ventricle volume (median estimate = −1.21, 89% HDI: [−2.11, −0.31]; Fig 6, 7). These findings indicate that the increased CSF in female infants after ZIKV infection is confined to the extra-axial space. ZIKV-infected males displayed a similar near-credible trend toward increased in total CSF volume (median estimate: 0.75, 89% HDI: [−0.004, 1.48]) compared to controls without differences in lateral ventricle volumes (median estimate: 0.88, 89% HDI: [−0.31, 2.08]; Fig 6, 7). These alterations in CSF suggest that ZIKV may disrupt important developmental roles of CSF in metabolic and neurochemical communication across brain regions with compensatory or pathological structural responses that differ between males and females.

**Figure 6.**
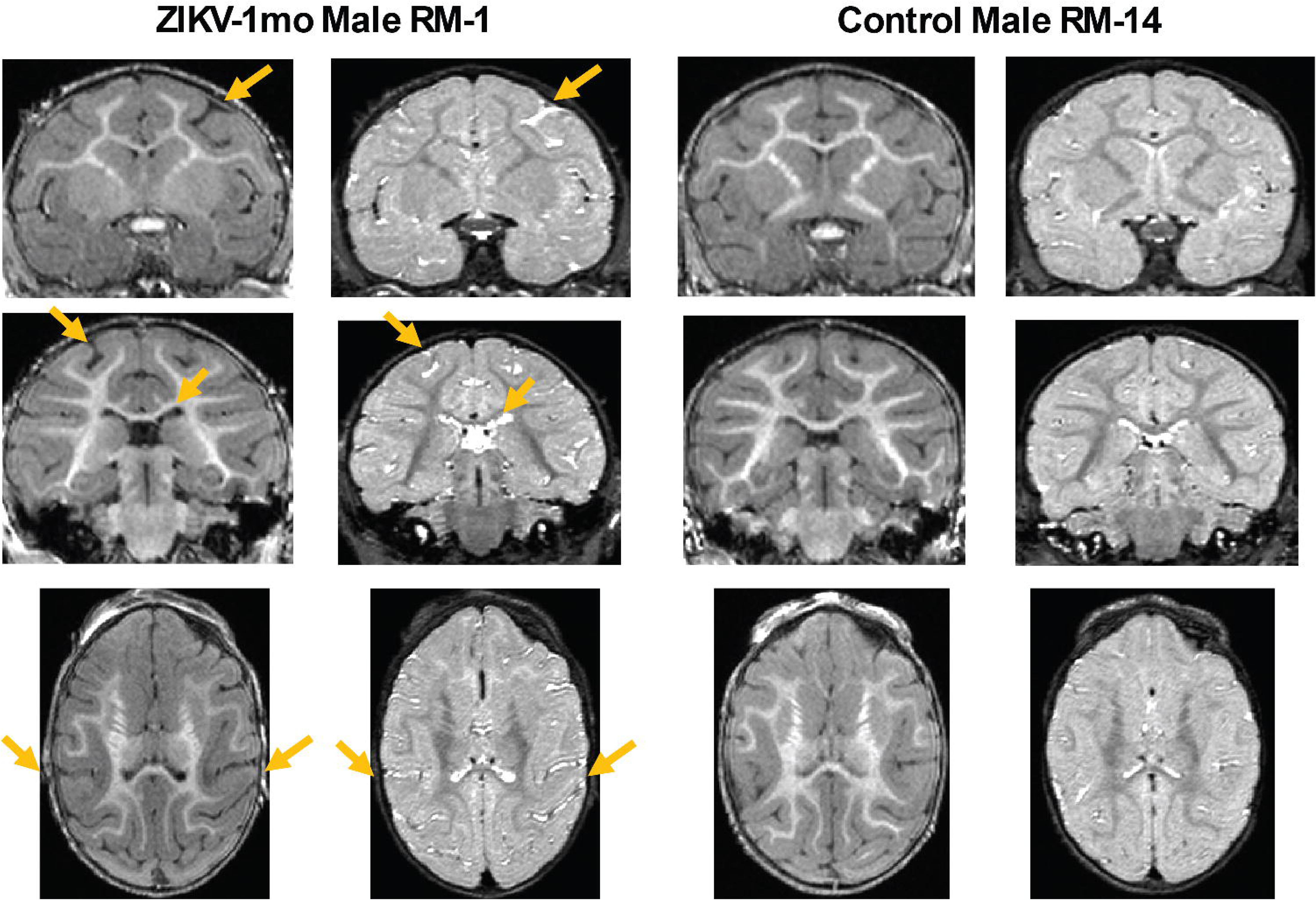
Increased CSF volume and widened sulci from postnatal ZIKV infection. T1- and T2-weighted MRIs from a representative ZIKV-infected male compared to an uninfected control male. Yellow arrows point to examples of sulci widening and increased CSF volume.

The limbic system displayed a striking, sex-specific impact of ZIKV infection affecting both subcortical and cortical structures. ZIKV-infected males showed a credible hypertrophy of the amygdala (median estimate = 1.52, 89% HDI: [0.56, 2.47]; Fig 7), suggesting a focal growth vulnerability in this emotion-regulatory center. In contrast, ZIKV-infected females showed no credible changes in amygdala size although volumetric reductions in the broader Temporal-Limbic cortex (combined gray and white matter) were seen. This atrophy was bilateral, affecting both the right (median estimate = −0.81, 89% HDI: [−1.30, −0.33]) and left (median estimate = −0.78, 89% HDI: [−1.30, −0.28]) hemispheres in ZIKV-infected females (Fig 7). There were no ZIKV vs. Control differences for hippocampal volume at this age, similar to prior reports ^25,30^.

**Figure 7.**
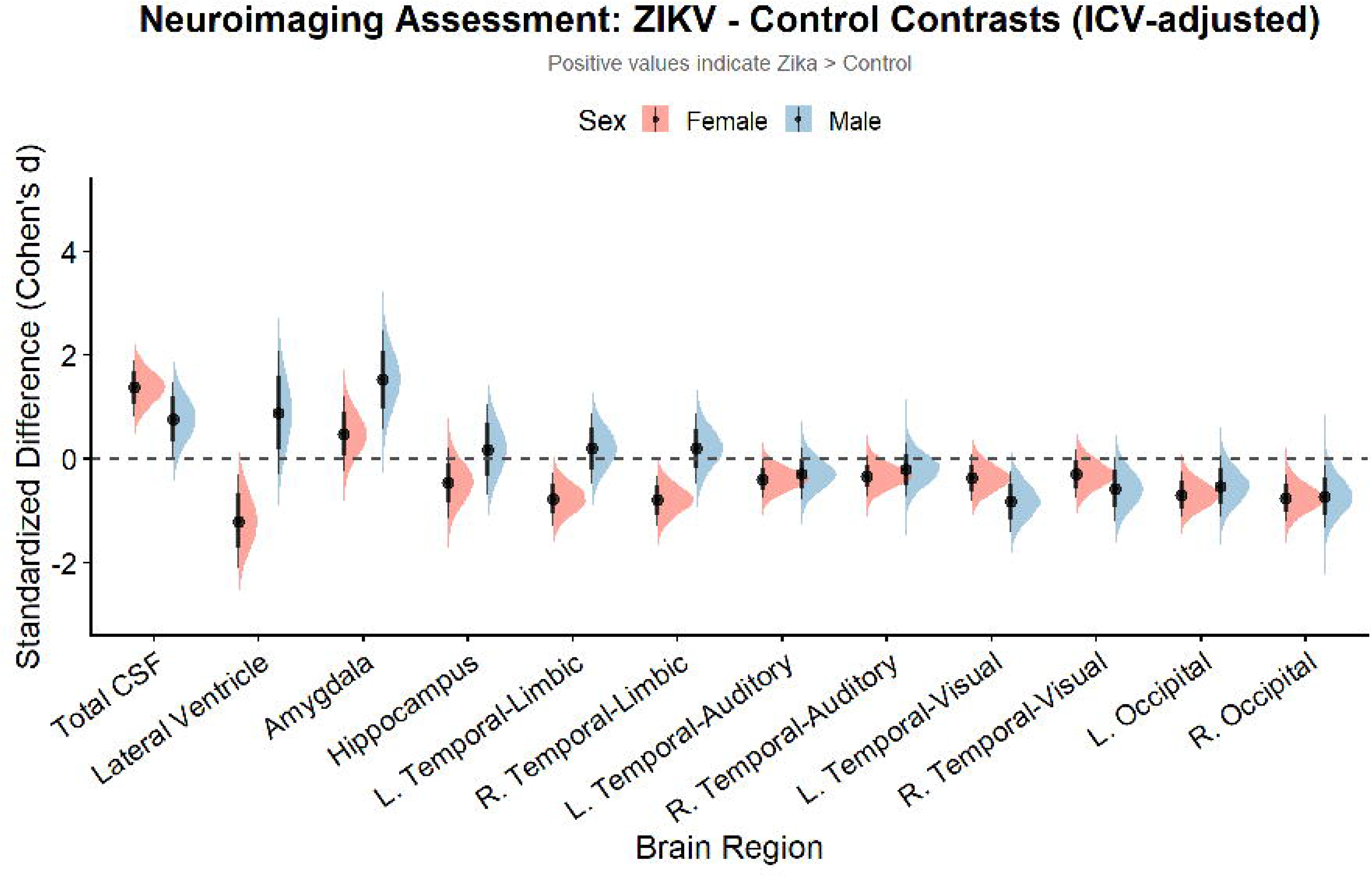
Sex dependent alterations in brain volume by ZIKV infection during infancy. Sex-specific posterior distributions of the ‘ZIKV - Control’ contrast of the brain region volumes across multiple brain regions obtained from the 3-month MRI scans (n = 6 ZIKV and 6 control (3 uninfected + 3 Poly-IC) per sex (24 scans total)). This plot displays the standardized difference between the ZIKV and control groups in different brain regions. Estimates are derived from separate Bayesian linear regression models for each region. For each region on the x-axis, the contrast is visualized side-by-side for females and males, distinguished by color. Each half-eye plot illustrates the full probability density of the estimated difference. The internal black lines denote the 89% (wider) and 66% (narrower, thicker) credible intervals, with the posterior median marked by a black dot. The horizontal dashed line at y=0 represents no difference between the groups; a contrast is considered meaningful when its credible interval does not overlap this line. L indicates left hemisphere and R indicates right hemisphere.

A widespread pattern of atrophy in sensory processing regions (combined gray and white matter) affected both sexes following ZIKV infection, though with hemispheric variations. Females displayed credible bilateral reductions in the occipital cortex (Right median estimate: −0.76, 89% HDI: [−1.21, −0.31]; Left median estimate: −0.70, 89% HDI: [−1.13, −0.26]), as well as a localized decrease in the left Temporal-Auditory cortex (median estimate: −0.40, 89% HDI: [−0.77, −0.02]; Fig 7). Males also exhibited deficits in the Temporal-Visual stream, with credible atrophy in the right occipital cortex (median estimate: −0.73, 89% HDI: [−1.33, −0.11]) and the left Temporal-Visual cortex (median estimate: −0.83, 89% HDI: [−1.41, −0.25]) compared to controls (Fig 7). All regions with credible or near-credible effects are summarized in supplementary Table S6. When the two control groups were analyzed separately (UIC, PIC, and ZIKV modeled as three groups; Supplementary Table S10), the greater amygdala volume in males, the increased CSF volume in females, the reduced temporal-limbic volume in females, the reduced lateral-ventricle volume in females, and the reduced left temporal-visual volume in males remained credible relative to both the naïve (UIC) and the immune-stimulated (PIC) control. The bilateral occipital and left temporal-auditory reductions, however, were credible relative to the Poly-IC control but not the naïve control; because Poly-IC is an acute innate-immune stimulus that does not reproduce the magnitude or duration of the immune response to ZIKV infection, these sensory-cortex effects are interpreted with caution and require confirmation in larger samples. A formal group-by-sex interaction analysis confirmed that these sex-specific effects reflect genuine differences between the sexes rather than a credible effect in one sex alongside a non-credible effect in the other. The interaction, i.e. the difference in the ZIKV effect between males and females, was very strong for the lateral-ventricle reduction (Cohen’s d = 2.09, 89% HDI [0.68, 3.42], probability of direction pd = 0.99) and the bilateral temporal-limbic reductions (right d = 1.00, [0.22, 1.70], pd = 0.98; left d = 0.98, [0.23, 1.74], pd = 0.98), strong for the male amygdala increase (d = 1.05, [−0.003, 2.15], pd = 0.94), and substantial for the female CSF increase (d = −0.61, [−1.41, 0.24], pd = 0.88) and the male left temporal-visual reduction (d = −0.46, [−1.08, 0.23], pd = 0.87). The occipital and auditory reductions showed no appreciable sex difference (pd 0.53 to 0.65) and are therefore not interpreted as sex-specific (Supplementary Table S11). Given the small number of animals in each sex-by-group subgroup, these sex-specific effects should be regarded as preliminary and require confirmation in larger cohorts.

These ICV-adjusted analyses provide critical insights into the nature of ZIKV-associated brain changes, revealing that the virus induces highly specific, sex-dependent reorganization rather than uniform pathology.

## DISCUSSION

This study demonstrates that ZIKV infection during early postnatal development in rhesus macaques leads to alterations in neuromotor, sensory orientation and emotional regulation functions during infancy that are context-dependent, accompanied by sex-specific changes in brain development. Importantly, other than the orientation deficits, these alterations are specific to ZIKV infection and not merely caused by innate immune activation during infancy. These findings extend our previous pilot work and provide compelling evidence that ZIKV neurotropism during infancy can have neurodevelopmental consequences well beyond when the original pathogen is no longer detectable. Notably, persistent elevations in extra-axial CSF and enlarged perivascular space volumes during infancy have been identified as early biomarkers for autism spectrum disorder (ASD) and subsequent deficits in motor development ^39–41^ and executive function ^42^.

Our most robust behavioral finding was the persistent alterations in emotional regulation that emerged during the week of acute infection and intensified through 9 weeks of age (poor state control scores). Emotional alterations were also detected at 5 months of age when ZIKV infected infants expressed more hostile behaviors compared to controls during the less salient threat conditions (Alone and Profile) of the human intruder paradigm. This pattern of hostile behavior during low threat conditions is atypical for rhesus on this paradigm and may be indicative of increased irritability ^43^. Irritability has similarly been reported in congenital ZIKV syndrome ^44–46^ and children with postnatal ZIKV exposure show poorer Personal-Social scores ^17^, suggesting shared impacts on emotion-regulatory circuits regardless of infection timing Unlike ZIKV exposure, innate immune stimulation during infancy did not result in heightened emotional reactivity, as the Poly-IC control group did not differ from uninfected controls. This suggests that persistent increases in emotional reactivity are not the result of a transient inflammatory response, but due to ZIKV triggering lasting changes in emotion regulatory circuits. Transcriptomic changes in regulatory brain regions recently revealed altered white matter micro-organization and aberrant neuronal expression of ASD risk genes in postnatally infected RMs ^47^. Permanent alterations of limbic system development and stress responsivity have been shown during critical developmental periods with other neurotropic viruses ^48–50^. Yet, the exact mechanism underlying these changes are still largely unknown.

Fetal ZIKV exposure has resulted in impaired motor, visual, and auditory function in humans and macaque models, even without microcephaly present at birth ^38,51–53^. Although there is limited data on postnatal ZIKV infection in humans, a prospective study showed that 15% of children with postnatal exposure exhibit adverse neurological outcomes by 20-30 months of age ^17^. The current study found ZIKV infected infant RMs had decreased neuromotor and orientation scores during early infancy compared to controls. Yet, when tested at older ages, performance on a visual acuity task revealed no significant difference between ZIKV infected and control RMs. The dissociation between early Orientation deficits and intact visual acuity at 4-6 months suggests these may reflect higher-order attentional processes or transient cortical delays that resolve by the end of the first year, as observed in fetal ZIKV RM model ^54^. The Poly-IC group also exhibited Orientation deficits, suggesting that early-life immune activation broadly impacts developing attentional systems, consistent with ZIKV-induced neuroinflammation disrupting the brain even without direct viral persistence ^47,53,55,56^. Continued monitoring of these animals is needed to determine whether neuromotor or visual issues worsen with age, as observed in congenital ZIKV syndrome ^26,51,52^.

A major limitation of our study is that infant RMs were not reared by their mothers, however, we have shown that supplementing our rearing techniques with increased socialization and a primary human caregiver results in species-typical social and emotional development ^57–60^. In fact, ZIKV-infected infant RMs exhibit a preference for their primary caregiver similar to controls, demonstrating a strong attachment bond that is similar to age-, sex-, and rearing-matched controls. Yet, ZIKV-infected RMs displayed sex-specific alterations in emotional expression during this caregiver separation, such that males showed blunted distress vocalizations compared to controls, while females showed no credible differences in distress behaviors (i.e. fearful, hostile and self-directed). These findings are consistent with ZIKV neuroinvasion altering regulatory systems modulating emotional responses but does not disrupt the core attachment bond. Two studies in fetal-exposure RMs showed intensified maternal attachment ^53,54^, with one study reporting blunted affective-cue integration during adolescence ^61^, underscoring complex ZIKV impacts on the social-emotional domain regardless of infection timing.

The neuroimaging findings revealed an increase in total CSF volume in both ZIKV infected sexes. Notably, this pattern diverges from our pilot data in postnatally infected female macaques, which showed persistent enlargement of lateral ventricles only ^25,26^, but is broadly consistent with numerous reports of ventriculomegaly and hydrocephalus in human congenital ZIKV infection ^62,63^. Together, these data show that increased CSF volume not just in ventricles, but in the extra-axial space may be a hallmark of ZIKV-induced neuropathology, regardless of early developmental timing of infection. ZIKV-infected females showed a credible decrease in lateral ventricle volume alongside their elevated total CSF, indicating that the excess CSF was confined to the extra-axial space rather than the ventricular compartment — a distinct pattern that suggests redistribution rather than simple overproduction. These neuroimaging findings are consistent with developmental alterations in CSF dynamics, which play an important role in brain communication, function and development ^64,65^. Sex-specific differences in choroid plexus transcriptomes can produce divergent ventricular phenotypes from the same upstream insult ^66–68^, potentially explaining the opposing lateral ventricle patterns between sexes. In humans, increased extra-axial CSF during infancy predicts ASD diagnosis, motor deficits, and poorer executive function ^39–42,69–71^, suggesting the CSF expansion in our ZIKV-infected infants may represent a candidate early biomarker warranting further investigation.

While ZIKV disrupted the CSF fluid homeostasis in both sexes, other compensatory mechanisms of neurodevelopment differed by sex. ZIKV-infected male RMs exhibited specific, focal hypertrophy in the amygdala. Conversely, females exhibited credible reductions in the temporal-limbic cortex, together with less robust occipital and auditory reductions that were credible relative to the immune-stimulated control only, paralleling findings in mouse models of in utero exposure that demonstrated greater neurological alterations in females and not males ^72^. Data on postnatal infection remain particularly sparse, and our previous macaque studies were limited to female-only cohorts ^25,26^. Notably, the Poly-IC control was not developmentally inert: it produced orientation deficits and heightened responses in the most salient stress condition. Inclusion of the Poly-IC control group provides a stringent comparison for the effect of innate immune activation and renders the pooled control groups (uninfected + Poly-IC) conservative for detecting ZIKV-specific effects. At the same time, because a single course of Poly-IC does not reproduce the magnitude, duration, or kinetics of the immune response to a replicating ZIKV infection, it remains an informative but imperfect proxy for the immune component of infection.

The male-specific amygdala enlargement, which persisted after adjusting for ICV, provides evidence for focal ZIKV-induced alterations in this critical threat detection and emotion-regulatory structure. This contrasts with amygdala volume reductions reported in both postnatal RMs ^26^ and prenatal human ZIKV exposure ^63^, suggesting divergent pathological pathways to emotional dysregulation. Hypertrophy in limbic structures may reflect neuroinflammatory edema or compensatory gliosis rather than healthy tissue growth ^39,73^. Indeed, histological assessments in infant macaques have shown that while ZIKV infection can induce neuronal apoptosis, it simultaneously triggers a robust inflammatory response in the limbic system characterized by activated microglia ^47^. While our previous study in females found amygdala volume reductions at 12 months ^26^, the current study shows amygdala enlargement in males, suggesting fundamentally different sex-specific pathological responses to early ZIKV infection. This discrepancy underscores the critical importance of including both sexes in neurodevelopmental research. Sex differences in fetal-maternal immune responses—including higher ZIKV RNA in the amniotic fluid of male fetuses ^53^ and hormonal modulation of susceptibility in mouse models ^72^—likely underlie the distinct volumetric patterns observed, underscoring the importance of including both sexes in neurodevelopmental studies.

The mechanisms underlying these persistent neurodevelopmental alterations remain to be fully elucidated. While ZIKV RNA peaked within days and cleared from plasma by two weeks post-infection, the observed brain changes at 3, as well as at 6 and 12 months in prior cohorts, suggest that initial viral neuroinvasion triggered cascading developmental disruptions that persist beyond acute infection ^25,26,56^. The dissociation between brain structure and behavioral changes suggests complex structure-function relationships. Transcriptomic work in this model demonstrates downregulation of ASD-risk genes alongside widespread microglial activation ^47^, while ZIKV-induced inflammation may disrupt choroid plexus barrier integrity and CSF dynamics through the ChP neuroimmune interface ^74,75^. Downregulation of oligodendrocyte differentiation genes and consequent white matter disorganization ^47^ may further drive behavioral deficits without proportional changes in gross regional volumes.

Comparing postnatal infection outcomes with fetal ZIKV exposure reveals both overlapping and distinct consequences. While congenital ZIKV syndrome primarily manifests with microcephaly, cortical malformations, and severe motor impairments ^6,44^, postnatal infection appears to produce more subtle alterations in brain structure, socioemotional behavior, and cognitive function without gross morphological abnormalities. However, shared features are emerging. Both fetal and postnatal infections alter species typical social and emotional reactivity ^25,26,54^. Fetal ZIKV exposure has additionally been associated with persistent visual pathway disruptions ^38,54^, and deficits in integrating affect with decision-making ^61^. While ventriculomegaly is a well-documented feature of congenital ZIKV infection and was observed in our prior pilot cohort ^25,26,62,63^, the current study reveals extra-axial CSF expansion as a feature of postnatal infection, suggesting that the compartmental distribution of excess CSF may differ by developmental timing. ZIKV infection during either window likely disrupts ChP-CSF signaling ^76,77^, with compartmental differences reflecting distinct timing-dependent vulnerabilities in postnatal versus fetal neurogenesis. Understanding these differential outcomes is essential for comprehensive clinical management of ZIKV-exposed children.

Several limitations warrant consideration. The pre-infection neuromotor differences between groups, though followed by progressive deterioration post-infection, indicates individual variability in typical development that can complicate the interpretation of infection outcomes, suggesting the need to include infant neurobehavioral assessment scores into randomized group stratification before infection studies. Because ZIKV challenge occurred at a median of 35 days of life (approximately 5 weeks of age), the assessments at 2 to 4 weeks of age precede infection. A sensitivity analysis estimating each week separately (age modelled as a factor; Supplementary Table S12) indicated that this pre-challenge difference is specific to the neuromotor domain: for state control, no ZIKV-specific difference (credible relative to both controls) was present before infection and the effect emerged only at the first post-infection assessment, whereas the small neuromotor difference was already credible at 3 weeks. We therefore interpret the neuromotor findings as reflecting, in part, pre-existing individual variation, while the emotional-reactivity findings are more confidently attributable to infection. The temporal mismatch between some neuroimaging and behavioral assessments limited our ability to assess direct brain-behavior relationships; thus, future longitudinal studies with closely matched timepoints are needed. Because these structure-function associations were not directly tested, and because cerebrospinal fluid physiology and viral presence within the central nervous system were not assessed in this cohort, our interpretations of altered CSF dynamics and ZIKV neuroinvasion are indirect inferences from structural imaging and prior work and should be regarded as hypotheses for future studies. The mechanisms underlying ZIKV neuropathology remain unclear at the macro-structural level of our neuroimaging approaches, though single-cell data suggests these are driven by specific cellular programs of apoptosis, inflammation, and differentiation^47^. Cellular and molecular approaches are needed to investigate the relative contributions of direct viral damage, acute neuroinflammation, and persistent inflammatory changes. The lack of a dengue-immune priming group limits interpretation, as prior flavivirus exposure is common in endemic regions and may modulate ZIKV pathogenesis ^78–81^, though recent prenatal studies in macaques suggest ZIKV exposure disrupts neurodevelopment regardless of maternal dengue immunity ^54^. Finally, whether these alterations persist, worsen, or resolve over longer developmental periods requires continued longitudinal follow-up through juvenile and adolescent stages.

Despite these limitations, our findings raise hypotheses about potential clinical and public health consequences of ZIKV exposure that extend beyond congenital Zika syndrome. Given that children account for 10-31% of ZIKV infections in endemic areas ^15–17^, often through mosquito bites or breast milk transmission ^18^, and that postnatal infections may be mild or subclinical, the potential burden of subtle neurodevelopmental sequelae could be substantial in regions with endemic transmission. The increased emotional reactivity and attention deficits observed here may presage longer-term difficulties with emotional regulation, anxiety, behavioral control, and academic performance. Early identification and developmental surveillance of children with documented or suspected postnatal ZIKV exposure may be warranted, particularly in the first 2-3 years of life when emotional and attentional systems undergo rapid maturation. Screening tools assessing emotional reactivity, attention, neuromotor control and visual-spatial processing could be incorporated into well-child visits in endemic regions. Importantly, given that behavioral interventions targeting emotional regulation and attention can be effective when implemented early ^82,83^, timely identification could enable interventions to mitigate long-term impacts. The pronounced male vulnerability in brain volumetric changes suggests that sex-specific surveillance and intervention approaches may be needed, though the presence of behavioral alterations in both sexes indicates that all ZIKV-exposed infants warrant monitoring.

In conclusion, this study provides robust evidence that ZIKV infection during early postnatal development produces alterations in emotional reactivity that persist through infancy. It also causes sex-specific brain and CSF changes—male amygdala hypertrophy, female temporal-limbic atrophy, and elevated extra-axial CSF in both sexes—that may serve as early biomarkers of neurodevelopmental risk. These results motivate future longitudinal studies to determine whether neurodevelopmental surveillance and targeted early interventions are warranted for children with postnatal ZIKV exposure.

## METHODS

### Animals

Twenty-four infant Indian RMs (*Macaca mulatta*), 12 males and 12 females, were included in this study. Twelve infants were infected with ZIKV at ∼1 month of age (median age 35 days; ZIKV-1; 6 male, 6 female), six infants served as age-, sex-, and rearing-matched uninfected controls (UIC; 3 male, 3 female), and six infants served as an innate immune stimulation group (Poly-IC; 3 male, 3 female). The infants were delivered naturally by their dams while housed in indoor/outdoor social groups at the Emory National Biomedical Research Center (ENBRC). All dams were from the specific pathogen-free colony (negative for Herpes B, SIV, SRV, and STLV1), had not been previously used for infectious disease or vaccine studies, and did not have any clinical signs of infection during pregnancy. Infants were then removed from their dams at 12-15 days of age and transported to the ENBRC nursery facility. All infants were pair-housed in warming incubators for the first 3-4 weeks. Infants were hand-fed formula every 2–3 h for the first month, then via self-feeders until 6 months of age, as per standard ENBRC protocol. Soft blankets, plush toys, and hanging fleece surrogates were provided and changed daily by a primary human caregiver that handled and socialized with the infants 3-4 times per day for a total of 6 h daily. Soft chow (Purina Primate Chow, Old World Monkey formulation) and fruits were introduced starting at 1 month of age. Water was provided *ad libitum*. At 5-6 weeks of age, infants transitioned into age-appropriate caging and housed in pairs or trios consisting of full tactile, visual, and auditory contact with conspecifics. During the acute ZIKV-infection period, infants were housed in an ABSL-2+ (Animal Biosafety Level 2 enhanced) room until the virus was no longer detected in blood or urine. All housing was indoors on a 12 h light–dark cycle.

All procedures in this study were approved by the Emory University Institutional Animal Care and Use Committee and were conducted in an AAALAC accredited facility in full compliance with the United States Public Health Service Policy on Humane Care and use of Laboratory Animals.

### Virus and infection

The ZIKV of Puerto Rican origin (PRVABC59, GenBank accession number: KU501215.1) used in this study had been passaged four times and titrated on Vero cells. Experimental infections were performed via the subcutaneous route using 10^5^ plaque forming units (pfu) of ZIKV PRVABC59.

### ZIKV RNA detection by qRT-PCR

Total RNA was extracted from 140 μl of plasma samples using the QIAamp Viral RNA Mini Kit (Qiagen). RNA from urine was isolated using the QIAamp Circulating Nucleic Acid Kit (Qiagen) for the extraction of larger volumes (up to 2 ml). ZIKV RNA standard was generated by annealing two oligonucleotides spanning the target ZIKV prM-E gene region and performing in vitro transcription using the MEGAscript T3 Transcription Kit (Ambion). Purified RNA was reverse transcribed using the High-Capacity cDNA Reverse Transcription Kit (Applied Biosystems) and random hexamers. For quantitation of viral RNA, a standard curve was generated using 10-fold dilutions of ZIKV RNA standard, and qRT-PCR was performed using TaqMan Gene Expression Master Mix (Applied Biosystems) and ZIKV primers (1 μM) and probe (250 nM). The ZIKV primer-probe set targeting the prM-E gene region included forward primer ZIKV/PR 907 (5′-TTGGTCATGATACTGCTGATTGC-3′), reverse primer ZIKV/PR 983c (5′-CCTTCCACAAAGTCCCTATTGC-3′), and probe ZIKV/PR 932 FAM (5′-CGGCATACAGCATCAGGTGCATAGGAG-3′). The probe was labeled with 6-carboxyfluorescein (FAM) at the 5′ end and two quenchers, an internal ZEN quencher and 3′ end Iowa Black FQ (Integrated DNA Technologies). The standard curve had an R2 value greater than 0.99. Viral RNA copies were interpolated from the standard curve using the sample cycle threshold (CT) value >40 were deemed undetectable. For plasma samples, viral RNA copies were represented as copies per milliliter of plasma with a limit of detection (LOD) 939 copies/ml.

### Plasma Cytokines

Cytokine concentrations were measured in plasma samples collected before and 24 hours after injection from Poly-IC compared to uninfected controls. Unfortunately, after ZIKV viral quantification, there was not enough plasma remaining for cytokine assay from the ZIKV-infected group. Plasma was analyzed by the Emory Multiplexed Immunoassay Core (Emory University, Atlanta, GA) using a multiplex electrochemiluminescence immunoassay kit from Meso Scale Discovery (Rockville, MD) following the manufacturer’s protocols for the U-Plex Custom Biomarker (NHP) Assays kit (Catalog #K15068M) and plates were read with the MSD MESO QuickPlex SQ 120MM. Cytokine responses were expressed as log-fold changes at 24 h relative to baseline values.

### Neurobehavioral Assessment

We evaluated neonatal macaque neurobehavior with a well-validated assessment of developed infants for rhesus macaques, Infant Neurobehavioral Assessment Scale [INAS], adapted from the study by Schneider and Suomi ^31,32,84^, which is based on the human Brazelton Newborn Behavioral Assessment Scale ^85^. Twenty-nine test items in the INAS aligned with the neurodevelopmental areas of interest and make up the Orientation, Motor maturity and activity, Neuromotor (sensory) responsiveness, and State Control constructs. This neonatal neurobehavioral test has previously been used to define neonatal development of prenatally ZIKV-exposed infants ^53,86^. Ratings were based on a five-point Likert scale ranging from 0 to 2. The INAS was administered weekly between 2-9 weeks of age. During the week of ZIKV infection or Poly-IC administration, INAS testing was performed at 6 days post-infection, to avoid potential confounds from handling or sedation during sample collections, as well as avoid peak viremia.

Four examiners (J.R., R.R., K.L., A.V.S.) were present for all neurobehavioral testing and scoring to ensure test administration reliability (Cohen’s Kappa = 0.9). Items were administered in a consistent sequence across all animals to optimize performance and decrease handling time. Assessments were hand-scored on a printout of the scoring form during administration. Higher scores reflect optimal scores for Orientation, Motor, and Neuromotor, whereas higher State Control scores reflect poor emotional regulation and higher reactivity.

### Acute Stress Assessment

The potential impact of postnatal ZIKV infection on emotional regulation was examined using the Human Intruder Paradigm ^34^ that is based on the Stranger Approach task for assessing behavioral inhibition and anxiety in children ^33^. Infants were previously trained to quickly transfer from their home cage to a transport box. On the testing day, the subject was separated from the cagemate, transported from the nursery to a novel testing room, and placed in a modified housing wire-mesh cage, with one side in clear acrylic to allow for unobstructed viewing of the animal’s behavior. The paradigm consisted of three conditions: Alone condition, animal remained alone in the cage for 9 min to acclimate to the environment and obtain a baseline measure of behavior. Profile condition: An unfamiliar person (researcher wearing a rubber mask) entered the room, sat on a stool 2 m from the test cage, while presenting his/her profile (without eye contact) to the animal for 9 min. Then, the person left the room, giving the animal a 3-min break. Stare condition: The unfamiliar person reentered the room, but this time stared directly at the animal for 9 min. Two examiners (S.F., K.L.) with good inter-rater reliability (Cohen’s Kappa = 0.88) scored all behaviors.

### Attachment Assessment

To evaluate whether postnatal ZIKV infection would impact attachment bonds, infant RMs were evaluated using a choice discrimination between two stimuli, a primary human caregiver and another familiar human ^57^. The 10-minute discrimination task was performed in a large rectangular enclosure (5’ x 5’ x 10’) made of wire mesh and several clear acrylic panels for unobstructed behavioral observation. Infants entered the enclosure between the two stimuli and were allowed to freely explore the large enclosure. Infants could choose between spending time near their primary human caregiver or the other familiar human. A 2.5’ square area immediately in front of the caregiver or familiar human was considered their proximity zone, which was used to calculate an index of preference. The index of preference is calculated by finding the difference between the time spent in each proximity zone and dividing by the total time spent in both proximity zones. This results in a preference score ranging from –1 to +1 which allows for assessing whether the infant has a preference for the primary human caregiver (+1) as opposed to the familiar human (−1) or no preference (0). Three examiners (M.A., K.L., A.V.S.) scored all the behavioral testing and had good inter-rater reliability (Cohen’s Kappa = 0.89).

The behavioral ethogram for acute stress assessment and attachment assessment is included in supplementary materials (Table S1).

### Neuroimaging assessment

To investigate the potential impact of postnatal ZIKV infection, structural MRI was performed at 3 months of age using a Siemens 3T Tim Trio system (Siemens Medical Solutions) with a Tx/Rx 8-channel volume coil. Data were acquired in a single session, which included T1-and T2-weighted structural MRI scans. Animals were scanned in the supine position in the same orientation, achieved by placement and immobilization of the head in a custom-made head holder via ear bars and a mouthpiece. After initial telazol induction and intubation, scans were collected under isoflurane anesthesia (0.8–1%, inhalation, to effect).

End-tidal CO2, inhaled CO2, O2 saturation, heart rate, respiratory rate, blood pressure, and body temperature were monitored continuously and maintained during each MRI session. High-resolution T1-weighted MRI scans were acquired for volumetrics using a three-dimensional magnetization prepared rapid gradient echo (3D-MPRAGE) parallel image sequence [repetition time/echo time (TR/TE) = 2600/3.46 ms; field of view (FOV), 116mm× 116 mm; voxel size, 0.5 mm3 isotropic; six averages] with GeneRalized Auto calibrating Partially Parallel Acquisitions (GRAPPA) acceleration factor of R = 2. T2-weighted MRI scans were collected using a 3D fast spin-echo sequence (TR/TE = 3200/373 ms; FOV, 128mm× 128 mm; voxel size, 0.5 mm3 isotropic; three averages) with GRAPPA (R = 2) to aid with registration and delineation of anatomical borders.

Structural MRI data processing and analysis data sets were done using AutoSeg_3.3.2 segmentation package ^87^ to generate the volumes of brain WM and GM, CSF, and cortical (temporal visual area, temporal auditory area, and prefrontal, frontal, parietal, occipital lobes, and cerebellum) and subcortical (hippocampus, amygdala, caudate, and putamen) brain areas. Image processing steps included the following: (i) averaging T1 and T2 images to improve signal-to-noise ratio, (ii) intensity inhomogeneity correction, (iii) rigid-body registration of the subject MRI to the 3-month UNC-Emory infant RM atlases ^88^, (iv) tissue segmentation and skull stripping, and (v) registration of the atlas to the subject’s brain to generate cortical parcellations (affine followed by deformable ANTS registration).

Cortical and subcortical regions were defined based on neurohistological, cytoarchitectonic and connectivity/functional criteria ^90–92^ as well as published macaque MRI anatomical parcellations ^12,93^ and previous publications by our group ^25,26,94^. ICV was defined as total GM + total WM + total CSF in the ventricles and in the subarachnoid cavity, and volumes of each cortical lobe (temporal auditory, temporal visual, occipital) were defined as the total GM + total WM ^88,94^.

### Statistical Analyses

For each task, bespoke statistical analyses were conducted within a Bayesian framework using the R statistical environment (version 4.4.0) and the *brms* package (version 2.21.0) ^95^. This approach was chosen for its flexibility in handling complex hierarchical models and its ability to formally incorporate prior knowledge. Inference was based on the full posterior distribution: for each effect we report the posterior median with an 89% highest-density interval (HDI), and we describe an effect as credible when its 89% HDI excludes zero. Because a single interval can be misleading at small sample sizes, we also report the probability of direction (the posterior probability that an effect lies in the estimated direction), so that evidence is interpreted as graded rather than dichotomous. Priors were chosen to reflect available knowledge while remaining conservative: where suitable data existed we synthesized informative priors from previously published control cohorts (for the INAS domains and the neuroimaging control volumes), and weakly-informative priors were used elsewhere. Such priors regularize the estimates and are particularly valuable at the sample sizes available here (six animals per group, three per sex), where they stabilize otherwise ill-conditioned models; for example, the low-frequency attachment behaviours produced unstable, near-separated estimates under flat default priors that were resolved by weakly-informative priors. Reporting the full posterior with an HDI and the probability of direction provides graded, interpretable evidence that degrades smoothly as the sample shrinks, in contrast to the unstable dichotomous outcomes of null-hypothesis significance testing at small sample sizes. Throughout, we interpret effects cautiously, distinguish those that are robust across both control groups and across sensitivity analyses from those that are more tentative, and identify the sex-specific and small-subgroup findings as requiring confirmation in larger cohorts.

For robust inference based on matching sample sizes between the test and control groups, two control groups (PIC and UIC) were pooled together into one control group for the following analyses: visual acuity, acute stress, attachment, and neuroimaging. We confirmed that pooling was justified with a sensitivity analysis that refit each model using a three-level group factor (UIC, PIC, and ZIKV) and extracted, within each sex, the ZIKV vs. UIC, ZIKV vs. PIC, and PIC vs. UIC contrasts; the full contrasts are reported in Supplementary Tables S7-S10.

For the ZIKV-infected group only, we additionally examined viral-load burden (the area under the viremia curve, AUC): we tested whether burden differed by sex (log(AUC) ∼ Sex) and whether standardized log-AUC was associated with each principal outcome, entering it as a predictor in that outcome’s model family and, for the sex-specific findings, within the relevant sex. These exploratory analyses used weakly-informative priors, were refit excluding one animal with atypically low burden, and are reported in full in Supplementary Table S13. Specifics of each task are described in detail below:

### Neurobehavioral Assessment

Given the ordinal, Likert-scale nature of the INAS constructs, four separate Bayesian cumulative ordinal regression models were fit, one for each primary outcome: Orientation, Neuromotor, Motor, and State Control. Each model utilized a logit link function. The full model structure included fixed effects for Treatment (UIC, PIC, ZIKV), and Age (in weeks, as a continuous predictor), along with their two-way interaction to explore how the effect of treatment might change over time. To account for the non-independence of repeated observations from the same subject, a random intercept was included for each Animal ID. Models were fit only on data with non-missing values for their respective dependent variable (i.e., via listwise deletion for each outcome). To verify that the timing of the treatment effects was not an artefact of modelling age as a continuous linear predictor, we additionally refit each INAS model with age as a factor (Treatment by factor(age in weeks)), which estimates the treatment contrast at each week independently; these per-week contrasts (posterior median, 89% HDI, and probability of direction) are reported in Supplementary Table S12.

An informative prior specification strategy was employed, leveraging data from previous control animal studies [^32,86^; unpublished data]. This prior information was first synthesized by calculating sample-size weighted means and pooled standard deviations to establish robust expectations for control group behavior. These expectations directly informed the priors for the model cutpoints (intercepts) on the latent log-odds scale. For example, for outcomes with higher expected mean scores (e.g., Orientation, ∼1.3), priors for the cutpoints were centered slightly lower to reflect the increased probability of observing higher scores, whereas for outcomes with lower expected means (e.g., State Control, ∼0.8), priors were centered higher.

Priors for the fixed effect coefficients (βs) were specified to distinguish between control-related effects and effects involving the novel ZIKV treatment. The difference between the two control groups (PIC vs. UIC) was assigned a tight prior centered at zero (Normal(0, 0.4)), reflecting a strong prior belief in their similarity. Other control-related effects, such as Sex and Age, were assigned weakly informative priors (Normal(0, 0.8)). In contrast, all effects involving the ZIKV group (both its main effect and its interactions) were assigned less informative, wider priors (Normal(0, 1.5)) to reflect the absence of prior experimental data and to allow the current data to primarily drive the posterior estimates for this group. The standard deviation of the random intercepts was given a robust, weakly informative Student’s t-distribution prior (Student-t(3, 0, 2.5)).

Models were fit using Markov Chain Monte Carlo (MCMC) sampling with four chains, each running for 3,000 iterations with a 1,000-iteration warmup period. Model convergence was assessed by ensuring potential scale reduction factors (R-hat) were below 1.01 and by visual inspection of trace plots. Posterior predictive checks were conducted to evaluate model fit. To investigate specific group differences, pairwise contrasts were computed directly from the model’s posterior distribution as emmeans package only has an experimental implementation for ordinal regression models. A comprehensive grid of predictor combinations (Treatment, Sex, and Age at low (0.1 percentile), medium (0.5 percentile), and high (0.9 percentile) values) was created, and the brms::posterior_linpred() function was used to generate posterior draws of the latent predictor for each combination. The posterior distribution for each contrast (e.g., ZIKV vs. UIC) was then calculated by directly subtracting the draws of the corresponding groups. An effect was considered credible if its 89% Highest Posterior Density Interval (HDI), calculated using the tidybayes package, did not contain zero ^96^.

### Visual Attention Task

The primary dependent variable, “% looking,” was measured as a percentage and is treated as a continuous outcome. Because percentage data are bounded between 0 and 100 and our observations occasionally attained the bound of 100%, we first transformed the data to the open interval (0, 1) using the Smithson and Verkuilen (2006) transformation ^97^. Specifically, each percentage was scaled by dividing by 100 and then adjusted via the formula: adjusted value = ((y × (N − 1) + 0.5) / N), where *y* is the raw proportion and *N* is the total number of observations. This transformation minimizes boundary issues and renders the data appropriate for beta regression analysis.

Given the non-normal, continuous, and bounded nature of the transformed variable, analyses were performed using Bayesian beta regression implemented via the *brms* package in R. Missing values in “% looking” were incorporated directly into the model using the built-in Bayesian imputation capability (via the mi() function in *brms*), allowing joint inference on both the missing and observed data. The regression model included fixed effects for the group (UIC, PIC, or ZIKV; coded as a factor) and its interactions with additional predictors—contrast level and age. Although age was recorded numerically, it was modeled as a categorical variable to capture potential nonlinear or group-specific effects.

Prior to analysis, exploratory tests were conducted to assess differences between the control groups UIC and PIC. When pairwise comparisons using estimated marginal means (EMMs; computed via the *emmeans* package) indicated that groups UIC and PIC did not differ statistically (89% HDI did included 0), these groups were collapsed into a single group (hereafter “control”), which was then compared with ZIKV group. Random intercepts were included for left/right acuity, acuity number, and AnimalID to account for the hierarchical structure and potential non-independence of repeated measurements within subjects and testing sessions.

The full model specification was formulated as follows:

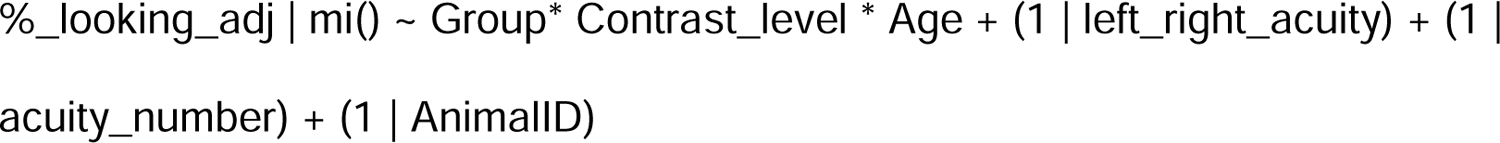

Posterior distributions were obtained using Markov chain Monte Carlo (MCMC) sampling with four chains, a total of 4,000 iterations per chain (including warmup), and appropriate adjustments to sampler parameters (e.g., increasing adapt_delta) to address potential divergent transitions and ensure convergence (assessed via Rhat statistics, effective sample sizes, and trace plots). Model convergence was further evaluated with posterior predictive checks.

Pairwise comparisons of treatment effects were performed using the *emmeans* package to estimate marginal means and compute posterior contrasts for each combination of age and contrast level. Results were summarized as posterior means with 89% HDI credible intervals; differences were deemed significant when the 89% intervals did not include zero. Additionally, posterior distributions for these contrasts were visualized using *tidybayes* and *ggplot2* to generate “half-eye” plots that display both the density estimates and credible intervals, with distinct color coding reflecting different age groups and contrast levels.

This integrative Bayesian approach allowed for robust estimation and direct probabilistic interpretation of the group differences on % looking while accommodating the complexities of the data, including non-normality, bounded outcomes, and missing data.

### Acute Stress Assessment

Behavioral outcomes for the acute stress assessment were categorized into two groups: (1) behaviors with prior control data available ^25,34,59,84,98–100^ and (2) behaviors without prior control data. All outcomes, including frequencies and durations (rounded to the nearest integer) were modeled using negative binomial regression with a log link function to account for overdispersion.

All behavioral outcomes included the predictors: Group: levels were “Control” (reference level) and “ZIKV”; Sex: levels were “Female” (reference level) and “Male” and Condition: levels were “Alone” (reference level), “Profile,” and “Stare”

The specified model included a full three-way interaction:

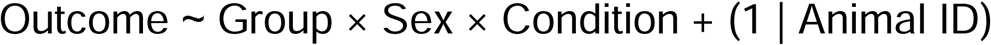

#### Models for Behaviors with Prior Control Data

For behaviors with historical control data weighted means and standard errors were computed from previously published studies ^25,34,59,84,98–100^ including unpublished data. The weighted mean for each behavioral outcome and condition was calculated using sample size as weights. Standard errors were also computed using a weighted average method.

Intercept priors were established on the log scale using the weighted control means for the baseline condition (“Alone”). For instance, the intercept prior for the baseline was calculated as:

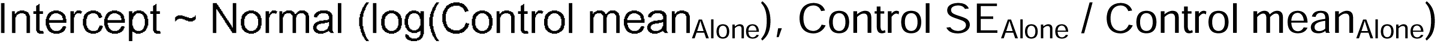

Condition contrasts (“Profile” and “Stare” relative to “Alone”) were similarly calculated on the log scale, with their standard errors derived using error propagation (delta method):

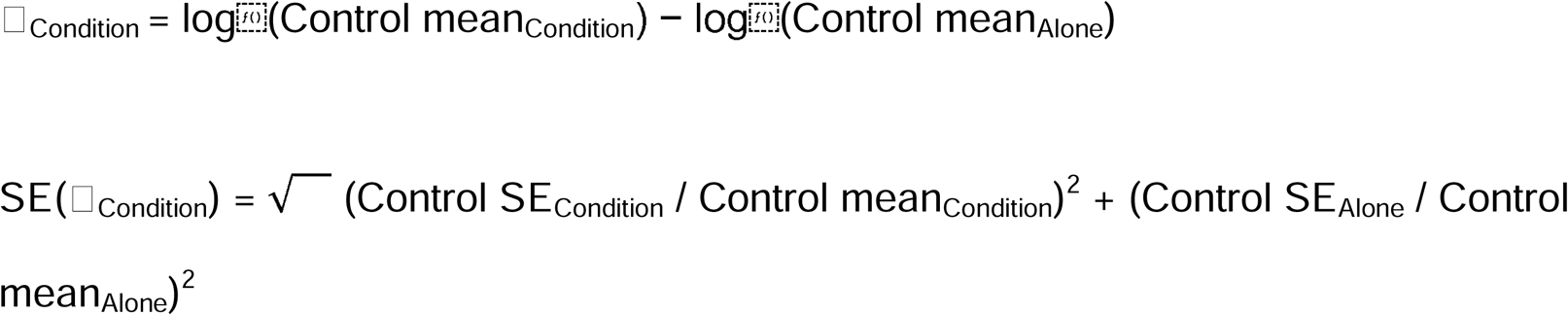

Additional weakly informative priors included sex difference: Normal(0,1), treatment and interactions: Normal(0,1), overdispersion parameter (shape): Exponential(1).

*Models for Behaviors without Prior Control Data:* Behaviors without prior control data were assigned generic weakly informative priors including intercept: Normal(0,5), main effects and interactions: Normal(0,1), overdispersion parameter (shape): Exponential(1).

Post-hoc pairwise contrasts for all main effects and interactions were calculated using the emmeans package. All estimates are reported on the log-response scale (log rate ratios). We report 89% credible intervals, which balance stability and precision and can be interpreted as having an 89% probability of containing the true population value.

#### Model Fitting and Validation

All models were fit using the brm() function in the brms package in R. Each behavior was modeled separately in iterative loops with 4 chains per model and 4000 iterations per chain (including 1000 warm-up iterations). Convergence assessed using potential scale reduction factor (R□) and effective sample sizes. Posterior predictive checks were conducted to ensure adequate model fit and accurate representation of observed data variability.

### Attachment Assessment

Each behavioral outcome was modeled separately to assess the differences in each behavior as well as to account for differences in measurement scale and data structure. We tested the differences in the behavioral variables between controls (uninfected controls (UIC), Poly-IC (PIC)), and ZIKV infected groups and their interactions with sex. We leveraged data from previous experiments performing this task ^57,101,102^ to specify informative priors for the control group. We weighted the previous means and standard errors with the sample size to obtain the value for specifying priors. For the low-frequency behaviors, which were prone to separation when the small control groups were divided by sex, we used weakly-informative priors (Normal(0, 1)) on the group and sex coefficients, consistent with the specification used for the scream and index-of-preference models.

#### Modeling of Frequency Outcomes

Behavioral responses recorded as frequencies were modeled using negative binomial regression with a log link. These included the frequencies of screams, attention seeking behaviors, affiliative behaviors, fearful behaviors, hostile behaviors, anxious behaviors, displacement behaviors, and falls. Self-directed behaviors, measured as a duration was also modeled similarly. A negative binomial model is better suited for count data compared to a poisson model when overdispersion is a concern.

#### Prior specification

Of the included behaviors, prior information was available from earlier experiments for the number of screams ^57,101,102^. As there were multiple prior studies with different sample sizes, the prior mean frequency was obtained by weighting the means with the sample sizes in the studies. These studies were largely balanced by sex, so no weighting based on sex was necessary. The data (i.e. mean frequency and standard error) were first transformed to the log scale using the relation:

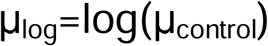

and the standard error was approximated using the delta method:

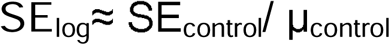

Based on the previous studies, the weighted control mean frequency was 32 with a weighted standard error of 22. Thus, the intercept prior was specified as a normal distribution with mean log(32)≈3.466 and standard deviation 0.688. In contrast, group effects (i.e. the differences between the experimental groups and control), as well as the sex main effects and interactions, were assigned weakly informative priors (e.g., Normal(0,1)) because no separate previous estimates were available for these effects. Each model was thus parameterized as:

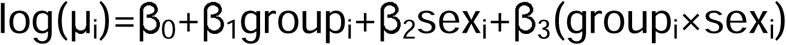

with the intercept β_0_ informed by prior control data, and subsequent coefficients β_1−3_ assigned diffuse (zero-centered) priors.

#### Modeling the Index of Preference

The Index of Preference was measured on a bounded scale from –1 to 1. To model this outcome appropriately, it was first transformed to the (0, 1) interval via the transformation:

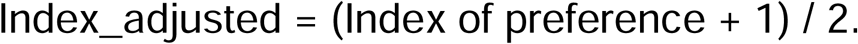

The transformed variable was then modeled using a beta regression with a logit link. Based on previous control-group data ^57,101,102^, the weighted mean of index_adjusted was estimated to be 0.9 with a weighted standard error of 0.0532. When working with a beta regression with a logit link, the intercept parameter is interpreted on the logit scale. To obtain the corresponding prior for the intercept:

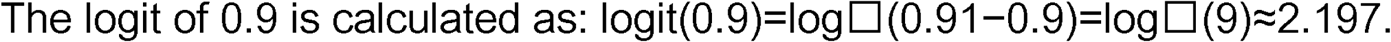

Using the delta method, the standard error can be obtained as: SE_logit_≈∣g′(p)∣×SE_raw_ where g(p) is the logit of the mean calculated above.

The derivative of the logit function at p is: d/dp (logit(p))=1 / p(1-p).

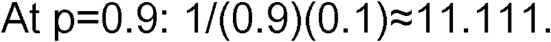

Hence, the standard error on the logit scale is approximated as:

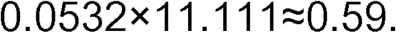

Thus, the intercept for the beta regression was specified with a prior of Normal(2.197,0.59).

Treatment and sex effects were again modeled with weakly informative priors centered at zero.

#### Pairwise Contrasts and Posterior Diagnostics

For all models, estimated marginal means (EMMs) were computed using the *emmeans* package to facilitate pairwise comparisons between groups within each sex. These contrasts were examined to determine whether significant differences existed between conditions. In addition, posterior predictive checks were conducted, and diagnostic plots of the posterior distributions of the fixed effects were generated using the *tidybayes* and *ggplot2* packages. These visualizations provided representations of the parameter estimates and assisted in assessing model convergence and overall fit.

### Neuroimaging Assessment

To investigate the impact of ZIKV infection on neurodevelopment, we modeled the volume of multiple brain regions separately. We chose the following regions based on their relevance in the behavioral metrics we studied: Total CSF, Lateral Ventricles, Amygdala, Hippocampus, Temporal-Limbic Cortex, Temporal-Auditory Cortex, Temporal-Visual Cortex, and Occipital Cortex. For the cortices, we modeled the left and right separately. Our approach of modeling brain regions separately accounts for the unique volumetric properties of each region and allows for region-specific inferences. Prior to the main analyses, the two control groups—uninfected controls (UIC) and Poly-IC (PIC) animals—were compared on key global measures, Total Brain Volume (TBV) and Total Intracranial Volume (ICV), using independent samples t-tests. As no significant differences were found, the groups were combined into a single ‘Control’ group for all subsequent analyses to increase statistical power. For each brain region, we then conducted a series of Bayesian linear regression analyses to test for differences between this combined control group and the ZIKV-infected group, as well as to assess potential interactions with sex. A key feature of our analysis was the integration of historical data on brain region volumes from previous neuroimaging experiments ^25,103^ and our unpublished data to construct informative priors for the baseline (control group) volume of each brain region. We used standard weakly informative priors for the ZIKV group as no prior data was available for the ZIKV group.

#### Modeling of Volumetric Outcomes

All brain region volumes were modeled directly on their raw scale, measured in cubic millimeters (mm³), using a Gaussian likelihood. This approach was chosen to maintain the natural interpretability of the model parameters and to allow for the direct application of informative priors derived from historical data, which were also on the raw mm³ scale.

#### Prior Specification

To directly incorporate sex-specific historical data, a no-intercept model parameterization (∼ 0 + group:sex) was used. This approach explicitly models the mean volume for each of the four experimental subgroups (e.g., Control Females, ZIKV Males), allowing priors to be set on each group’s mean parameter directly.

#### Priors for Control Group Parameters

For each brain region, historical control data ^25,94,103,104^ was stratified by sex. The mean (μ_prior) and standard deviation (σ_prior) of volume were calculated separately for historical control females and control males from this external data. These statistics were used to define informative priors on the raw (mm³) scale for the corresponding model coefficients. The prior for the groupControl:sexFemale coefficient was set to Normal(μ_prior_female_, σ_prior_female_). The prior for the groupControl:sexMale coefficient was set to Normal(μ_prior_male_, σ_prior_male_).

#### Priors for ZIKV Group Parameters

As no historical data was available for ZIKV-infected animals, weakly informative priors were assigned to the coefficients for the ZIKV subgroups. These were specified as Normal(4000, 20000), a diffuse prior on the raw mm³ scale that allows a wide range of values and lets the data dominate inference.

#### Model Structure and Covariate Adjustment

A four-level interaction term, group:sex, was created to represent each of the four experimental subgroups. The models were fit without an overall intercept to estimate the mean for each of these levels directly.

To determine if regional volume differences were disproportionate to an animal’s overall size, models included standardized Total Intracranial Volume (ICV) as a covariate:

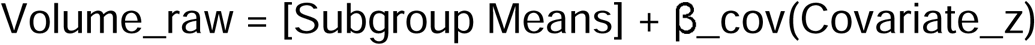

A wide prior of Normal(0, 3000) was used for β_cov as the volume was on raw scale. This allowed the data to be the primary influence on the estimates. This adjusted analysis is essential for distinguishing between a region being smaller simply because the entire brain is smaller, versus the region being atypically small even for a given brain size. Including both analyses provides a more complete picture of the neuropathological effects.

#### Pairwise Contrasts and Posterior Diagnostics

For all models, we computed estimated marginal means (EMMs) using the emmeans R package to calculate effects of interest from the posterior distributions of the four subgroup means. This allowed for targeted post-hoc pairwise comparisons, such as comparing ZIKV vs. Control within females and separately within males. To facilitate interpretation and comparison across regions, the resulting posterior distributions for these contrasts (in mm³) were also standardized by dividing them by the pooled standard deviation of the corresponding brain region from the experimental data. This provided a standardized effect size (Cohen’s d) for each comparison, in addition to the estimate in natural units.

## ACKNOWLEDGMENTS

Authors would like to thank former Chahroudi and Raper laboratory members that assisted with data collection and processing; Winni Weng, Joseph Park, and Kristin Edwards. We also thank veterinary, animal care, behavioral management and Imaging Core staff at the Emory National Biomedical Research Center (ENBRC) for their excellent care of the animals and technical support throughout the study.

## AUTHOR CONTRIBUTIONS

A.C. and J.R. conceived and designed the study and acquired funding & supervised the project. Viral preparation, inoculations, viral load quantification, and cytokine/chemokine assays were performed and overseen by K.M., N.S., P.W., V.S., V.-V.E., M.S.S., and A.C. Neuroimaging data were acquired by Z.A.K.-B., R.A.T.M., R.L., J.A.-P., N.S., and M.M.S. and processed using the pipeline developed by R.V. and M.S. Behavioral testing and data collection/scoring were performed by K.L., A.v.S., S.F., M.A., R.R., and J.R. N.D. performed the statistical analyses, developed the analysis code, and created the figures. N.D. and J.R. wrote the original draft. All authors reviewed, edited, and approved the final manuscript.

## DATA AVAILABILITY

The datasets generated and analyzed in this study — including the plasma viral load, cytokine/chemokine, neurobehavioral (INAS), visual acuity, acute-stress, attachment, and 3-month neuroimaging (regional volume) data — are publicly available as CSV files in the ZIKV-longitudinal-study repository on GitHub (https://github.com/desai-nisarg/ZIKV-longitudinal-study). The source data underlying Figures 1–7 and the Supplementary Figures are included in that repository. Any further information required to reanalyze the data reported in this paper is available from the corresponding author (J.R.) upon reasonable request.

## CODE AVAILABILITY

All custom code used to process and analyze the data and to generate the results and figures reported in this study — including the Bayesian (brms) model scripts for the neurobehavioral, visual acuity, acute-stress, attachment, neuroimaging, cytokine, and viral-load (AUC) analyses — is publicly available in the ZIKV-longitudinal-study repository on GitHub (https://github.com/desai-nisarg/ZIKV-longitudinal-study).

