## Supplementary Materials for "Altered neurodevelopment, CSF volume, and emotional reactivity following postnatal Zika virus infection in infant macaques"

**Supplementary Table S1. Behavioral Ethogram for Acute Stressor and Attachment Assessments**

| **Category and Specific Behaviors** | **Measurement** | **Brief Descriptions** |
|---|---|---|
| Freeze[1] | Duration(sec) | Rigid, motionless posture except slight head movement |
| Hostile | Cumulative Frequency | |
| Threat | Frequency | Any of the following: open mouth (no teeth exposed), head-bobbing, or ear flapping |
| Cage aggression | Frequency | Vigorously slaps, shakes, or slams body against cage |
| Affiliative | Cumulative Frequency | |
| Coo | Frequency | Clear soft pitch and intensity, sounds like “ooooh” |
| Grunt | Frequency | Deep, muffled, low intensity, almost gurgling sound |
| Lipsmack | Frequency | Rapid movement of pursed lips, accompanied by a smacking sound |
| Anxiety | Cumulative Frequency | |
| Scratch | Frequency | Rapid scratch of body with hands or feet |
| Body shake | Frequency | Shake of the whole body or just head and shoulders region |
| Tooth grind | Frequency | Repetitive, audible rubbing of upper and lower teeth |
| Yawn | Frequency | Open mouth widely, exposing teeth |
| Self-directed Behaviors | Cumulative Duration | |
| Self-grooming | Duration | Use of hands or mouth to smooth or pick through fur |

| | | |
|---|---|---|
| Self-clasping[1] | Duration | Non-manipulatory enclosing or holding of<br>a limb or body part with arms |
| Other Self-directed | Duration | Sucking thumb, eye poke |
| Scream vocalizations | Frequency | High pitch, high intensity screech or loud chirp |
| Threat bark vocalizations | Frequency | Low pitch, high intensity, rasping, guttural |
| Fearful | Cumulative Frequency | |
| Withdrawal[1] | Frequency | Quick, jerky motion away from the stimulus object (jump back) |
| Grimace | Frequency | Refracted lips, exposed clenched teeth |
| Proximity[2] | Duration (seconds) | Infant enters a predetermined 2.5’ X 2.5’ area inside the cage immediately in front of the PHC or FC |
| Latency[2] | Duration (seconds) | Amount of time before the infant approaches the PHC after entering the testing cage |
| Attention Seeking[2] | Cumulative Frequency | |
| Reach[2] | Frequency | Subject stretches hand(s), arm(s) or leg(s) towards the PHC or FH. |
| Girn[2] | Frequency | Soft, low-frequency, nasal whine. Lips are slightly open or closed. |
| Gecker[2] | Frequency | Loud, staccato distress vocalization that infants produce when ignored or rejected by their mothers/caregiver. |
| Tantrum[2] | Frequency | Sustained and high intensity fit, typically includes body jerks or lowering/laying the body toward the cage floor. |

[1] Behaviors that were only observed during the human intruder paradigm.
[2] Behaviors that were only observed during the two-choice attachment discrimination assessment.

Visual Acuity Task:

#### Supplementary Table S2. Beta Regression Coefficients for % Looking

| Parameter | Estimate [89% HDI] | Credible |
|---|---|---|
| **Intercept** | **1.446 [-0.345, 2.989]** | **No** |
| **Group: Zika** | **0.216 [-0.28, 0.701]** | **No** |
| **Contrast: Medium** | **-0.143 [-1.617, 1.306]** | **No** |
| **Contrast: High** | **0.024 [-1.531, 1.55]** | **No** |
| **Age: 6Months** | **-0.023 [-1.409, 1.422]** | **No** |
| **Group: Zika x Contrast: Medium** | **-0.079 [-0.746, 0.582]** | **No** |
| **Group: Zika x Contrast: High** | **-0.614 [-1.307, 0.072]** | **No** |
| **Group: Zika x Age: 6Months** | **0.084 [-0.577, 0.742]** | **No** |
| **Contrast: Medium x Age: 6Months** | **0.071 [-1.825, 1.909]** | **No** |
| **Contrast: High x Age: 6Months** | **0.052 [-1.833, 1.921]** | **No** |
| **Group: Zika x Contrast: Medium x Age: 6Months** | **-0.272 [-1.189, 0.664]** | **No** |
| **Group: Zika x Contrast: High x Age: 6Months** | **0.412 [-0.527, 1.35]** | **No** |

**Supplementary Table S3. Group differences by Age and Contrast Level**

| Contrast | Contrast Level | Age Group | Estimate [89% HDI] | Credible |
|---|---|---|---|---|
| **Zika - Control** | **Low** | **4 Months** | **0.024 [-0.041, 0.112]** | **No** |
| **Zika - Control** | **Medium** | **4 Months** | **0.016 [-0.053, 0.107]** | **No** |
| **Zika - Control** | **High** | **4 Months** | **-0.056 [-0.163, 0.021]** | **No** |
| **Zika - Control** | **Low** | **6 Months** | **0.034 [-0.026, 0.131]** | **No** |
| **Zika - Control** | **Medium** | **6 Months** | **-0.0056 [-0.091, 0.072]** | **No** |
| **Zika - Control** | **High** | **6 Months** | **0.0095 [-0.056, 0.093]** | **No** |

Acute Stress Assessment:
All RM infants displayed graded species-typical responses across the conditions of the human intruder paradigm. As expected, freezing duration peaked in the Profile condition relative to both Alone (Alone - Profile median: -2.84; 89% HDI: [-3.87, -1.79]) and Stare (Profile - Stare median: 2.83; 89% HDI: [1.46, 4.11]), an elevation present within both control and ZIKV groups. Hostile behaviors increased from Alone to Profile (Alone - Profile; median: -1.00; 89% HDI: [-1.54, -0.45]) and again from Profile to Stare (median: -1.77; 89% HDI: [-2.26, -1.23]), a pattern that held within both control and ZIKV-infected RM infants. Anxious behaviors also increased with the increasing salience of the conditions (Alone - Profile; median: -2.25; 89% HDI: [-3.19, -1.33]; Profile - Stare; median: -2.75; 89% HDI: [-3.49, -2.00]; Alone - Stare; median: -5.01; 89% HDI: [-5.83, -4.16]), with these patterns mirrored within both groups. Self-directed behaviors were higher under greater provocation, with frequencies increasing from Alone to both Profile (Alone - Profile; median: -1.17; 89% HDI: [-1.91, -0.44]) and Stare (Alone - Stare; median: -1.59; 89% HDI: [-2.46, -0.79]), consistently in both ZIKV and control groups.

Other effects not relating to Zika status included a significant interaction between sex and condition in screams and threats toward the intruder. Females produced more screams than males overall (Female - Male median: 0.68; 89% HDI: [0.20, 1.18]), a difference present in both the Profile (median: 0.96; 89% HDI: [0.23, 1.67]) and Stare (median: 0.76; 89% HDI: [0.04, 1.49]) conditions, but not in the Alone condition. For threats toward the intruder, males exhibited more threats than females, but only in the Profile condition (Female - Male median: -1.18; 89% HDI: [-2.25, -0.10]). No other overall sex main effects or Sex × Condition interactions with 89% HDIs excluding zero were found for the remaining behavioral categories.

Attachment Assessment:

### Supplementary Table S4. Beta Regression Coefficients for Index of Preference

| Parameter | Estimate [89% HDI] | Credible |
|---|---|---|
| Intercept | 2.173 [1.461, 2.9] | Yes |
| Group: Zika | -0.244 [-1.004, 0.52] | No |
| Sex: Male | -0.295 [-1.059, 0.466] | No |
| Group: Zika x Sex: Male | -0.333 [-1.31, 0.66] | No |

### Supplementary Table S5: Attachment-assessment pairwise contrasts (ZIKV vs. combined control) stratified by sex, for all measured behaviors, under the weakly-informative (harmonized) prior specification. Estimates are posterior median differences with 89% highest-density intervals (log scale for the negative-binomial behaviors; latent logit scale for the beta-distributed index of preference); a contrast is credible when the interval excludes zero. Threat barks are reported separately from hostile behaviors, consistent with the acute-stress assessment and Supplementary Table S8

| Behavior | Sex | Estimate [89% HDI] | Credible |
|---|---|---|---|
| Affiliative (freq) | Female | 0.13 [-0.48, 0.71] | No |
| Affiliative (freq) | Male | -0.29 [-0.92, 0.38] | No |
| Anxious (freq) | Female | 0.24 [-0.84, 1.29] | No |
| Anxious (freq) | Male | -0.06 [-1.52, 1.21] | No |
| Attention-seeking (freq) | Female | -0.40 [-1.49, 0.69] | No |
| Attention-seeking (freq) | Male | -0.69 [-1.96, 0.72] | No |
| Displacement (freq) | Female | -0.29 [-1.36, 0.69] | No |
| Displacement (freq) | Male | -0.03 [-1.23, 1.13] | No |
| Falls (freq) | Female | -0.16 [-1.01, 0.67] | No |
| Falls (freq) | Male | -0.59 [-1.62, 0.48] | No |
| Fearful (freq) | Female | -0.11 [-1.51, 1.36] | No |
| Fearful (freq) | Male | 0.19 [-1.62, 1.91] | No |
| Hostile (freq) | Female | 0.17 [-0.92, 1.25] | No |
| Hostile (freq) | Male | 0.55 [-0.77, 1.83] | No |
| Index of preference | Female | -0.23 [-1.00, 0.54] | No |
| Index of preference | Male | -0.57 [-1.48, 0.29] | No |
| Scream (freq) | Female | -0.32 [-0.99, 0.35] | No |
| Scream (freq) | Male | -1.11 [-1.84, -0.33] | Yes |
| Self-directed (dur, s) | Female | -0.20 [-1.54, 1.22] | No |
| Self-directed (dur, s) | Male | 0.32 [-1.17, 2.02] | No |
| Threat barks (freq) | Female | -0.66 [-1.44, 0.15] | No |
| Threat barks (freq) | Male | -0.33 [-1.22, 0.57] | No |

Neuroimaging Assessment:

**Supplementary Table S6: Pairwise contrasts stratified by sex for all brain regions. Estimates are the posterior median of the standardized Zika minus control difference with the 89% credible interval.**

| Region | Contrast | Sex | Posterior median | Lower 89% CI | Upper 89% CI |
|---|---|---|---|---|---|
| Total CSF | Zika - Control | Female | 1.36 | 0.80 | 1.91 |
| Total CSF | Zika - Control | Male | 0.75 | 0.00 | 1.48 |
| Lat. Ventricle | Zika - Control | Female | -1.21 | -2.11 | -0.31 |
| Lat. Ventricle | Zika - Control | Male | 0.88 | -0.31 | 2.08 |
| Amygdala | Zika - Control | Female | 0.47 | -0.26 | 1.19 |
| Amygdala | Zika - Control | Male | 1.52 | 0.56 | 2.47 |
| Hippocampus | Zika - Control | Female | -0.47 | -1.15 | 0.20 |
| Hippocampus | Zika - Control | Male | 0.17 | -0.71 | 1.05 |
| RT Limbic GM WM | Zika - Control | Female | -0.81 | -1.30 | -0.33 |
| RT Limbic GM WM | Zika - Control | Male | 0.20 | -0.48 | 0.85 |
| LT Limbic GM WM | Zika - Control | Female | -0.78 | -1.30 | -0.28 |
| LT Limbic GM WM | Zika - Control | Male | 0.19 | -0.49 | 0.87 |
| RT Visual GM WM | Zika - Control | Female | -0.30 | -0.76 | 0.17 |
| RT Visual GM WM | Zika - Control | Male | -0.59 | -1.20 | 0.02 |
| LT Visual GM WM | Zika - Control | Female | -0.38 | -0.81 | 0.07 |
| LT Visual GM WM | Zika - Control | Male | -0.83 | -1.41 | -0.25 |
| RT Auditory GM WM | Zika - Control | Female | -0.34 | -0.72 | 0.05 |
| RT Auditory GM WM | Zika - Control | Male | -0.21 | -0.72 | 0.29 |
| LT Auditory GM WM | Zika - Control | Female | -0.40 | -0.77 | -0.02 |
| LT Auditory GM WM | Zika - Control | Male | -0.30 | -0.80 | 0.20 |
| RT Occipital GM WM | Zika - Control | Female | -0.76 | -1.21 | -0.31 |
| RT Occipital GM WM | Zika - Control | Male | -0.73 | -1.33 | -0.11 |
| LT Occipital GM WM | Zika - Control | Female | -0.70 | -1.13 | -0.26 |
| LT Occipital GM WM | Zika - Control | Male | -0.54 | -1.13 | 0.05 |

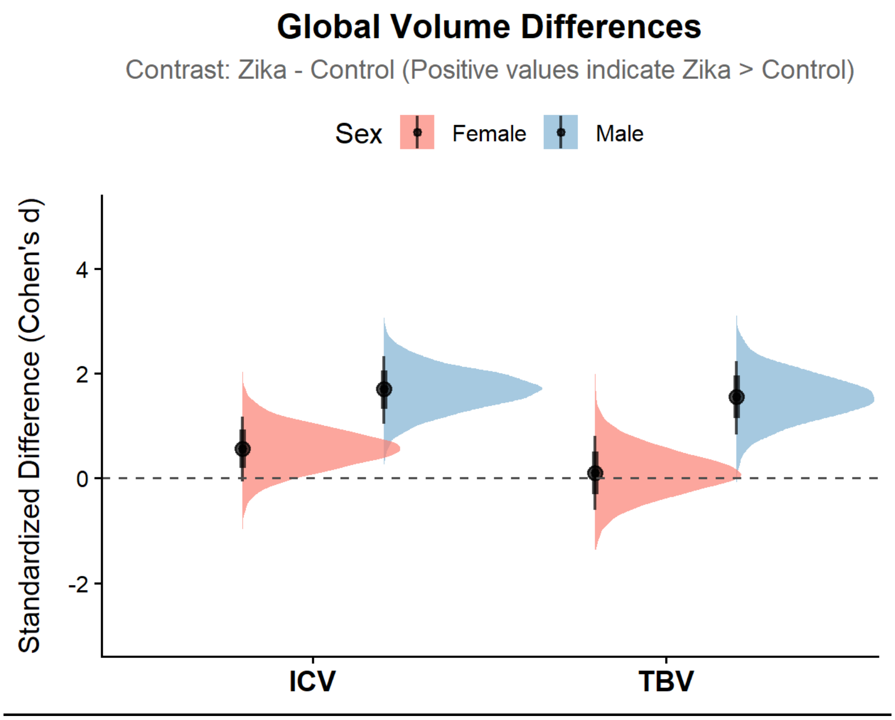


***Supplementary Figure S1. Sex dependent alterations in brain volume by ZIKV infection during infancy.*** *Sex-specific posterior distributions of the 'ZIKV - Control' contrast of the brain region volumes across multiple brain regions obtained from the 3-month MRI scans. This plot displays the standardized difference between the Zika and control groups in different brain regions. Estimates are derived from separate Bayesian linear regression models for each region. For each region on the x-axis, the contrast is visualized side-by-side for females and males, distinguished by color. Each half-eye plot illustrates the full probability density of the estimated difference. The internal black lines denote the 89% (wider) and 66% (narrower, thicker) credible intervals, with the posterior median marked by a black dot. The horizontal dashed line at y=0 represents no difference between the groups; a contrast is considered meaningful when its credible interval does not overlap this line.*

**Supplementary Table S7: Acute stress (Human Intruder) pairwise contrasts stratified by sex and condition, comparing ZIKV with each control and the two controls with each other. Estimates are posterior mean differences on the log-rate scale with 89% highest-density intervals; a contrast is credible when the interval excludes zero.**

| Behavior | Sex | Condition | Contrast | Estimate [89% HDI] | Rate ratio | Credible |
|---|---|---|---|---|---|---|
| Affiliative Vocals | Female | Alone | ZIKV - UIC | 0.49 [-0.41, 1.34] | 1.63 | No |
| Affiliative Vocals | Female | Alone | ZIKV - PIC | 0.23 [-0.81, 1.31] | 1.26 | No |
| Affiliative Vocals | Female | Alone | PIC - UIC | 0.24 [-0.70, 1.20] | 1.27 | No |
| Affiliative Vocals | Female | Profile | ZIKV - UIC | 0.30 [-0.79, 1.40] | 1.35 | No |
| Affiliative Vocals | Female | Profile | ZIKV - PIC | 0.03 [-1.33, 1.39] | 1.03 | No |
| Affiliative Vocals | Female | Profile | PIC - UIC | 0.25 [-0.98, 1.45] | 1.28 | No |
| Affiliative Vocals | Female | Stare | ZIKV - UIC | 0.32 [-0.71, 1.38] | 1.37 | No |
| Affiliative Vocals | Female | Stare | ZIKV - PIC | 0.24 [-1.00, 1.57] | 1.27 | No |
| Affiliative Vocals | Female | Stare | PIC - UIC | 0.07 [-1.10, 1.30] | 1.07 | No |
| Affiliative Vocals | Male | Alone | ZIKV - UIC | 0.93 [-0.14, 1.97] | 2.52 | No |
| Affiliative Vocals | Male | Alone | ZIKV - PIC | 0.13 [-1.08, 1.31] | 1.13 | No |
| Affiliative Vocals | Male | Alone | PIC - UIC | 0.81 [-0.37, 1.97] | 2.24 | No |
| Affiliative Vocals | Male | Profile | ZIKV - UIC | 0.80 [-0.50, 2.06] | 2.23 | No |
| Affiliative Vocals | Male | Profile | ZIKV - PIC | -0.25 [-1.57, 1.23] | 0.78 | No |
| Affiliative Vocals | Male | Profile | PIC - UIC | 1.06 [-0.33, 2.47] | 2.89 | No |
| Affiliative Vocals | Male | Stare | ZIKV - UIC | 0.61 [-0.64, 1.75] | 1.85 | No |
| Affiliative Vocals | Male | Stare | ZIKV - PIC | 0.26 [-1.26, 1.80] | 1.30 | No |
| Affiliative Vocals | Male | Stare | PIC - UIC | 0.38 [-1.18, 1.82] | 1.46 | No |
| Anxious Behavior | Female | Alone | ZIKV - UIC | -0.57 [-1.76, 0.62] | 0.57 | No |
| Anxious Behavior | Female | Alone | ZIKV - PIC | -0.83 [-2.49, 0.68] | 0.43 | No |
| Anxious Behavior | Female | Alone | PIC - UIC | 0.26 [-1.00, 1.50] | 1.30 | No |
| Anxious Behavior | Female | Profile | ZIKV - UIC | -0.44 [-1.96, 1.06] | 0.65 | No |
| Anxious Behavior | Female | Profile | ZIKV - PIC | -1.55 [-3.48, 0.38] | 0.21 | No |
| Anxious Behavior | Female | Profile | PIC - UIC | 1.10 [-0.47, 2.70] | 3.01 | No |
| Anxious Behavior | Female | Stare | ZIKV - UIC | -0.38 [-1.71, 1.12] | 0.68 | No |
| Anxious Behavior | Female | Stare | ZIKV - PIC | -0.89 [-2.61, 0.95] | 0.41 | No |
| Anxious Behavior | Female | Stare | PIC - UIC | 0.50 [-0.97, 2.02] | 1.65 | No |
| Anxious Behavior | Male | Alone | ZIKV - UIC | -0.40 [-1.95, 1.11] | 0.67 | No |
| Anxious Behavior | Male | Alone | ZIKV - PIC | -0.64 [-2.63, 1.37] | 0.53 | No |
| Anxious Behavior | Male | Alone | PIC - UIC | 0.22 [-1.52, 1.79] | 1.25 | No |
| Anxious Behavior | Male | Profile | ZIKV - UIC | 0.34 [-1.46, 2.13] | 1.41 | No |
| Anxious Behavior | Male | Profile | ZIKV - PIC | -0.84 [-3.08, 1.31] | 0.43 | No |
| Anxious Behavior | Male | Profile | PIC - UIC | 1.18 [-0.91, 3.12] | 3.26 | No |
| Anxious Behavior | Male | Stare | ZIKV - UIC | -0.25 [-1.93, 1.45] | 0.78 | No |
| Anxious Behavior | Male | Stare | ZIKV - PIC | -0.57 [-2.75, 1.49] | 0.57 | No |
| Anxious Behavior | Male | Stare | PIC - UIC | 0.32 [-1.61, 2.31] | 1.38 | No |
| Fearful Behavior | Female | Alone | ZIKV - UIC | -0.39 [-1.80, 1.17] | 0.67 | No |
| Fearful Behavior | Female | Alone | ZIKV - PIC | -0.44 [-2.49, 1.52] | 0.64 | No |
| Fearful Behavior | Female | Alone | PIC - UIC | 0.04 [-1.42, 1.53] | 1.04 | No |
| Fearful Behavior | Female | Profile | ZIKV - UIC | -0.66 [-2.68, 1.34] | 0.52 | No |
| Fearful Behavior | Female | Profile | ZIKV - PIC | -1.32 [-4.14, 1.33] | 0.27 | No |

| | | | | | | |
|---|---|---|---|---|---|---|
| Fearful Behavior | Female | Profile | PIC - UIC | 0.68 [-1.35, 2.72] | 1.97 | No |
| Fearful Behavior | Female | Stare | ZIKV - UIC | 0.08 [-1.84, 2.00] | 1.08 | No |
| Fearful Behavior | Female | Stare | ZIKV - PIC | 0.14 [-2.56, 2.74] | 1.15 | No |
| Fearful Behavior | Female | Stare | PIC - UIC | -0.03 [-1.95, 2.02] | 0.97 | No |
| Fearful Behavior | Male | Alone | ZIKV - UIC | -0.61 [-2.65, 1.37] | 0.54 | No |
| Fearful Behavior | Male | Alone | ZIKV - PIC | -0.78 [-3.52, 1.92] | 0.46 | No |
| Fearful Behavior | Male | Alone | PIC - UIC | 0.17 [-1.87, 2.18] | 1.18 | No |
| Fearful Behavior | Male | Profile | ZIKV - UIC | -0.73 [-3.48, 1.90] | 0.48 | No |
| Fearful Behavior | Male | Profile | ZIKV - PIC | -2.26 [-5.78, 1.29] | 0.10 | No |
| Fearful Behavior | Male | Profile | PIC - UIC | 1.49 [-1.35, 4.16] | 4.42 | No |
| Fearful Behavior | Male | Stare | ZIKV - UIC | -0.11 [-2.73, 2.51] | 0.89 | No |
| Fearful Behavior | Male | Stare | ZIKV - PIC | 0.10 [-3.48, 3.65] | 1.10 | No |
| Fearful Behavior | Male | Stare | PIC - UIC | -0.20 [-2.96, 2.58] | 0.82 | No |
| Freezing | Female | Alone | ZIKV - UIC | 0.05 [-1.27, 1.41] | 1.05 | No |
| Freezing | Female | Alone | ZIKV - PIC | 0.46 [-1.38, 2.34] | 1.58 | No |
| Freezing | Female | Alone | PIC - UIC | -0.41 [-1.86, 1.00] | 0.67 | No |
| Freezing | Female | Profile | ZIKV - UIC | 0.33 [-1.39, 2.06] | 1.39 | No |
| Freezing | Female | Profile | ZIKV - PIC | 0.27 [-2.17, 2.48] | 1.31 | No |
| Freezing | Female | Profile | PIC - UIC | 0.05 [-1.74, 1.89] | 1.05 | No |
| Freezing | Female | Stare | ZIKV - UIC | 0.79 [-1.02, 2.56] | 2.20 | No |
| Freezing | Female | Stare | ZIKV - PIC | 1.58 [-1.24, 4.13] | 4.84 | No |
| Freezing | Female | Stare | PIC - UIC | -0.80 [-2.86, 1.30] | 0.45 | No |
| Freezing | Male | Alone | ZIKV - UIC | -0.40 [-2.26, 1.34] | 0.67 | No |
| Freezing | Male | Alone | ZIKV - PIC | 0.17 [-2.29, 2.75] | 1.18 | No |
| Freezing | Male | Alone | PIC - UIC | -0.60 [-2.55, 1.29] | 0.55 | No |
| Freezing | Male | Profile | ZIKV - UIC | 0.02 [-2.19, 2.18] | 1.02 | No |
| Freezing | Male | Profile | ZIKV - PIC | -0.10 [-2.91, 2.80] | 0.91 | No |
| Freezing | Male | Profile | PIC - UIC | 0.09 [-2.12, 2.55] | 1.10 | No |
| Freezing | Male | Stare | ZIKV - UIC | 0.16 [-2.15, 2.57] | 1.18 | No |
| Freezing | Male | Stare | ZIKV - PIC | 1.38 [-2.38, 4.78] | 3.98 | No |
| Freezing | Male | Stare | PIC - UIC | -1.20 [-4.02, 1.58] | 0.30 | No |
| Hostile Behavior | Female | Alone | ZIKV - UIC | 1.58 [0.65, 2.54] | 4.86 | Yes |
| Hostile Behavior | Female | Alone | ZIKV - PIC | 1.81 [0.64, 3.03] | 6.11 | Yes |
| Hostile Behavior | Female | Alone | PIC - UIC | -0.24 [-1.30, 0.83] | 0.79 | No |
| Hostile Behavior | Female | Profile | ZIKV - UIC | 1.69 [0.59, 2.79] | 5.41 | Yes |
| Hostile Behavior | Female | Profile | ZIKV - PIC | 1.53 [0.17, 2.89] | 4.63 | Yes |
| Hostile Behavior | Female | Profile | PIC - UIC | 0.16 [-1.15, 1.34] | 1.17 | No |
| Hostile Behavior | Female | Stare | ZIKV - UIC | 0.88 [-0.23, 1.93] | 2.41 | No |
| Hostile Behavior | Female | Stare | ZIKV - PIC | -0.47 [-1.66, 0.80] | 0.63 | No |
| Hostile Behavior | Female | Stare | PIC - UIC | 1.36 [0.21, 2.48] | 3.89 | Yes |
| Hostile Behavior | Male | Alone | ZIKV - UIC | 1.70 [0.52, 2.78] | 5.48 | Yes |
| Hostile Behavior | Male | Alone | ZIKV - PIC | 2.29 [0.82, 3.89] | 9.87 | Yes |
| Hostile Behavior | Male | Alone | PIC - UIC | -0.59 [-2.00, 0.75] | 0.55 | No |
| Hostile Behavior | Male | Profile | ZIKV - UIC | 1.62 [0.32, 2.86] | 5.03 | Yes |
| Hostile Behavior | Male | Profile | ZIKV - PIC | 1.25 [-0.12, 2.76] | 3.50 | No |
| Hostile Behavior | Male | Profile | PIC - UIC | 0.36 [-1.08, 1.82] | 1.43 | No |
| Hostile Behavior | Male | Stare | ZIKV - UIC | 0.53 [-0.67, 1.81] | 1.70 | No |
| Hostile Behavior | Male | Stare | ZIKV - PIC | -0.39 [-1.85, 1.02] | 0.68 | No |

| | | | | | | |
|---|---|---|---|---|---|---|
| Hostile Behavior | Male | Stare | PIC - UIC | 0.94 [-0.45, 2.31] | 2.55 | No |
| Screams | Female | Alone | ZIKV - UIC | -0.10 [-0.85, 0.67] | 0.91 | No |
| Screams | Female | Alone | ZIKV - PIC | -0.05 [-0.96, 0.90] | 0.95 | No |
| Screams | Female | Alone | PIC - UIC | -0.05 [-0.85, 0.85] | 0.96 | No |
| Screams | Female | Profile | ZIKV - UIC | -0.49 [-1.43, 0.48] | 0.61 | No |
| Screams | Female | Profile | ZIKV - PIC | -0.64 [-1.74, 0.50] | 0.53 | No |
| Screams | Female | Profile | PIC - UIC | 0.15 [-0.94, 1.16] | 1.16 | No |
| Screams | Female | Stare | ZIKV - UIC | -0.54 [-1.51, 0.40] | 0.58 | No |
| Screams | Female | Stare | ZIKV - PIC | -0.34 [-1.52, 0.80] | 0.71 | No |
| Screams | Female | Stare | PIC - UIC | -0.20 [-1.24, 0.93] | 0.82 | No |
| Screams | Male | Alone | ZIKV - UIC | -0.13 [-1.01, 0.79] | 0.88 | No |
| Screams | Male | Alone | ZIKV - PIC | -0.12 [-1.24, 0.90] | 0.89 | No |
| Screams | Male | Alone | PIC - UIC | -0.002 [-1.00, 1.05] | 1.00 | No |
| Screams | Male | Profile | ZIKV - UIC | -0.58 [-1.71, 0.56] | 0.56 | No |
| Screams | Male | Profile | ZIKV - PIC | -0.64 [-1.89, 0.67] | 0.52 | No |
| Screams | Male | Profile | PIC - UIC | 0.08 [-1.18, 1.44] | 1.08 | No |
| Screams | Male | Stare | ZIKV - UIC | -0.87 [-2.05, 0.26] | 0.42 | No |
| Screams | Male | Stare | ZIKV - PIC | -0.76 [-2.06, 0.53] | 0.47 | No |
| Screams | Male | Stare | PIC - UIC | -0.12 [-1.36, 1.14] | 0.89 | No |
| Self Directed Behavior | Female | Alone | ZIKV - UIC | -0.53 [-1.88, 0.90] | 0.59 | No |
| Self Directed Behavior | Female | Alone | ZIKV - PIC | -0.37 [-2.18, 1.56] | 0.69 | No |
| Self Directed Behavior | Female | Alone | PIC - UIC | -0.16 [-1.49, 1.36] | 0.86 | No |
| Self Directed Behavior | Female | Profile | ZIKV - UIC | -0.83 [-2.56, 0.82] | 0.43 | No |
| Self Directed Behavior | Female | Profile | ZIKV - PIC | -0.18 [-2.49, 1.96] | 0.83 | No |
| Self Directed Behavior | Female | Profile | PIC - UIC | -0.65 [-2.42, 1.14] | 0.52 | No |
| Self Directed Behavior | Female | Stare | ZIKV - UIC | -0.49 [-2.21, 1.19] | 0.61 | No |
| Self Directed Behavior | Female | Stare | ZIKV - PIC | -0.91 [-3.13, 1.35] | 0.40 | No |
| Self Directed Behavior | Female | Stare | PIC - UIC | 0.39 [-1.40, 2.21] | 1.48 | No |
| Self Directed Behavior | Male | Alone | ZIKV - UIC | -0.26 [-2.09, 1.53] | 0.77 | No |
| Self Directed Behavior | Male | Alone | ZIKV - PIC | 0.38 [-2.09, 2.84] | 1.46 | No |
| Self Directed Behavior | Male | Alone | PIC - UIC | -0.59 [-2.58, 1.38] | 0.55 | No |
| Self Directed Behavior | Male | Profile | ZIKV - UIC | -1.20 [-3.46, 0.94] | 0.30 | No |
| Self Directed Behavior | Male | Profile | ZIKV - PIC | 0.77 [-2.23, 3.94] | 2.16 | No |
| Self Directed Behavior | Male | Profile | PIC - UIC | -1.98 [-4.50, 0.62] | 0.14 | No |
| Self Directed Behavior | Male | Stare | ZIKV - UIC | -0.90 [-3.14, 1.32] | 0.41 | No |
| Self Directed Behavior | Male | Stare | ZIKV - PIC | -2.14 [-4.98, 0.86] | 0.12 | No |
| Self Directed Behavior | Male | Stare | PIC - UIC | 1.24 [-1.22, 3.68] | 3.45 | No |
| Threat Barks | Female | Alone | ZIKV - UIC | -0.66 [-1.45, 0.20] | 0.51 | No |
| Threat Barks | Female | Alone | ZIKV - PIC | -0.77 [-1.73, 0.26] | 0.47 | No |
| Threat Barks | Female | Alone | PIC - UIC | 0.10 [-0.83, 0.97] | 1.11 | No |
| Threat Barks | Female | Profile | ZIKV - UIC | -1.46 [-2.50, -0.47] | 0.23 | Yes |
| Threat Barks | Female | Profile | ZIKV - PIC | -1.14 [-2.43, 0.05] | 0.32 | No |
| Threat Barks | Female | Profile | PIC - UIC | -0.32 [-1.45, 0.79] | 0.73 | No |
| Threat Barks | Female | Stare | ZIKV - UIC | -1.09 [-2.10, -0.07] | 0.34 | Yes |
| Threat Barks | Female | Stare | ZIKV - PIC | -1.59 [-2.67, -0.37] | 0.20 | Yes |
| Threat Barks | Female | Stare | PIC - UIC | 0.50 [-0.60, 1.56] | 1.65 | No |
| Threat Barks | Male | Alone | ZIKV - UIC | -0.40 [-1.40, 0.57] | 0.67 | No |
| Threat Barks | Male | Alone | ZIKV - PIC | 0.15 [-1.02, 1.28] | 1.17 | No |

| | | | | | | |
|---|---|---|---|---|---|---|
| Threat Barks | Male | Alone | PIC - UIC | -0.54 [-1.68, 0.55] | 0.58 | No |
| Threat Barks | Male | Profile | ZIKV - UIC | -0.92 [-2.10, 0.24] | 0.40 | No |
| Threat Barks | Male | Profile | ZIKV - PIC | 0.07 [-1.25, 1.34] | 1.08 | No |
| Threat Barks | Male | Profile | PIC - UIC | -0.99 [-2.32, 0.34] | 0.37 | No |
| Threat Barks | Male | Stare | ZIKV - UIC | -0.62 [-1.75, 0.45] | 0.54 | No |
| Threat Barks | Male | Stare | ZIKV - PIC | 0.54 [-0.85, 1.87] | 1.71 | No |
| Threat Barks | Male | Stare | PIC - UIC | -1.16 [-2.50, 0.19] | 0.31 | No |

**Supplementary Table S8: Attachment assessment pairwise contrasts stratified by sex, comparing ZIKV with each control and the two controls with each other, under the weakly-informative prior specification. Estimates are posterior mean differences on the log scale with 89% highest-density intervals; the multiplicative effect is exp(estimate); a contrast is credible when the interval excludes zero. Threat barks, previously combined with hostile behaviors into a single hostile/defensive composite, are here reported separately.**

| Behavior | Sex | Contrast | Estimate [89% HDI] | Mult. effect | Credible |
|---|---|---|---|---|---|
| Affiliative (freq) | Female | ZIKV - UIC | 0.17 [-0.50, 0.84] | 1.19 | No |
| | Female | ZIKV - PIC | 0.09 [-0.71, 0.87] | 1.09 | No |
| | Female | PIC - UIC | 0.07 [-0.66, 0.85] | 1.08 | No |
| | Male | ZIKV - UIC | -0.18 [-0.93, 0.62] | 0.84 | No |
| | Male | ZIKV - PIC | -0.46 [-1.32, 0.39] | 0.63 | No |
| | Male | PIC - UIC | 0.30 [-0.55, 1.17] | 1.36 | No |
| Anxious (freq) | Female | ZIKV - UIC | 0.15 [-0.99, 1.28] | 1.16 | No |
| | Female | ZIKV - PIC | 0.32 [-1.13, 1.75] | 1.37 | No |
| | Female | PIC - UIC | -0.15 [-1.36, 1.06] | 0.86 | No |
| | Male | ZIKV - UIC | -0.17 [-1.61, 1.29] | 0.84 | No |
| | Male | ZIKV - PIC | 0.33 [-1.63, 2.27] | 1.39 | No |
| | Male | PIC - UIC | -0.49 [-2.14, 1.22] | 0.61 | No |
| Attention-seeking (freq) | Female | ZIKV - UIC | -0.38 [-1.44, 0.89] | 0.68 | No |
| | Female | ZIKV - PIC | -0.58 [-1.97, 0.83] | 0.56 | No |
| | Female | PIC - UIC | 0.21 [-0.98, 1.43] | 1.23 | No |
| | Male | ZIKV - UIC | -0.69 [-2.12, 0.76] | 0.50 | No |
| | Male | ZIKV - PIC | -0.50 [-2.55, 1.40] | 0.60 | No |
| | Male | PIC - UIC | -0.18 [-1.86, 1.52] | 0.84 | No |
| Displacement (freq) | Female | ZIKV - UIC | -0.30 [-1.37, 0.78] | 0.74 | No |
| | Female | ZIKV - PIC | -0.47 [-1.72, 0.83] | 0.62 | No |
| | Female | PIC - UIC | 0.16 [-0.94, 1.31] | 1.17 | No |
| | Male | ZIKV - UIC | -0.07 [-1.27, 1.16] | 0.93 | No |
| | Male | ZIKV - PIC | 0.58 [-1.25, 2.40] | 1.78 | No |
| | Male | PIC - UIC | -0.65 [-2.34, 1.01] | 0.52 | No |
| Falls (freq) | Female | ZIKV - UIC | -0.31 [-1.19, 0.58] | 0.74 | No |
| | Female | ZIKV - PIC | 0.19 [-0.98, 1.38] | 1.21 | No |
| | Female | PIC - UIC | -0.50 [-1.50, 0.54] | 0.60 | No |
| | Male | ZIKV - UIC | -0.76 [-1.87, 0.31] | 0.47 | No |

| | | | | | |
|---|---|---|---|---|---|
| | Male | ZIKV - PIC | -0.25 [-1.54, 1.20] | 0.78 | No |
| | Male | PIC - UIC | -0.50 [-1.75, 0.73] | 0.61 | No |
| Fearful (freq) | Female | ZIKV - UIC | -0.13 [-1.64, 1.23] | 0.88 | No |
| | Female | ZIKV - PIC | -0.10 [-2.08, 1.85] | 0.91 | No |
| | Female | PIC - UIC | -0.02 [-1.40, 1.45] | 0.98 | No |
| | Male | ZIKV - UIC | 0.16 [-1.63, 1.96] | 1.17 | No |
| | Male | ZIKV - PIC | 0.14 [-2.35, 2.57] | 1.15 | No |
| | Male | PIC - UIC | 0.05 [-1.87, 2.03] | 1.05 | No |
| Hostile (freq) | Female | ZIKV - UIC | 0.13 [-0.97, 1.28] | 1.14 | No |
| | Female | ZIKV - PIC | 0.08 [-1.25, 1.47] | 1.09 | No |
| | Female | PIC - UIC | 0.04 [-1.13, 1.20] | 1.04 | No |
| | Male | ZIKV - UIC | 0.46 [-0.89, 1.75] | 1.58 | No |
| | Male | ZIKV - PIC | 1.00 [-0.99, 2.91] | 2.71 | No |
| | Male | PIC - UIC | -0.53 [-2.26, 1.15] | 0.59 | No |
| Index of preference | Female | ZIKV - UIC | 0.02 [-0.83, 0.87] | 1.02 | No |
| | Female | ZIKV - PIC | -0.59 [-1.54, 0.35] | 0.55 | No |
| | Female | PIC - UIC | 0.62 [-0.32, 1.52] | 1.86 | No |
| | Male | ZIKV - UIC | -0.31 [-1.25, 0.67] | 0.74 | No |
| | Male | ZIKV - PIC | -0.87 [-2.01, 0.27] | 0.42 | No |
| | Male | PIC - UIC | 0.56 [-0.50, 1.67] | 1.75 | No |
| Scream (freq) | Female | ZIKV - UIC | -0.28 [-0.95, 0.44] | 0.76 | No |
| | Female | ZIKV - PIC | -0.22 [-1.11, 0.63] | 0.81 | No |
| | Female | PIC - UIC | -0.06 [-0.87, 0.80] | 0.94 | No |
| | Male | ZIKV - UIC | -0.85 [-1.72, -0.003] | 0.43 | Yes |
| | Male | ZIKV - PIC | -1.44 [-2.34, -0.51] | 0.24 | Yes |
| | Male | PIC - UIC | 0.59 [-0.37, 1.51] | 1.80 | No |
| Self-directed (dur, s) | Female | ZIKV - UIC | -0.23 [-1.56, 1.16] | 0.79 | No |
| | Female | ZIKV - PIC | -0.34 [-2.27, 1.39] | 0.71 | No |
| | Female | PIC - UIC | 0.11 [-1.34, 1.47] | 1.12 | No |
| | Male | ZIKV - UIC | 0.29 [-1.33, 1.88] | 1.34 | No |
| | Male | ZIKV - PIC | 0.42 [-2.04, 2.60] | 1.53 | No |
| | Male | PIC - UIC | -0.12 [-2.10, 1.84] | 0.88 | No |
| Threat barks (freq) | Female | ZIKV - UIC | -0.63 [-1.45, 0.22] | 0.53 | No |
| | Female | ZIKV - PIC | -0.83 [-1.83, 0.14] | 0.43 | No |
| | Female | PIC - UIC | 0.20 [-0.68, 1.09] | 1.22 | No |
| | Male | ZIKV - UIC | -0.42 [-1.41, 0.56] | 0.66 | No |
| | Male | ZIKV - PIC | -0.17 [-1.35, 1.06] | 0.84 | No |
| | Male | PIC - UIC | -0.24 [-1.43, 0.92] | 0.78 | No |

**Supplementary Table S9: Visual-acuity pairwise contrasts (overall and contrast-level by age), comparing ZIKV with each control and the two controls with each other. Estimates are differences in the proportion of time looking with 89% highest-density intervals; a contrast is credible when the interval excludes zero.**

| Model | Contrast level | Age | Contrast | Estimate [89% HDI] | Credible |
|---|---|---|---|---|---|
| Global (overall) | | | ZIKV - UIC | 0.04 [-0.01, 0.09] | No |
| Global (overall) | | | ZIKV - PIC | -0.01 [-0.06, 0.03] | No |
| Global (overall) | | | PIC - UIC | 0.05 [0.004, 0.12] | Yes |
| Interaction (contrast x age) | High | 4 Months | ZIKV - UIC | -0.004 [-0.12, 0.09] | No |
| Interaction (contrast x age) | High | 4 Months | ZIKV - PIC | -0.10 [-0.22, 0.000] | No |
| Interaction (contrast x age) | High | 4 Months | PIC - UIC | 0.09 [-0.002, 0.23] | No |
| Interaction (contrast x age) | Low | 4 Months | ZIKV - UIC | 0.04 [-0.04, 0.14] | No |
| Interaction (contrast x age) | Low | 4 Months | ZIKV - PIC | 0.02 [-0.07, 0.13] | No |
| Interaction (contrast x age) | Low | 4 Months | PIC - UIC | 0.01 [-0.09, 0.12] | No |
| Interaction (contrast x age) | Medium | 4 Months | ZIKV - UIC | 0.10 [-0.01, 0.22] | No |
| Interaction (contrast x age) | Medium | 4 Months | ZIKV - PIC | -0.03 [-0.14, 0.05] | No |
| Interaction (contrast x age) | Medium | 4 Months | PIC - UIC | 0.14 [0.01, 0.27] | Yes |
| Interaction (contrast x age) | High | 6 Months | ZIKV - UIC | -0.003 [-0.09, 0.08] | No |
| Interaction (contrast x age) | High | 6 Months | ZIKV - PIC | 0.03 [-0.05, 0.15] | No |
| Interaction (contrast x age) | High | 6 Months | PIC - UIC | -0.04 [-0.16, 0.06] | No |
| Interaction (contrast x age) | Low | 6 Months | ZIKV - UIC | 0.08 [-0.01, 0.20] | No |
| Interaction (contrast x age) | Low | 6 Months | ZIKV - PIC | -0.002 [-0.08, 0.08] | No |
| Interaction (contrast x age) | Low | 6 Months | PIC - UIC | 0.08 [-0.01, 0.22] | No |
| Interaction (contrast x age) | Medium | 6 Months | ZIKV - UIC | -0.003 [-0.10, 0.10] | No |
| Interaction (contrast x age) | Medium | 6 Months | ZIKV - PIC | -0.01 [-0.11, 0.09] | No |
| Interaction (contrast x age) | Medium | 6 Months | PIC - UIC | 0.003 [-0.10, 0.12] | No |

### Supplementary Table S10: Neuroimaging pairwise contrasts stratified by sex, comparing ZIKV with each control and the two controls with each other, for all regional volumes. Estimates are standardized mean differences (Cohen's d) with 89% highest-density intervals; prob(>0) is the posterior probability that the difference exceeds zero; a contrast is credible when the interval excludes zero.

| Region | Sex | Contrast | Estimate [89% HDI] | Prob(>0) | Credible |
|---|---|---|---|---|---|
| Amygdala | Female | ZIKV - UIC | 0.24 [-0.68, 1.15] | 0.67 | No |
| Amygdala | Female | ZIKV - PIC | 0.62 [-0.31, 1.52] | 0.86 | No |
| Amygdala | Female | PIC - UIC | -0.38 [-1.40, 0.65] | 0.27 | No |
| Amygdala | Male | ZIKV - UIC | 1.26 [0.18, 2.26] | 0.97 | Yes |
| Amygdala | Male | ZIKV - PIC | 1.27 [0.11, 2.37] | 0.96 | Yes |
| Amygdala | Male | PIC - UIC | -0.02 [-0.95, 0.93] | 0.49 | No |
| Hippocampus | Female | ZIKV - UIC | -0.46 [-1.28, 0.35] | 0.18 | No |
| Hippocampus | Female | ZIKV - PIC | -0.51 [-1.33, 0.29] | 0.15 | No |
| Hippocampus | Female | PIC - UIC | 0.05 [-0.86, 0.97] | 0.53 | No |
| Hippocampus | Male | ZIKV - UIC | -0.06 [-1.04, 0.89] | 0.46 | No |
| Hippocampus | Male | ZIKV - PIC | 0.36 [-0.71, 1.39] | 0.72 | No |
| Hippocampus | Male | PIC - UIC | -0.43 [-1.30, 0.46] | 0.21 | No |
| L. Occipital | Female | ZIKV - UIC | -0.43 [-0.94, 0.10] | 0.09 | No |
| L. Occipital | Female | ZIKV - PIC | -0.91 [-1.42, -0.40] | 0.01 | Yes |
| L. Occipital | Female | PIC - UIC | 0.49 [-0.08, 1.05] | 0.92 | No |

| | | | | | |
|---|---|---|---|---|---|
| L. Occipital | Male | ZIKV - UIC | -0.44 [-1.06, 0.16] | 0.12 | No |
| L. Occipital | Male | ZIKV - PIC | -0.70 [-1.37, -0.02] | 0.05 | Yes |
| L. Occipital | Male | PIC - UIC | 0.27 [-0.31, 0.84] | 0.78 | No |
| L. Temporal-Auditory | Female | ZIKV - UIC | -0.24 [-0.70, 0.21] | 0.18 | No |
| L. Temporal-Auditory | Female | ZIKV - PIC | -0.53 [-0.97, -0.08] | 0.03 | Yes |
| L. Temporal-Auditory | Female | PIC - UIC | 0.29 [-0.20, 0.78] | 0.83 | No |
| L. Temporal-Auditory | Male | ZIKV - UIC | -0.16 [-0.68, 0.37] | 0.30 | No |
| L. Temporal-Auditory | Male | ZIKV - PIC | -0.47 [-1.04, 0.12] | 0.10 | No |
| L. Temporal-Auditory | Male | PIC - UIC | 0.31 [-0.19, 0.81] | 0.84 | No |
| L. Temporal-Limbic | Female | ZIKV - UIC | -0.64 [-1.29, -0.004] | 0.05 | Yes |
| L. Temporal-Limbic | Female | ZIKV - PIC | -0.98 [-1.62, -0.35] | 0.01 | Yes |
| L. Temporal-Limbic | Female | PIC - UIC | 0.34 [-0.37, 1.05] | 0.79 | No |
| L. Temporal-Limbic | Male | ZIKV - UIC | 0.05 [-0.73, 0.79] | 0.55 | No |
| L. Temporal-Limbic | Male | ZIKV - PIC | 0.23 [-0.60, 1.03] | 0.68 | No |
| L. Temporal-Limbic | Male | PIC - UIC | -0.18 [-0.89, 0.52] | 0.33 | No |
| L. Temporal-Visual | Female | ZIKV - UIC | -0.35 [-0.88, 0.17] | 0.14 | No |
| L. Temporal-Visual | Female | ZIKV - PIC | -0.36 [-0.89, 0.17] | 0.14 | No |
| L. Temporal-Visual | Female | PIC - UIC | 0.01 [-0.58, 0.59] | 0.51 | No |
| L. Temporal-Visual | Male | ZIKV - UIC | -0.66 [-1.29, -0.02] | 0.05 | Yes |
| L. Temporal-Visual | Male | ZIKV - PIC | -1.04 [-1.72, -0.33] | 0.01 | Yes |
| L. Temporal-Visual | Male | PIC - UIC | 0.38 [-0.22, 0.97] | 0.85 | No |
| Lateral Ventricle | Female | ZIKV - UIC | -1.21 [-2.29, -0.14] | 0.04 | Yes |
| Lateral Ventricle | Female | ZIKV - PIC | -1.27 [-2.34, -0.21] | 0.03 | Yes |
| Lateral Ventricle | Female | PIC - UIC | 0.05 [-1.16, 1.24] | 0.53 | No |
| Lateral Ventricle | Male | ZIKV - UIC | 0.97 [-0.32, 2.26] | 0.89 | No |
| Lateral Ventricle | Male | ZIKV - PIC | 0.65 [-0.74, 2.02] | 0.78 | No |
| Lateral Ventricle | Male | PIC - UIC | 0.33 [-0.81, 1.45] | 0.68 | No |
| R. Occipital | Female | ZIKV - UIC | -0.48 [-0.99, 0.05] | 0.07 | No |
| R. Occipital | Female | ZIKV - PIC | -1.02 [-1.51, -0.50] | 0.00 | Yes |
| R. Occipital | Female | PIC - UIC | 0.54 [-0.04, 1.11] | 0.93 | No |
| R. Occipital | Male | ZIKV - UIC | -0.58 [-1.20, 0.03] | 0.06 | No |
| R. Occipital | Male | ZIKV - PIC | -0.96 [-1.62, -0.29] | 0.02 | Yes |
| R. Occipital | Male | PIC - UIC | 0.37 [-0.21, 0.94] | 0.85 | No |
| R. Temporal-Auditory | Female | ZIKV - UIC | -0.28 [-0.75, 0.19] | 0.16 | No |
| R. Temporal-Auditory | Female | ZIKV - PIC | -0.39 [-0.84, 0.06] | 0.08 | No |
| R. Temporal-Auditory | Female | PIC - UIC | 0.11 [-0.40, 0.61] | 0.64 | No |
| R. Temporal-Auditory | Male | ZIKV - UIC | -0.08 [-0.63, 0.46] | 0.41 | No |
| R. Temporal-Auditory | Male | ZIKV - PIC | -0.43 [-1.03, 0.18] | 0.12 | No |
| R. Temporal-Auditory | Male | PIC - UIC | 0.35 [-0.17, 0.85] | 0.87 | No |
| R. Temporal-Limbic | Female | ZIKV - UIC | -0.78 [-1.35, -0.22] | 0.01 | Yes |
| R. Temporal-Limbic | Female | ZIKV - PIC | -0.90 [-1.47, -0.34] | 0.01 | Yes |
| R. Temporal-Limbic | Female | PIC - UIC | 0.12 [-0.51, 0.75] | 0.62 | No |
| R. Temporal-Limbic | Male | ZIKV - UIC | -0.11 [-0.81, 0.56] | 0.40 | No |
| R. Temporal-Limbic | Male | ZIKV - PIC | 0.46 [-0.32, 1.19] | 0.84 | No |
| R. Temporal-Limbic | Male | PIC - UIC | -0.57 [-1.21, 0.09] | 0.08 | No |
| R. Temporal-Visual | Female | ZIKV - UIC | -0.28 [-0.82, 0.28] | 0.20 | No |
| R. Temporal-Visual | Female | ZIKV - PIC | -0.28 [-0.81, 0.27] | 0.20 | No |
| R. Temporal-Visual | Female | PIC - UIC | -0.01 [-0.61, 0.60] | 0.49 | No |

| | | | | | |
|---|---|---|---|---|---|
| R. Temporal-Visual | Male | ZIKV - UIC | -0.42 [-1.07, 0.22] | 0.14 | No |
| R. Temporal-Visual | Male | ZIKV - PIC | -0.79 [-1.48, -0.07] | 0.04 | Yes |
| R. Temporal-Visual | Male | PIC - UIC | 0.36 [-0.26, 0.97] | 0.83 | No |
| Total CSF | Female | ZIKV - UIC | 1.35 [0.67, 2.02] | 1.00 | Yes |
| Total CSF | Female | ZIKV - PIC | 1.20 [0.52, 1.85] | 1.00 | Yes |
| Total CSF | Female | PIC - UIC | 0.15 [-0.56, 0.85] | 0.64 | No |
| Total CSF | Male | ZIKV - UIC | 0.36 [-0.50, 1.17] | 0.76 | No |
| Total CSF | Male | ZIKV - PIC | 0.63 [-0.31, 1.52] | 0.87 | No |
| Total CSF | Male | PIC - UIC | -0.27 [-1.02, 0.50] | 0.28 | No |

**Supplementary Table S11: Group x sex interaction for each brain region, i.e. the difference in the ZIKV (versus combined control) effect between males and females, from the ICV-adjusted models (n = 24 scans; ZIKV 6 female / 6 male, control 6 female / 6 male). Values are the standardized difference (Cohen's d, computed as the male effect minus the female effect; positive = larger or less-reduced in males) with the 89% highest-density interval (HDI), the probability of direction (pd), and the probability that the difference is practically meaningful, P(|d| > 0.2). A higher pd indicates stronger evidence that the ZIKV effect differs by sex.**

| Region | Interaction d (male - female) [89% HDI] | pd | P(\|d\| > 0.2) |
|---|---|---|---|
| Lat. Ventricle | 2.09 [0.68, 3.42] | 0.99 | 0.99 |
| RT Limbic GM WM | 1.00 [0.22, 1.70] | 0.98 | 0.97 |
| LT Limbic GM WM | 0.98 [0.23, 1.74] | 0.98 | 0.96 |
| Amygdala | 1.05 [-0.003, 2.15] | 0.94 | 0.94 |
| Total CSF | -0.61 [-1.41, 0.24] | 0.88 | 0.84 |
| LT Visual GM WM | -0.46 [-1.08, 0.23] | 0.86 | 0.79 |
| Hippocampus | 0.65 [-0.40, 1.58] | 0.85 | 0.86 |
| RT Visual GM WM | -0.28 [-0.98, 0.38] | 0.76 | 0.71 |
| LT Occipital GM WM | 0.16 [-0.51, 0.83] | 0.65 | 0.65 |
| RT Auditory GM WM | 0.12 [-0.43, 0.70] | 0.64 | 0.59 |
| LT Auditory GM WM | 0.10 [-0.46, 0.66] | 0.62 | 0.56 |
| RT Occipital GM WM | 0.04 [-0.66, 0.69] | 0.53 | 0.62 |

**Supplementary Table S12: Neurobehavioral (INAS) treatment contrasts with age modelled as a discrete instead of continuous variable. Each domain was refit as Treatment by factor(age in weeks), so the contrast at every week is estimated locally rather than from a single age trend. Values are posterior median differences on the latent logit scale with the 89% highest-density interval (HDI) and the probability of direction (pd). ZIKV challenge and Poly-IC administration occurred at a median of 35 days of life (approximately 5 weeks of age); assessments at 2-4 weeks (Timing = Pre) precede challenge, and the week-5 assessment was collected 6 days post-infection (Post).**

| Domain | Age (wk) | Contrast | Median [89% HDI] | pd | Credible | Timing |
|---|---|---|---|---|---|---|
| State control | 2 | ZIKV - UIC | 0.46 [-0.46, 1.46] | 0.78 | No | Pre |

| | | | | | | |
|---|---|---|---|---|---|---|
| | 3 | ZIKV - UIC | 0.38 [-0.67, 1.43] | 0.72 | No | Pre |
| | 4 | ZIKV - UIC | 1.92 [0.81, 3.04] | 1.00 | Yes | Pre |
| | 5 | ZIKV - UIC | 1.40 [0.39, 2.56] | 0.98 | Yes | Post |
| | 6 | ZIKV - UIC | 1.62 [0.54, 2.70] | 0.99 | Yes | Post |
| | 7 | ZIKV - UIC | 2.11 [1.05, 3.19] | 1.00 | Yes | Post |
| | 8 | ZIKV - UIC | 3.10 [2.04, 4.35] | 1.00 | Yes | Post |
| | 9 | ZIKV - UIC | 1.54 [0.53, 2.63] | 0.99 | Yes | Post |
| | 2 | ZIKV - PIC | 0.72 [-0.36, 1.78] | 0.86 | No | Pre |
| | 3 | ZIKV - PIC | 0.48 [-0.63, 1.62] | 0.76 | No | Pre |
| | 4 | ZIKV - PIC | 0.19 [-0.87, 1.28] | 0.61 | No | Pre |
| | 5 | ZIKV - PIC | 1.79 [0.61, 3.01] | 0.99 | Yes | Post |
| | 6 | ZIKV - PIC | 1.99 [0.76, 3.12] | 0.99 | Yes | Post |
| | 7 | ZIKV - PIC | 2.45 [1.30, 3.63] | 1.00 | Yes | Post |
| | 8 | ZIKV - PIC | 4.49 [2.97, 6.02] | 1.00 | Yes | Post |
| | 9 | ZIKV - PIC | 1.90 [0.73, 2.98] | 0.99 | Yes | Post |
| | 2 | PIC - UIC | -0.26 [-1.32, 0.86] | 0.65 | No | Pre |
| | 3 | PIC - UIC | -0.09 [-1.27, 1.16] | 0.55 | No | Pre |
| | 4 | PIC - UIC | 1.75 [0.51, 3.00] | 0.99 | Yes | Pre |
| | 5 | PIC - UIC | -0.39 [-1.71, 0.96] | 0.68 | No | Post |
| | 6 | PIC - UIC | -0.37 [-1.62, 1.01] | 0.67 | No | Post |
| | 7 | PIC - UIC | -0.33 [-1.58, 1.01] | 0.67 | No | Post |
| | 8 | PIC - UIC | -1.38 [-3.11, 0.17] | 0.92 | No | Post |
| | 9 | PIC - UIC | -0.36 [-1.60, 0.86] | 0.68 | No | Post |
| Neuromotor | 2 | ZIKV - UIC | -0.40 [-0.82, 0.04] | 0.93 | No | Pre |
| | 3 | ZIKV - UIC | -0.55 [-1.07, -0.06] | 0.96 | Yes | Pre |
| | 4 | ZIKV - UIC | -0.96 [-1.46, -0.45] | 1.00 | Yes | Pre |
| | 5 | ZIKV - UIC | -0.83 [-1.28, -0.39] | 1.00 | Yes | Post |
| | 6 | ZIKV - UIC | -1.02 [-1.48, -0.59] | 1.00 | Yes | Post |
| | 7 | ZIKV - UIC | -1.06 [-1.65, -0.45] | 1.00 | Yes | Post |
| | 8 | ZIKV - UIC | -0.68 [-2.02, 0.80] | 0.78 | No | Post |
| | 9 | ZIKV - UIC | 0.51 [-0.96, 2.16] | 0.70 | No | Post |
| | 2 | ZIKV - PIC | -0.31 [-0.78, 0.15] | 0.86 | No | Pre |
| | 3 | ZIKV - PIC | -0.70 [-1.23, -0.21] | 0.99 | Yes | Pre |
| | 4 | ZIKV - PIC | -0.10 [-0.55, 0.34] | 0.64 | No | Pre |
| | 5 | ZIKV - PIC | -0.93 [-1.39, -0.48] | 1.00 | Yes | Post |
| | 6 | ZIKV - PIC | -1.13 [-1.59, -0.69] | 1.00 | Yes | Post |
| | 7 | ZIKV - PIC | -1.27 [-1.74, -0.80] | 1.00 | Yes | Post |
| | 8 | ZIKV - PIC | -0.86 [-2.60, 0.94] | 0.79 | No | Post |
| | 9 | ZIKV - PIC | -0.51 [-2.72, 1.63] | 0.65 | No | Post |
| | 2 | PIC - UIC | -0.08 [-0.59, 0.43] | 0.60 | No | Pre |
| | 3 | PIC - UIC | 0.15 [-0.48, 0.72] | 0.66 | No | Pre |
| | 4 | PIC - UIC | -0.86 [-1.41, -0.27] | 0.99 | Yes | Pre |
| | 5 | PIC - UIC | 0.11 [-0.46, 0.62] | 0.62 | No | Post |
| | 6 | PIC - UIC | 0.11 [-0.42, 0.67] | 0.63 | No | Post |
| | 7 | PIC - UIC | 0.22 [-0.47, 0.87] | 0.70 | No | Post |
| | 8 | PIC - UIC | 0.20 [-1.45, 1.88] | 0.58 | No | Post |
| | 9 | PIC - UIC | 1.01 [-0.78, 2.94] | 0.82 | No | Post |
| Orientation | 2 | ZIKV - UIC | 0.06 [-1.22, 1.46] | 0.53 | No | Pre |
| | 3 | ZIKV - UIC | 1.89 [0.15, 3.52] | 0.96 | Yes | Pre |
| | 4 | ZIKV - UIC | 0.62 [-0.98, 2.02] | 0.75 | No | Pre |
| | 5 | ZIKV - UIC | -0.14 [-1.59, 1.44] | 0.56 | No | Post |
| | 6 | ZIKV - UIC | -0.79 [-2.20, 0.69] | 0.83 | No | Post |

| | | | | | | |
|---|---|---|---|---|---|---|
| | 7 | ZIKV - UIC | -1.75 [-3.25, -0.26] | 0.97 | Yes | Post |
| | 8 | ZIKV - UIC | -2.70 [-4.23, -1.25] | 1.00 | Yes | Post |
| | 9 | ZIKV - UIC | -1.24 [-2.72, 0.18] | 0.92 | No | Post |
| | 2 | ZIKV - PIC | -0.29 [-1.91, 1.27] | 0.62 | No | Pre |
| | 3 | ZIKV - PIC | 1.02 [-1.01, 2.94] | 0.81 | No | Pre |
| | 4 | ZIKV - PIC | 0.40 [-1.22, 2.15] | 0.66 | No | Pre |
| | 5 | ZIKV - PIC | 0.02 [-1.69, 1.72] | 0.51 | No | Post |
| | 6 | ZIKV - PIC | -1.27 [-2.98, 0.38] | 0.90 | No | Post |
| | 7 | ZIKV - PIC | -2.09 [-3.82, -0.34] | 0.97 | Yes | Post |
| | 8 | ZIKV - PIC | -2.86 [-4.51, -1.11] | 1.00 | Yes | Post |
| | 9 | ZIKV - PIC | 0.73 [-0.84, 2.31] | 0.78 | No | Post |
| | 2 | PIC - UIC | 0.34 [-1.09, 1.80] | 0.65 | No | Pre |
| | 3 | PIC - UIC | 0.85 [-0.89, 2.57] | 0.79 | No | Pre |
| | 4 | PIC - UIC | 0.23 [-1.35, 1.88] | 0.59 | No | Pre |
| | 5 | PIC - UIC | -0.18 [-1.93, 1.47] | 0.57 | No | Post |
| | 6 | PIC - UIC | 0.50 [-1.18, 2.23] | 0.68 | No | Post |
| | 7 | PIC - UIC | 0.31 [-1.41, 2.15] | 0.62 | No | Post |
| | 8 | PIC - UIC | 0.18 [-1.61, 1.88] | 0.56 | No | Post |
| | 9 | PIC - UIC | -1.94 [-3.57, -0.42] | 0.98 | Yes | Post |
| Motor | 2 | ZIKV - UIC | -0.20 [-1.34, 1.08] | 0.60 | No | Pre |
| | 3 | ZIKV - UIC | -0.01 [-1.41, 1.43] | 0.50 | No | Pre |
| | 4 | ZIKV - UIC | 0.25 [-1.18, 1.66] | 0.61 | No | Pre |
| | 5 | ZIKV - UIC | -0.23 [-1.70, 1.22] | 0.60 | No | Post |
| | 6 | ZIKV - UIC | -0.15 [-1.58, 1.36] | 0.56 | No | Post |
| | 7 | ZIKV - UIC | 0.88 [-0.72, 2.45] | 0.82 | No | Post |
| | 8 | ZIKV - UIC | 1.08 [-0.41, 2.72] | 0.87 | No | Post |
| | 9 | ZIKV - UIC | -0.74 [-2.28, 0.67] | 0.79 | No | Post |
| | 2 | ZIKV - PIC | -0.61 [-2.11, 0.84] | 0.75 | No | Pre |
| | 3 | ZIKV - PIC | -0.26 [-1.91, 1.39] | 0.60 | No | Pre |
| | 4 | ZIKV - PIC | -0.28 [-1.94, 1.30] | 0.62 | No | Pre |
| | 5 | ZIKV - PIC | -0.55 [-2.25, 1.14] | 0.70 | No | Post |
| | 6 | ZIKV - PIC | 0.57 [-1.02, 2.25] | 0.72 | No | Post |
| | 7 | ZIKV - PIC | 0.84 [-0.96, 2.64] | 0.78 | No | Post |
| | 8 | ZIKV - PIC | -0.30 [-2.33, 1.70] | 0.60 | No | Post |
| | 9 | ZIKV - PIC | -2.82 [-4.87, -0.71] | 0.99 | Yes | Post |
| | 2 | PIC - UIC | 0.42 [-0.99, 1.78] | 0.70 | No | Pre |
| | 3 | PIC - UIC | 0.23 [-1.28, 2.05] | 0.59 | No | Pre |
| | 4 | PIC - UIC | 0.56 [-1.10, 2.17] | 0.71 | No | Pre |
| | 5 | PIC - UIC | 0.31 [-1.36, 2.05] | 0.61 | No | Post |
| | 6 | PIC - UIC | -0.72 [-2.56, 0.79] | 0.76 | No | Post |
| | 7 | PIC - UIC | 0.04 [-1.71, 1.75] | 0.52 | No | Post |
| | 8 | PIC - UIC | 1.39 [-0.60, 3.19] | 0.88 | No | Post |
| | 9 | PIC - UIC | 2.08 [0.03, 4.09] | 0.95 | Yes | Post |

**Supplementary Table S13: Viral-load burden (area under the viremia curve, AUC) analyses within the ZIKV-infected group (n = 12; 6 female, 6 male). (A) Sex difference in burden from log(AUC) ~ Sex. (B) Association of standardized log-AUC with each principal outcome, entered as a predictor in that outcome's own model family: standardized mean difference per SD for neuroimaging (ICV-adjusted), log-rate for the acute-stress and attachment behaviors, and latent-logit for the ordinal INAS domains (fit with a random**

**age slope per animal). Estimates are posterior medians with 89% highest-density intervals; pd is the probability of direction; a contrast is credible when the interval excludes zero. Every model was refit excluding one male with atypically low burden (more than two log10 units below any other animal) as a sensitivity check.**

| Outcome | Comparison | Estimate [89% HDI] | pd | Credible |
|---|---|---|---|---|
| A. Sex difference in viral burden — log(AUC) ~ Sex; estimate = male minus female on the natural-log scale (exp = fold-change) | | | | |
| log(AUC) | All ZIKV (6F/6M) | -0.927 [-2.508, 0.69] | 0.828 | No |
| log(AUC) | ZIKV [excl. outlier] (6F/5M) | -0.182 [-0.949, 0.707] | 0.652 | No |
| log(AUC) | All ZIKV, cohort-adjusted | -1.465 [-2.831, 0.116] | 0.934 | No |
| B. Association of standardized log-AUC with each principal outcome — per-SD slope in that outcome's model family (see caption) | | | | |
| Amygdala | Male ZIKV, ICV-adj | -0.031 [-0.672, 0.642] | 0.534 | No |
| Amygdala | Male ZIKV, ICV-adj [excl. outlier] | -0.339 [-1.477, 0.884] | 0.681 | No |
| Total CSF | Female ZIKV, ICV-adj | -0.015 [-1.389, 1.253] | 0.507 | No |
| Total CSF | Female ZIKV, ICV-adj [excl. outlier] | -0.031 [-0.876, 0.791] | 0.525 | No |
| Lateral ventricle | Female ZIKV, ICV-adj | 0.219 [-1.091, 1.507] | 0.609 | No |
| Lateral ventricle | Female ZIKV, ICV-adj [excl. outlier] | 0.181 [-0.621, 0.996] | 0.645 | No |
| Right temporal-limbic | Female ZIKV, ICV-adj | 0.605 [-0.683, 1.901] | 0.774 | No |
| Right temporal-limbic | Female ZIKV, ICV-adj [excl. outlier] | 0.509 [-0.211, 1.282] | 0.866 | No |
| Left temporal-limbic | Female ZIKV, ICV-adj | 0.566 [-0.785, 1.915] | 0.743 | No |
| Left temporal-limbic | Female ZIKV, ICV-adj [excl. outlier] | 0.467 [-0.356, 1.261] | 0.826 | No |
| Left temporal-visual | Male ZIKV, ICV-adj | -0.025 [-0.251, 0.209] | 0.589 | No |
| Left temporal-visual | Male ZIKV, ICV-adj [excl. outlier] | 0.228 [-0.394, 0.766] | 0.765 | No |
| Hostile (Alone) | ZIKV, both sexes | 0.074 [-0.782, 0.861] | 0.557 | No |
| Hostile (Alone) | ZIKV, both sexes [excl. outlier] | -0.343 [-1.12, 0.449] | 0.761 | No |
| Hostile (Profile) | ZIKV, both sexes | -0.506 [-1.068, 0.03] | 0.952 | No |
| Hostile (Profile) | ZIKV, both sexes [excl. outlier] | -0.487 [-0.905, -0.052] | 0.958 | Yes |
| Threat barks | Female ZIKV, condition-adj | -0.346 [-1.516, 0.734] | 0.694 | No |
| Threat barks | Female ZIKV, condition-adj [excl. outlier] | -0.231 [-0.966, 0.431] | 0.721 | No |
| Scream (attachment) | Male ZIKV | 1.179 [-0.009, 2.245] | 0.935 | No |
| Scream (attachment) | Male ZIKV [excl. outlier] | -0.021 [-1.081, 1.036] | 0.513 | No |
| State control (INAS) | ZIKV, age-adj | 0.135 [-0.498, 0.768] | 0.634 | No |
| State control (INAS) | ZIKV, age-adj [excl. outlier] | -0.503 [-1.112, 0.08] | 0.916 | No |
| State control (INAS) x age | ZIKV | 0.169 [0.07, 0.273] | 0.993 | Yes |
| State control (INAS) x age | ZIKV [excl. outlier] | 0.051 [-0.064, 0.161] | 0.775 | No |
| Neuromotor (INAS) | ZIKV, age-adj | -0.012 [-0.199, 0.185] | 0.538 | No |
| Neuromotor (INAS) | ZIKV, age-adj [excl. outlier] | -0.046 [-0.269, 0.174] | 0.642 | No |
| Neuromotor (INAS) x age | ZIKV | -0.049 [-0.108, 0.016] | 0.896 | No |
| Neuromotor (INAS) x age | ZIKV [excl. outlier] | -0.021 [-0.093, 0.05] | 0.69 | No |
| Orientation (INAS) | ZIKV, age-adj | -0.285 [-0.785, 0.203] | 0.841 | No |
| Orientation (INAS) | ZIKV, age-adj [excl. outlier] | 0.125 [-0.324, 0.575] | 0.686 | No |

| | | | | |
|---|---|---|---|---|
| **Orientation (INAS) x age** | **ZIKV** | **0.166 [0.021, 0.315]** | **0.965** | **Yes** |
| **Orientation (INAS) x age** | **ZIKV [excl. outlier]** | **0.069 [-0.058, 0.209]** | **0.804** | **No** |

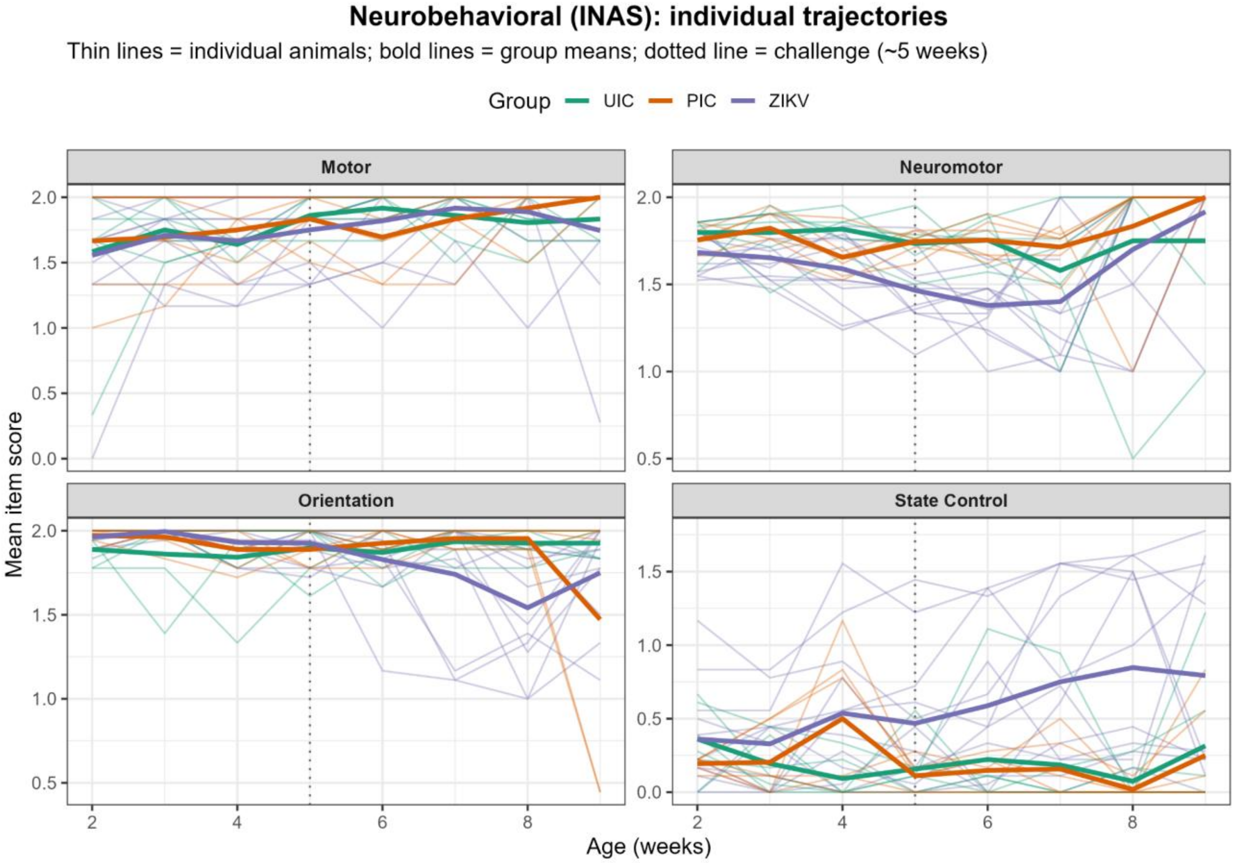


***Supplementary Figure S2. Individual neurobehavioral (INAS) trajectories.*** *Each thin line shows one animal's mean item score for the indicated domain across age (weeks); bold lines are group means, colored by treatment group (UIC, PIC, ZIKV). The vertical dotted line marks the timing of ZIKV challenge and Poly-IC administration (approximately 5 weeks of age). Panels show the Orientation, Neuromotor, Motor, and State Control domains.*

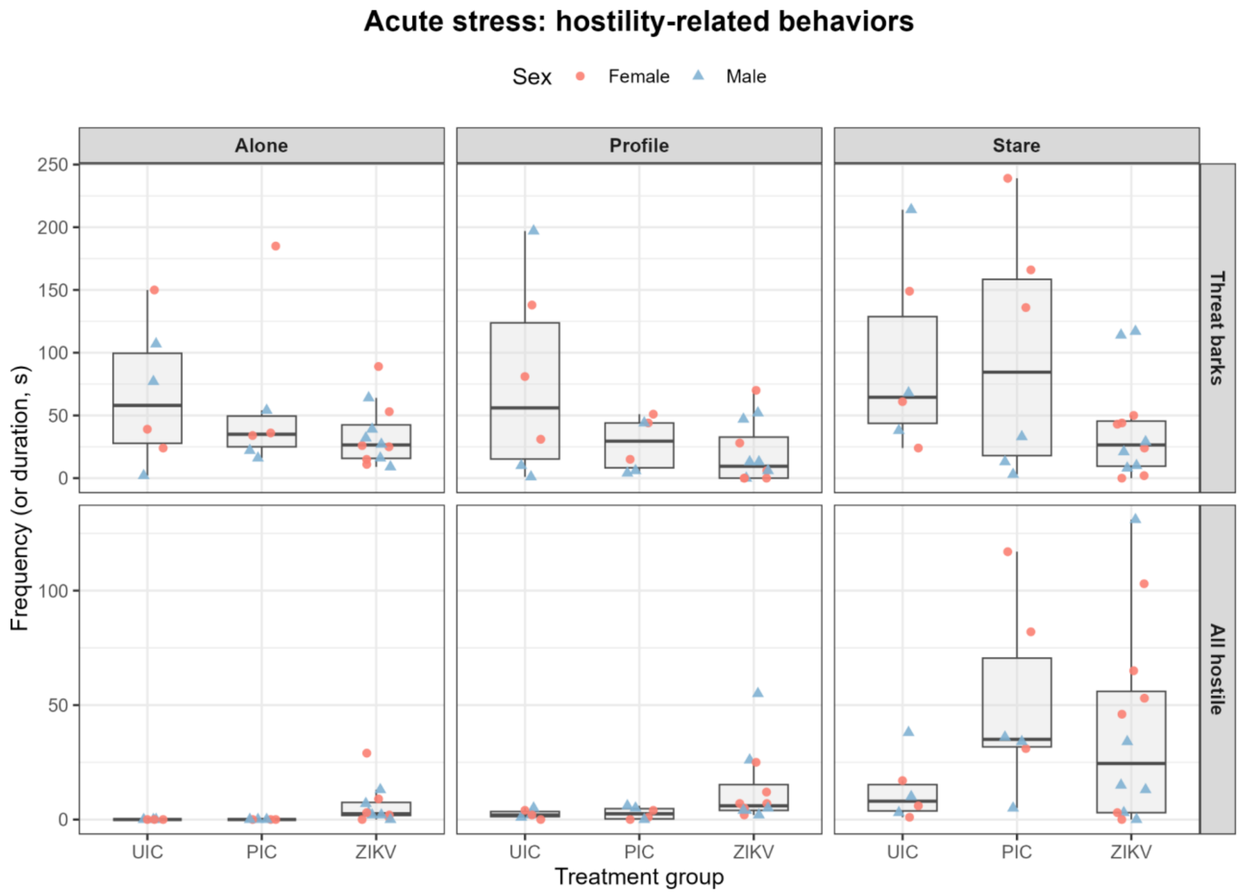


***Supplementary Figure S3a. Acute-stress (Human Intruder) data, hostility-related behaviors (threat barks; all hostile behavior).*** *Box-and-whisker plots (box = interquartile range, line = median, whiskers = range) with each individual animal shown as a point colored and shaped by sex, by treatment group (UIC, PIC, ZIKV) and condition (Alone, Profile, Stare).*

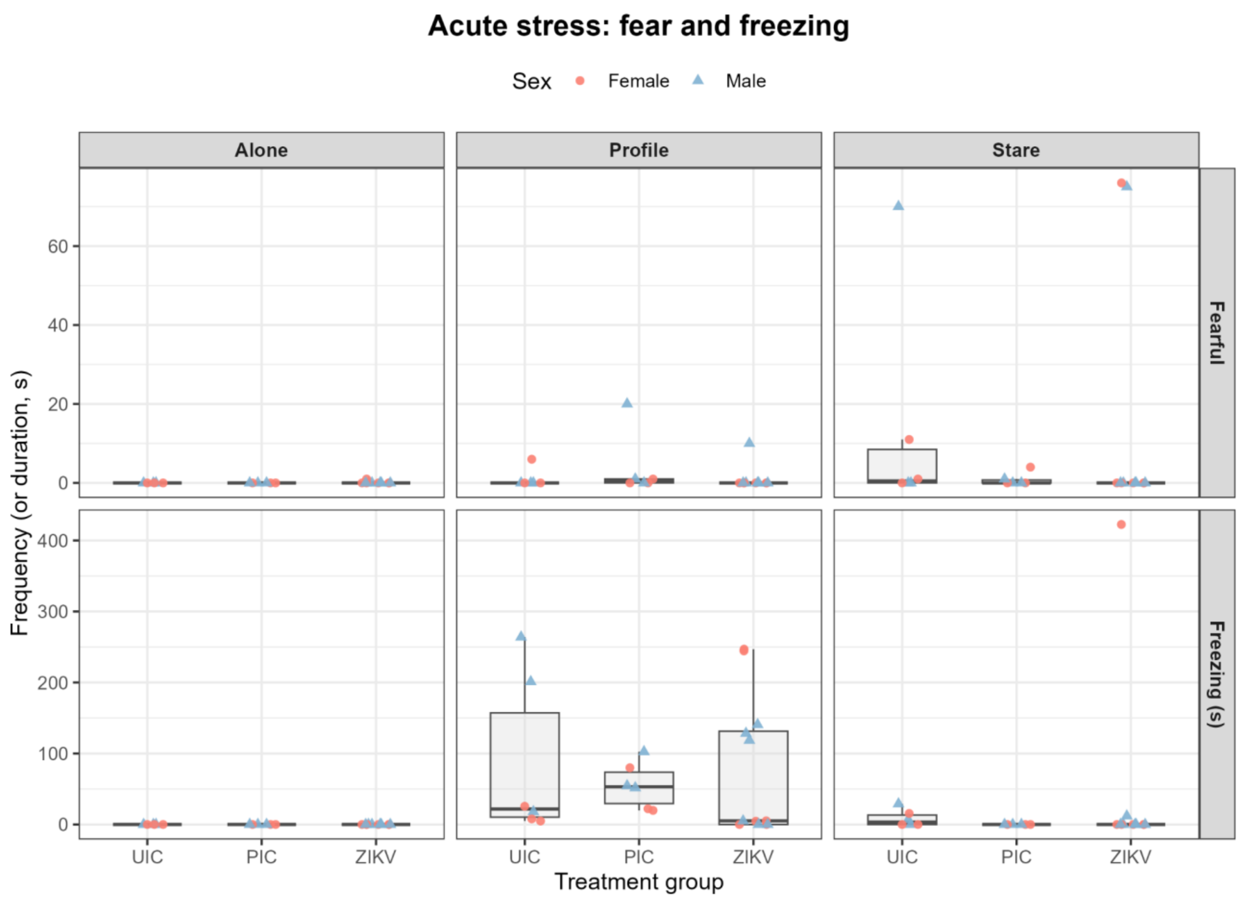


***Supplementary Figure S3b. Acute-stress (Human Intruder) data, fear and freezing (fearful behavior; freezing, seconds).*** *Plotting conventions as in Supplementary Figure S3a.*

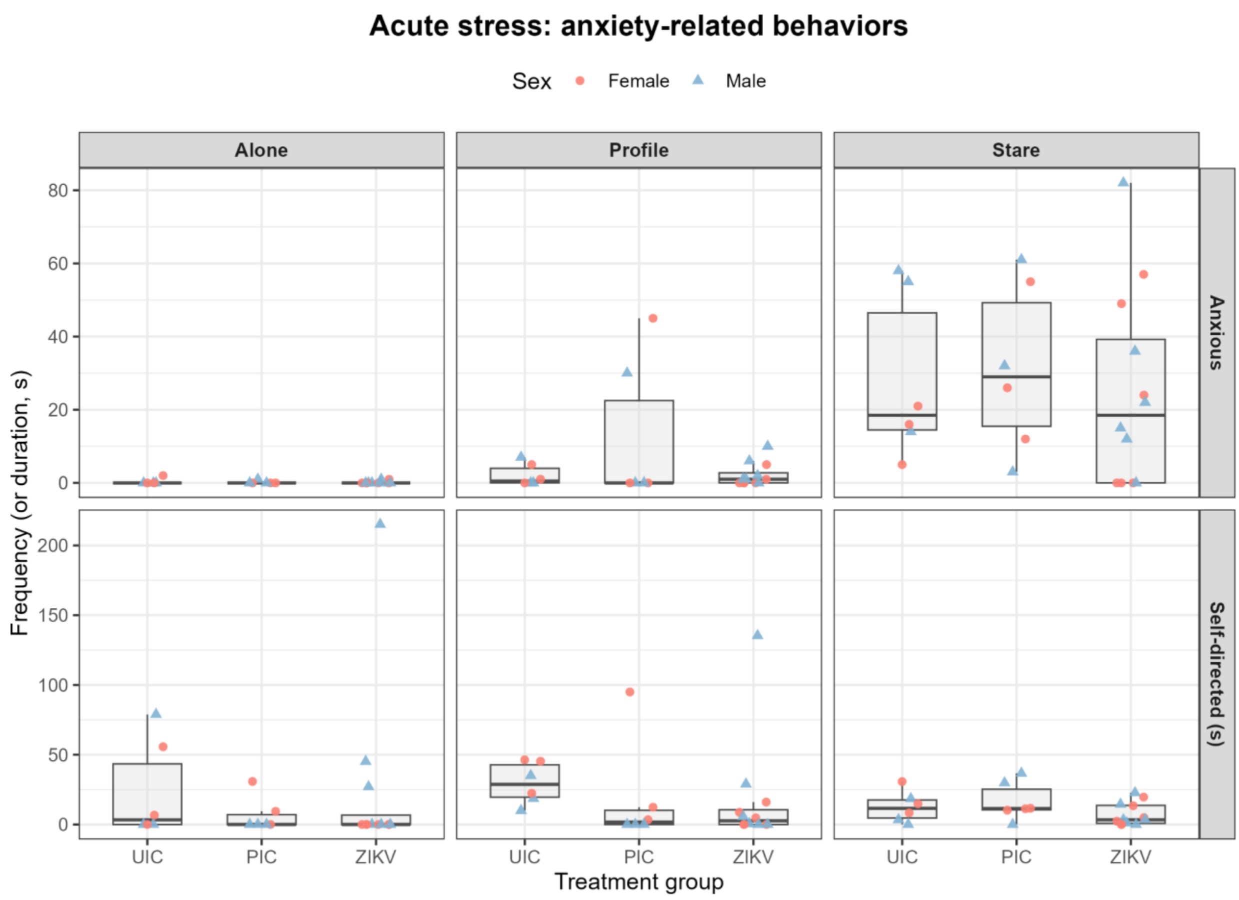


***Supplementary Figure S3c. Acute-stress (Human Intruder) data, anxiety-related behaviors (anxious behavior; self-directed behavior).*** *Plotting conventions as in Supplementary Figure S3a.*

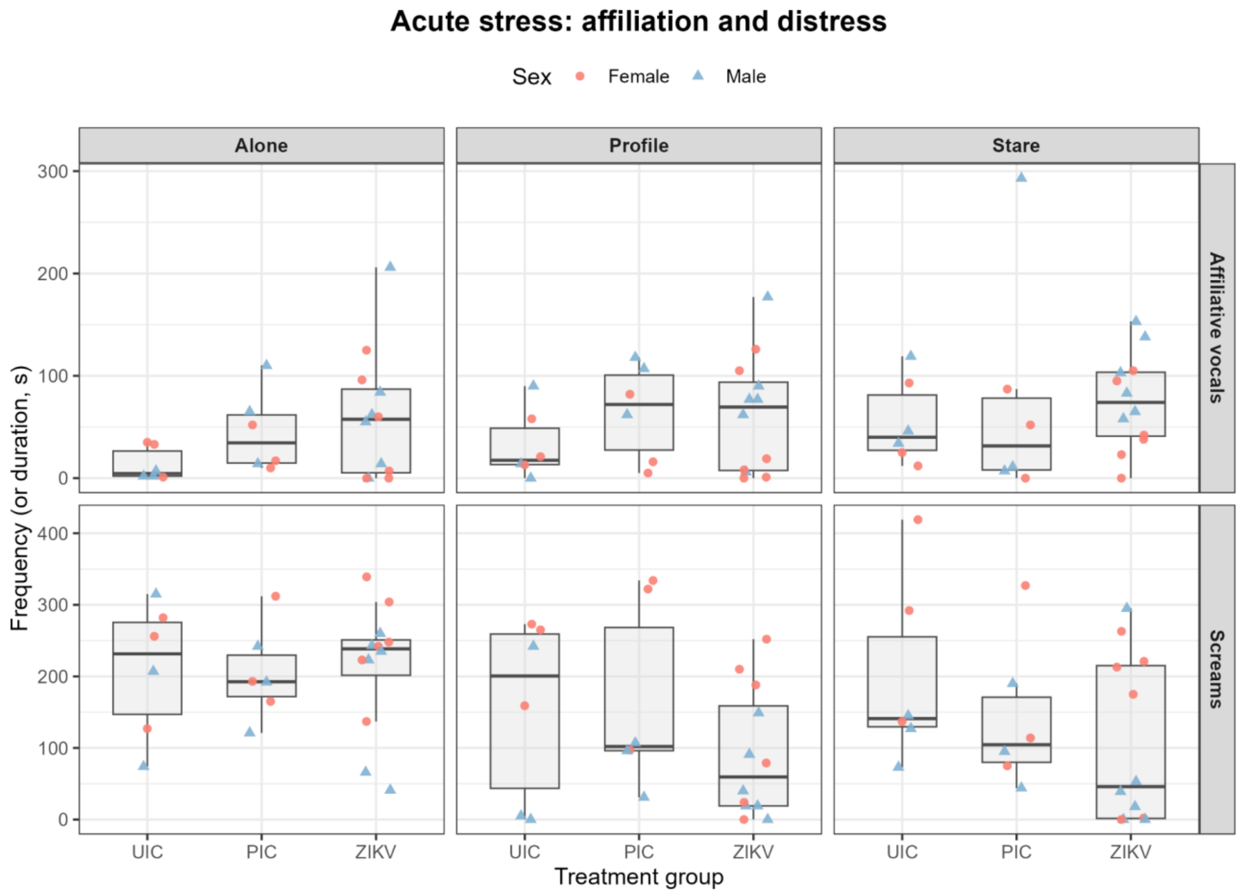


***Supplementary Figure S3d. Acute-stress (Human Intruder) data, affiliation and distress vocalizations (affiliative vocalizations; screams). Plotting conventions as in Supplementary Figure S3a.***

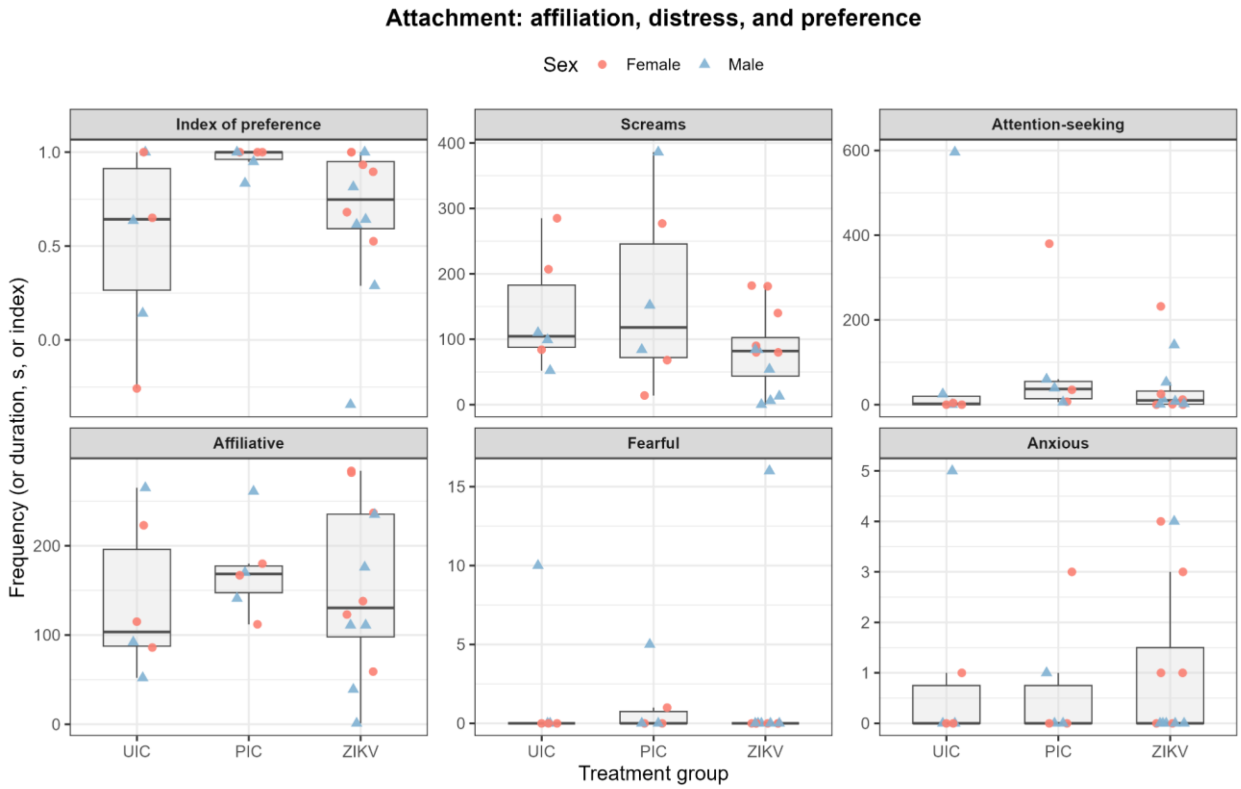


***Supplementary Figure S4a. Attachment-assessment data, affiliation, distress, and preference (index of preference; screams; attention-seeking; affiliative; fearful; anxious behaviors).*** *Box-and-whisker plots (box = interquartile range, line = median, whiskers = range) with each individual animal shown as a point colored and shaped by sex, by treatment group (UIC, PIC, ZIKV).*

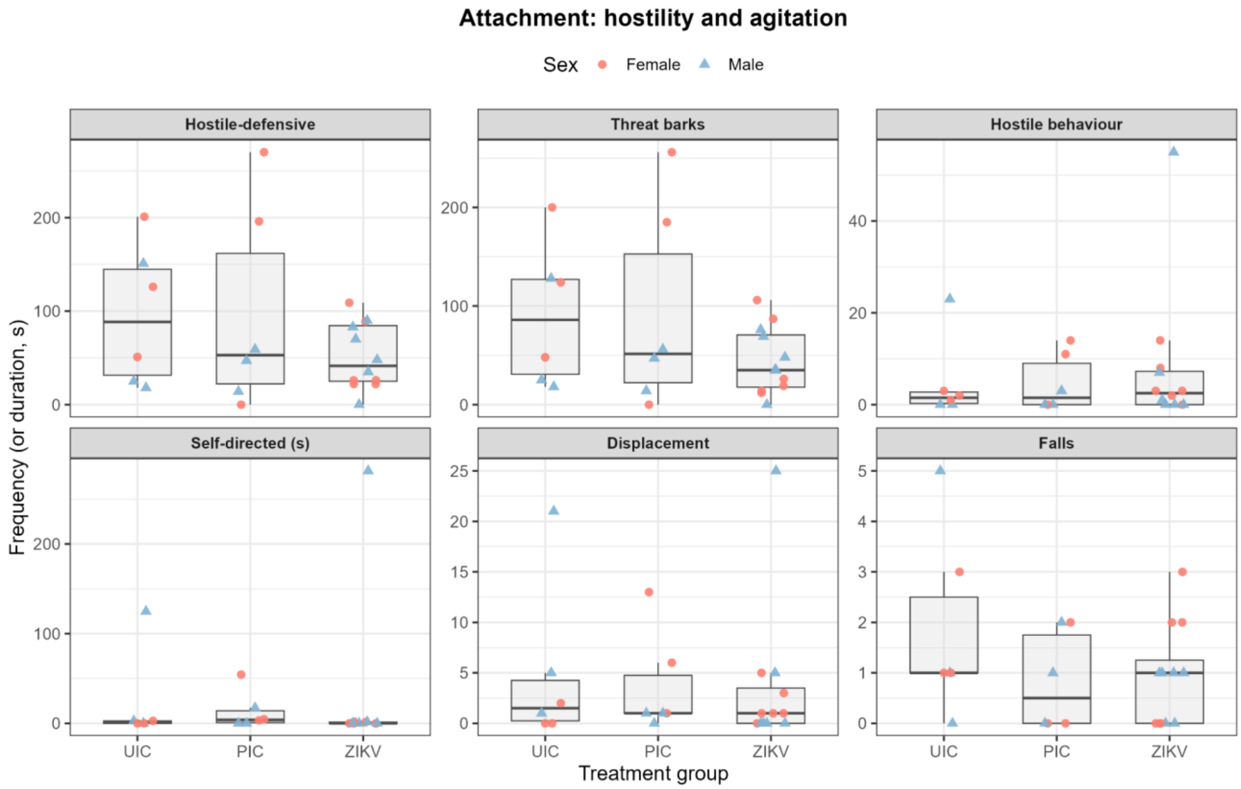


***Supplementary Figure S4b. Attachment-assessment data, hostility and agitation (hostile-defensive; threat barks; hostile behavior; self-directed behavior; displacement; falls).*** *Plotting conventions as in Supplementary Figure S4a.*

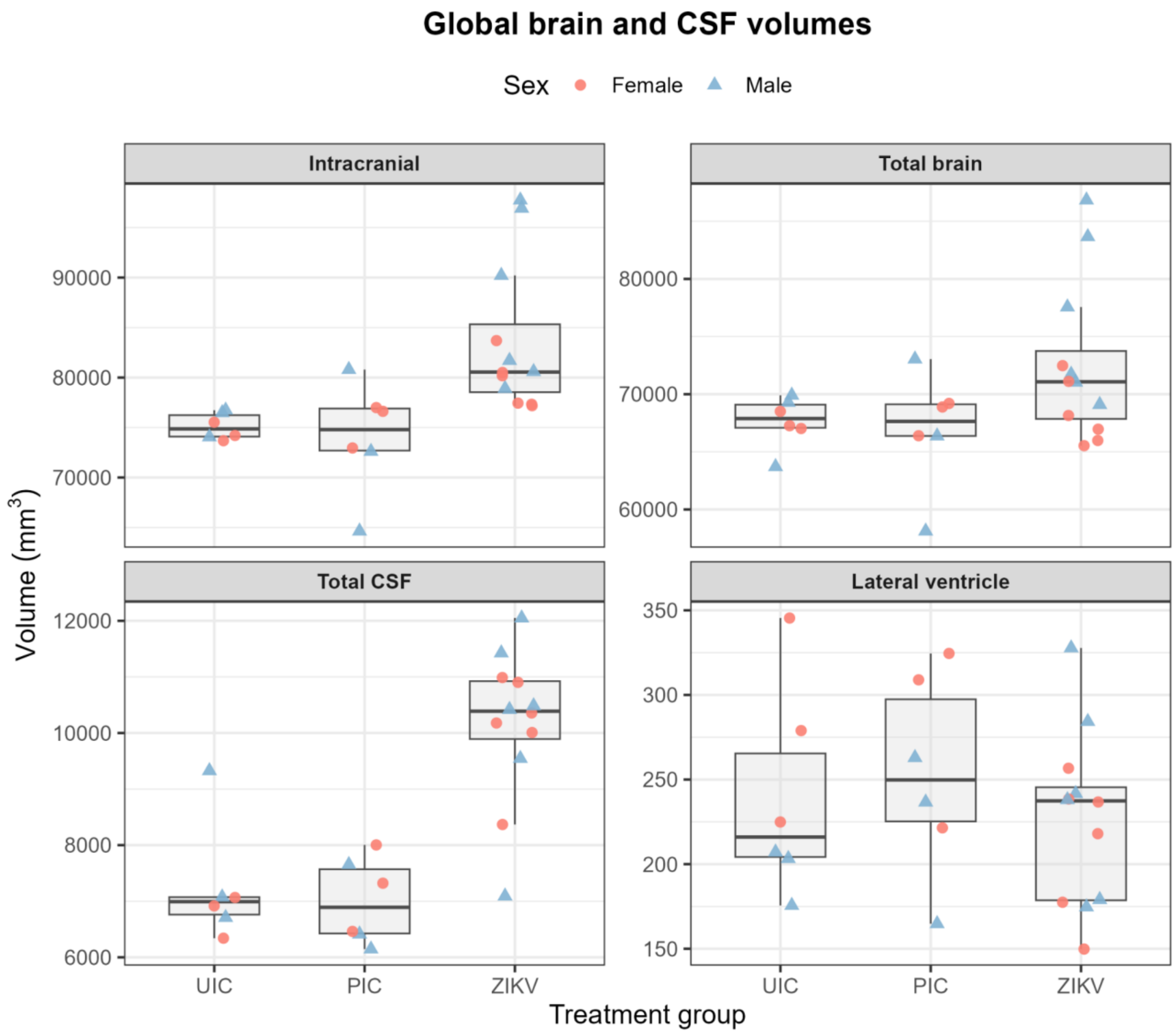


***Supplementary Figure S5a. Global brain and CSF volumes (total intracranial volume; total brain volume; total CSF volume; lateral ventricle).*** *Box-and-whisker plots (box = interquartile range, line = median, whiskers = range) of the raw volume (mm3), with each individual animal shown as a point colored and shaped by sex, by treatment group (UIC, PIC, ZIKV).*

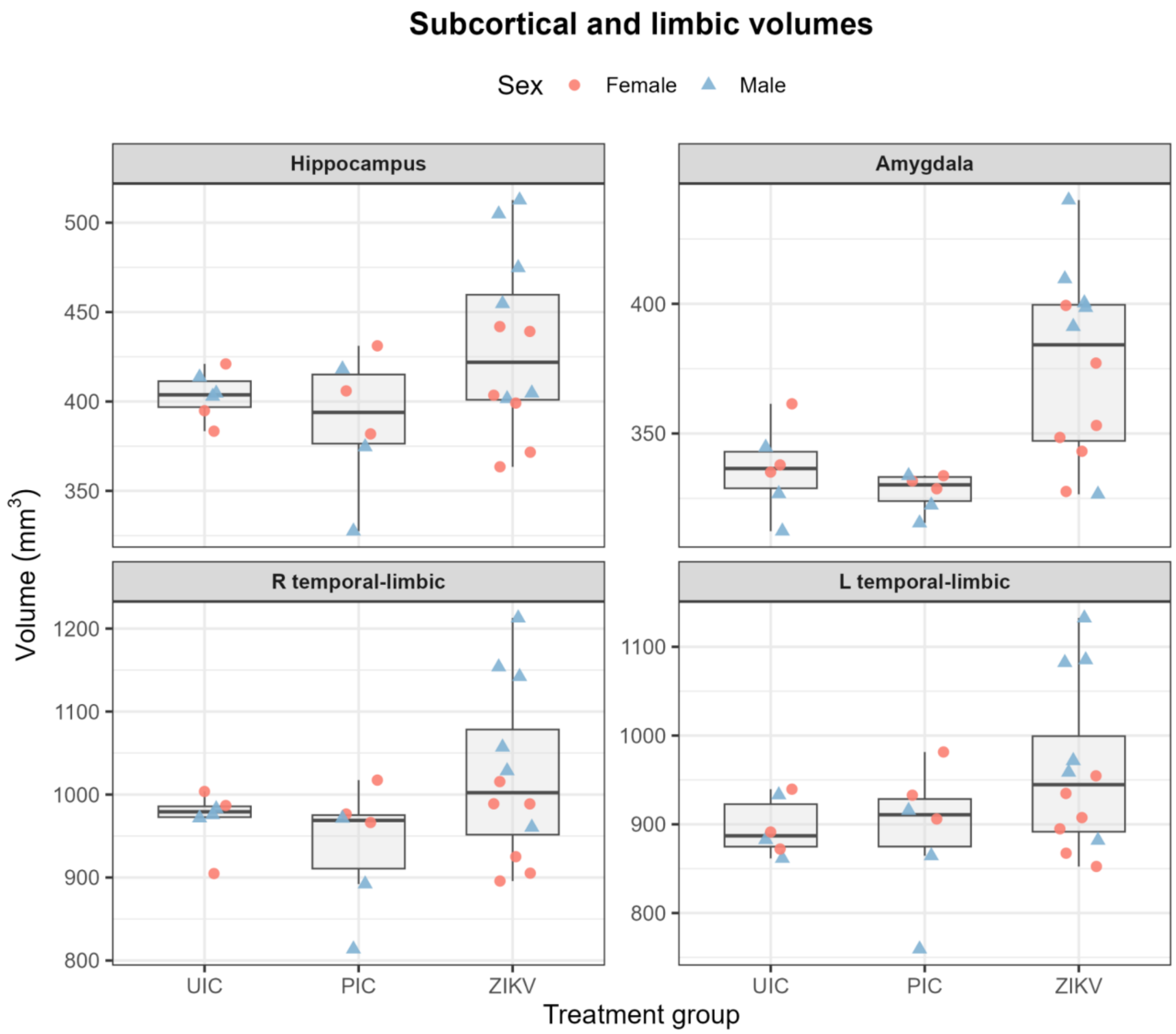


***Supplementary Figure S5b. Subcortical and limbic volumes (hippocampus; amygdala; right and left temporal-limbic).*** *Plotting conventions as in Supplementary Figure S5a.*

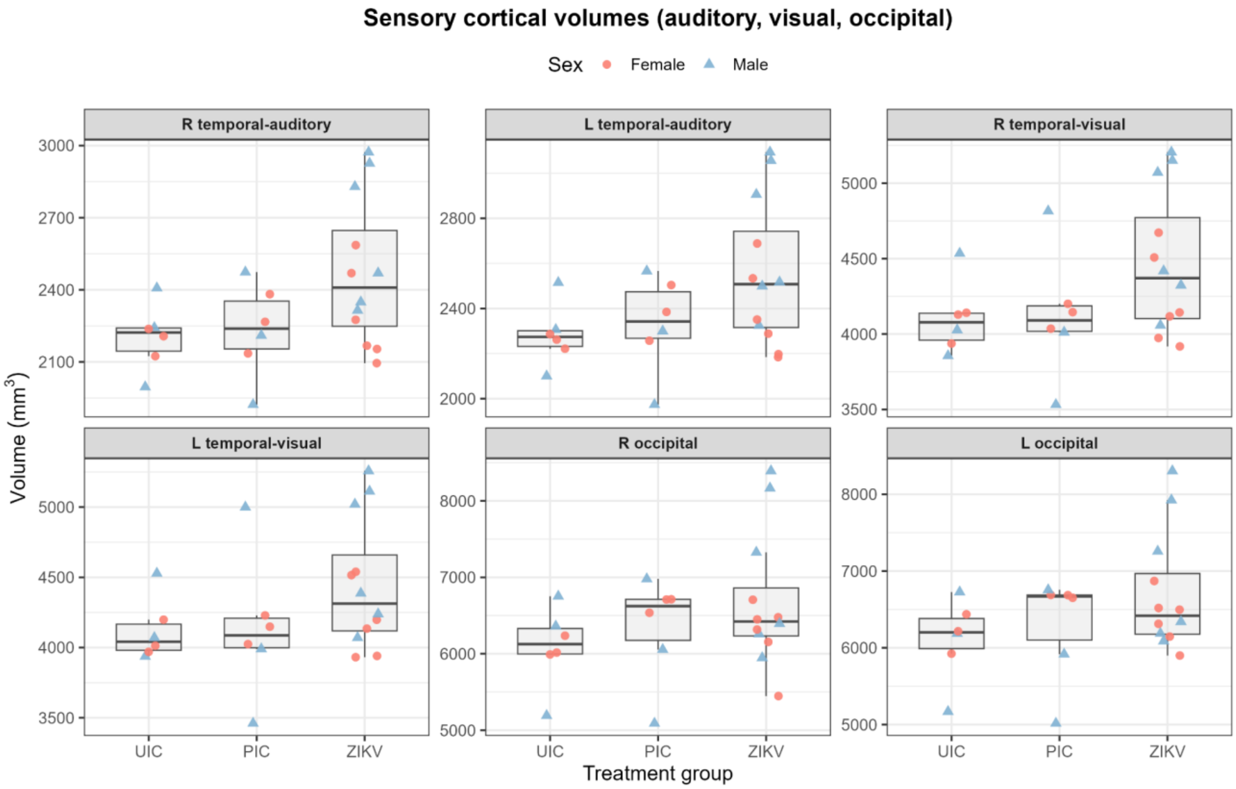


***Supplementary Figure S5c. Sensory cortical volumes (right and left temporal-auditory, temporal-visual, and occipital).*** *Plotting conventions as in Supplementary Figure S5a.*

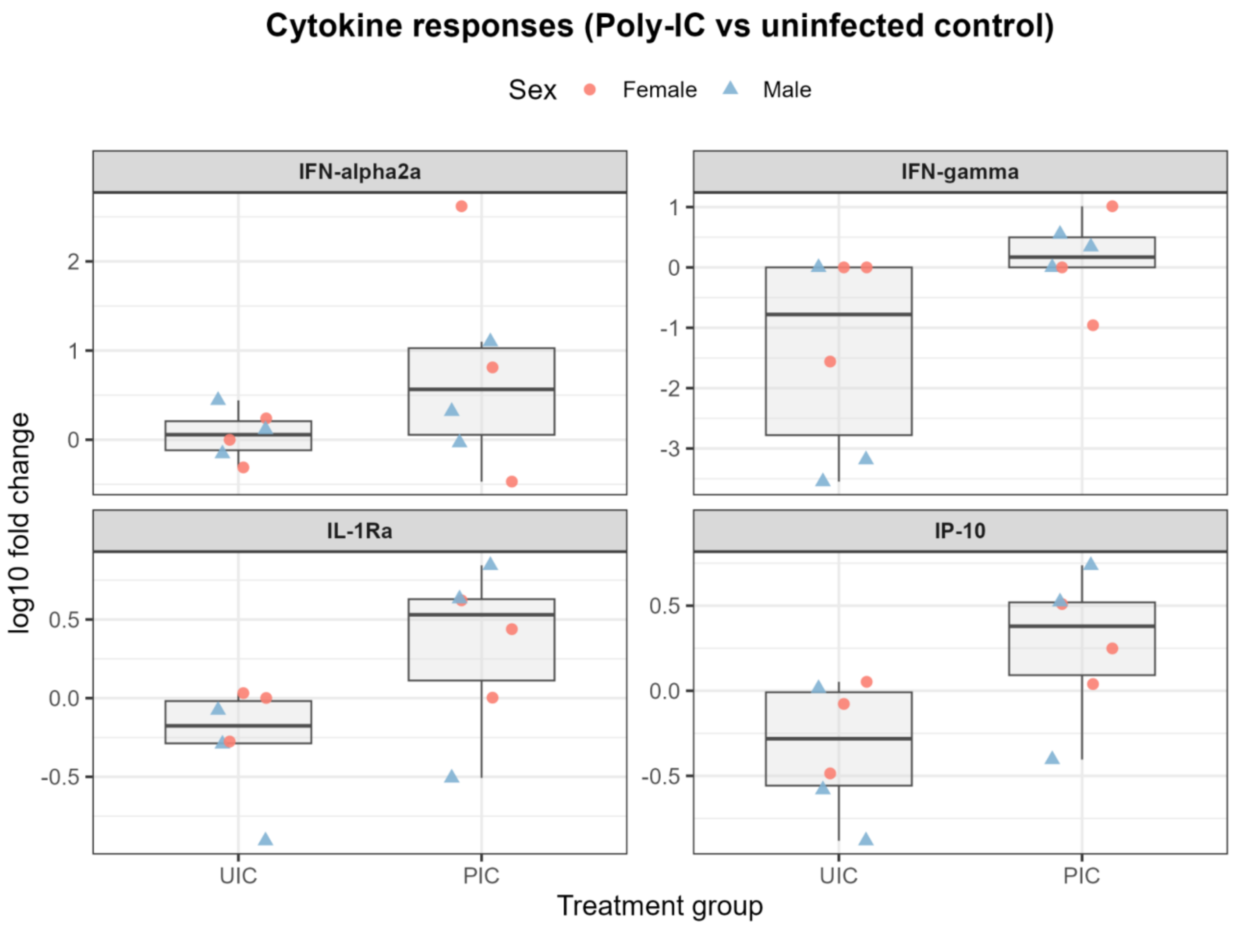


***Supplementary Figure S6. Individual cytokine responses.*** *Box-and-whisker plots (box = interquartile range, line = median, whiskers = range) of the log10 fold change for each analyte (IFN-alpha2a, IFN-gamma, IL-1Ra, IP-10) by control group, with each individual animal shown as a point colored and shaped by sex; cytokines were not measured in the ZIKV group owing to limited blood volume.*

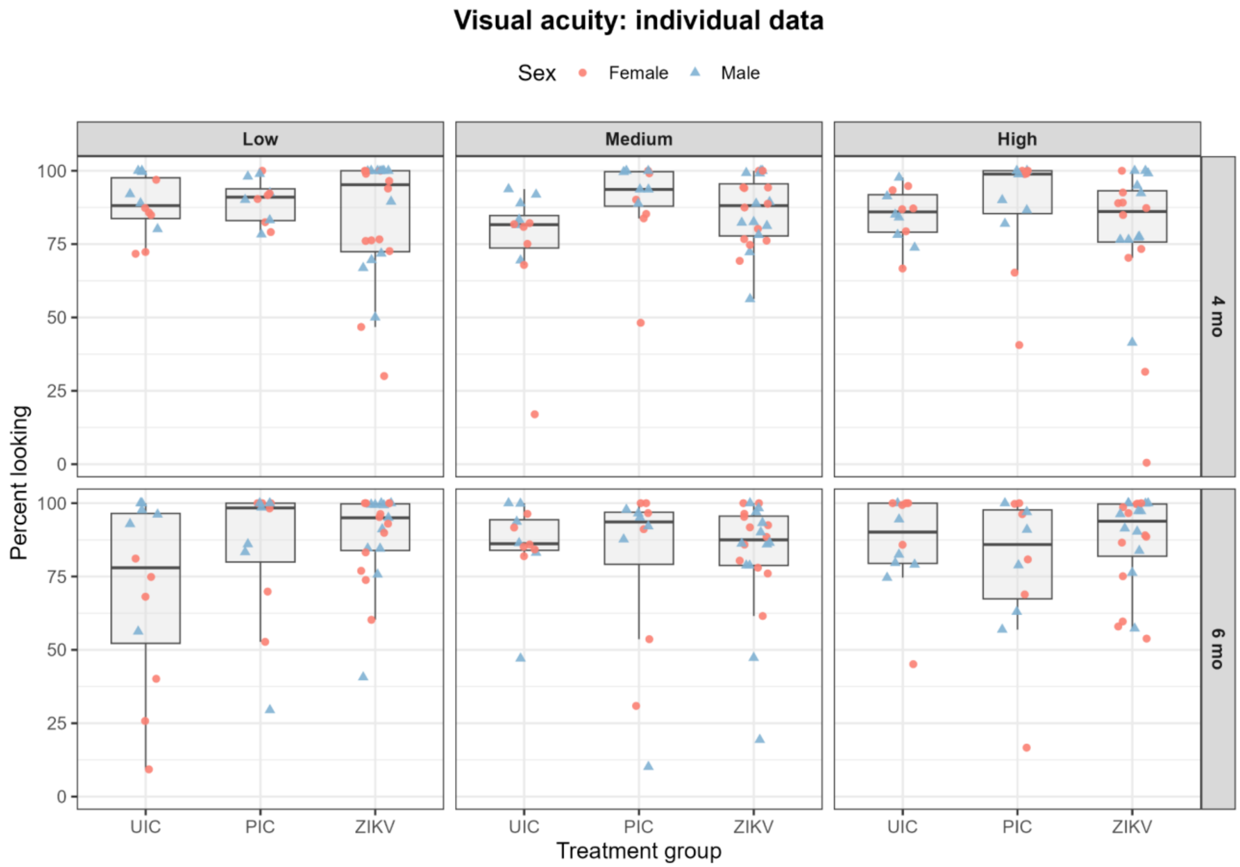


***Supplementary Figure S7. Individual visual-acuity data.*** *Box-and-whisker plots (box = interquartile range, line = median, whiskers = range) of percent looking by treatment group (UIC, PIC, ZIKV), age (4 and 6 months), and contrast level (low, medium, high), with each individual animal shown as a point colored and shaped by sex.*
